# Transcriptomic-Based Identification of FCRL6 as a Novel Diagnostic Biomarker in Diabetic Nephropathy

**DOI:** 10.1101/2024.11.01.621451

**Authors:** Adarsh Marinedrive, Sonalika Ray, Mohit Mazumder

## Abstract

Diabetic nephropathy (DN) is a microvascular complication that arises in a sizable portion of individuals with diabetes mellitus, representing a growing public health issue given its progressive injury of the kidney. DN is primarily driven by the upregulation of pro-inflammatory markers and pathways, which cause hyperfiltration and significant physiological changes to the renal structure. To further understand the molecular mechanisms by which DN progresses, we used bioinformatics and meta-analysis techniques to analyze two datasets from the Gene Expression Omnibus (GEO) database: GSE142025 and GSE142153. Differential gene expressions (DGEs) were identified using the Bioconductor DESeq2 library and further used in functional enrichment analysis. DGEs were also constructed into protein-protein interaction networks and used to find regulatory hub genes. In total, 911 DGEs were found, and enrichment using the Kyoto Encyclopedia of Genes and Genomes (KEGG) and Gene Ontology (GO) revealed strong correlations with immunoreactive responses and cascades. In our analysis, we discovered 12 protein-coding genes novel with regards to DN; we further identified Fc Receptor-Like 6 - FCRL6 - as a potential biomarker in nephropathy and described its involvement in the suppression of natural killer cell-mediated toxicity as a mechanism of poor immunoregulation in the diabetic kidney. The current study presents comparative analyses of the DN transcriptome and a possible diagnostic marker with applications in immunotherapy treatments of nephropathy.

## 1. Introduction

Diabetes mellitus (DM) is a highly prevalent metabolic condition in which the body is unable to effectively utilize and manage blood glucose [1]. Typically, individuals with DM suffer from persistent insulin resistance, leading to hyperglycemia and dysregulation of protein and lipid synthesis. The malignant disorder has secondary impacts on various systems; most periphery blood vessels are impaired by excess glycemia, developing atherosclerosis [2]. The hardening, thickening, and tightening of endothelial walls means patients with DM are at significantly higher risk of developing macrovascular complications due to a sustained limitation of oxygen flow (hypoxia) [3][4]. Thus is the case that coronary artery disease, peripheral artery disease, stroke, myocardial infarction, and heart failure are more likely to appear in individuals with diabetes than those without [5].

Diabetic nephropathy (DN) is a major microvascular complication - along with retinopathy and neuropathy - that arises in up to half of diabetic patients [6]. The condition sees a progressive loss of renal functioning with increased inflammation and hypertension limiting successful glomerular filtration. Around a third of DN patients progress to end-stage renal disease (ESRD), or kidney failure [7]. While all renal cells are exposed to an increased glucose concentration, only select ones exhibit significant changes to genetic and chemical regulation [8]. Progression of atherosclerosis in interlobar vessels and interlobular capillaries, for example, can be correlated to heightened glucose sensitivity in the present endothelial linings [9]. As such, the effects of diabetes on the kidneys are primarily due to hypoxia and pro-inflammatory upregulations [10][11]. Nephropathic influence on renal physiology is first seen through the thickening of the glomerular basement membrane, which is accompanied with changes to visceral podocytes [12]. A reduction in the filtration capacity of the glomerulus is noted in the secondary stage of DN progression. Glucose reuptake is severely upregulated in the proximal tubule, increasing the oxidative stress placed on various renal segments [13]. In addition, hyperabsorption of NaCl in the proximal tubule and loop of Henle structures in the nephron creates a sodium imbalance in the distal convoluted tubule region, preventing effective homeostatic regulation of solute filtration.

Later stages of DN are typically characterized by extreme hyperfiltration in the glomerulus, along with the widespread presence of inflammatory factors [14]. Changes to glycemic content in blood plasma leads to the upregulation of the renin-angiotensin-aldosterone system (RAAS), bolstering intracellular angiotensin II concentration [15][16]. Similarly, the metabolic activity of the Janus kinase/signal transducers and activators of transcription (JAK/STAT) pathway is increased in advanced stages of DN [17]. Heightening of the NF-κB system is seen most in glomeruli and intratubular regions [18]. The upregulation of these pathways, along with hyperglycemia, hypoxia, and the production of advanced glycation end products (AGEs), increases the production of pro-inflammatory molecules [19][20]; cytokines, members of the interleukin (IL) family, chemokines, vascular endothelial growth factor, and tumor necrosis factors α and I recruit monocytes and macrophages to the interstitium, mesangium, and tubules (**Figure 1**) [21][22][23]. The resulting inflammation and fibrogenesis often causes advanced kidney injury in the form of glomerulosclerosis and the formation of lesions in the capillary loops [24].

**Figure 1:**
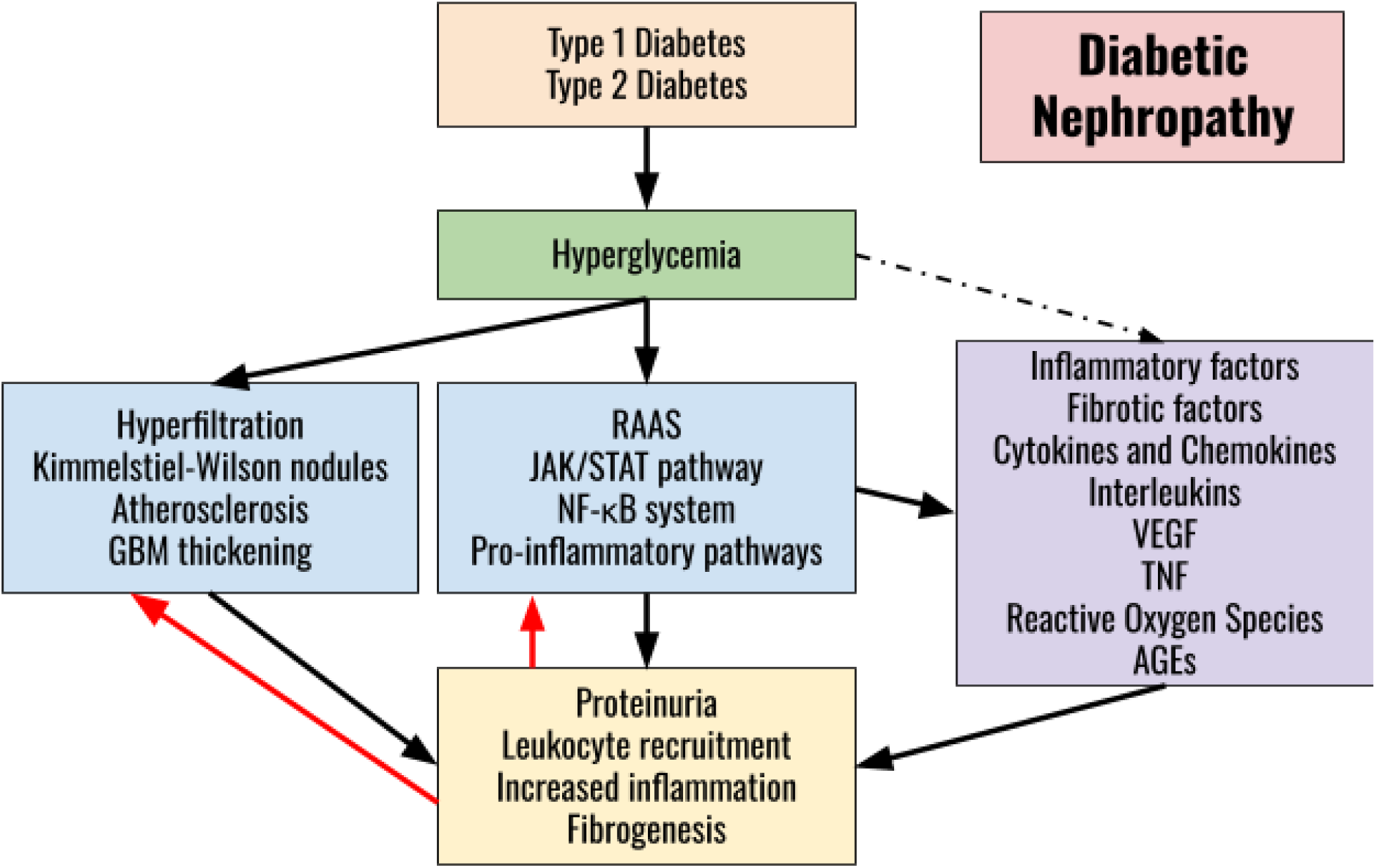
Overview of pathogenesis and stages of progression in diabetic nephropathy. Hyperglycemia induces physiological changes to the renal structure and the upregulation of inflammatory pathways; factors combine with products of these pathways to induce the pathological symptoms observed in DN. Leukocyte activation feeds back into the development of injury.

Excessive proteinuria is a clear indicator of DN progression [25]; Intracellular reactive oxygen species (ROS) - caused by glucose auto-oxidation and the formation of AGEs - combine with tubular inflammation to force hyperfiltration in glomeruli [26]. The presence and severity of DN are diagnosed through measurement of the urinary albumin-to-creatinine ratio (uACR) in conjunction with the estimated glomerular filtration rate (GFR) [27][28]; in many advanced cases of nephropathy, a ten-fold or higher change can be observed in urinary albumin (**Table 1**). Such characterization of proteinuria has been used to distinguish early DN (EDN) from advanced DN (ADN). In addition, serum uric acid and other biomarkers have been put forward as potential indicators of kidney dysfunction [29].

**Table 1:** National Kidney Foundation categorization of diabetic nephropathy by estimated glomerular filtration rate and urinary albumin-to-creatinine ratio and risk of ESRD. Green: No to low risk; Yellow: Moderately increased risk; Orange: Heavily increased risk; Red: Severely increased risk or identifiable kidney failure.

| eGFR Categories<br>(mL/min/1.73m<sup>2</sup>) | Albuminuria Categories<br>Urinary Albumin-to-Creatinine Ratio |  |  |
| --- | --- | --- | --- |
|  | A1 (<30 mg/g) | A2 (30-300 mg/mmol) | A3 (>300 mg/mmol) |
| G1 (≥ 90) | Normal | A2 G1 | A3 G1 |
| G2 (60-89) | Normal | A2 G2 | A3 G2 |
| G3a (45-59) | A1 G3a | A2 G3a | A3 G3a |
| G3b (30-44) | A1 G3b | A2 G3b | A3 G3b |
| G4 (15-29) | A1 G4 | A2 G4 | A3 G4 |
| G5 (<15) | A1 G5 | Kidney Failure | Kidney Failure |

Current treatments of DN are mainly restricted to generic amelioration of hyperglycemia. Measures to control blood pressure, cholesterol, and glucose content have shown the most positive results in reducing the progression of nephropathy [30]. Through either dietary restrictions or medical intervention, most patients are able to limit kidney injury, and in some cases, show signs of improvements in filtration output. Individuals with advanced nephropathy or ESRD are oftentimes forced to undergo regular hemodialysis to remove excess fluid and soluble waste [31]. Nephrologists may prescribe aldosterone inhibitors, angiotensin-converting enzyme inhibitors, or angiotensin II receptor blockers to counter the upregulation of RAAS and RAAS-induced pathways [32]. Other anti-inflammatory gene promoters have been tested, as have the converse inhibitors, with attempts to suppress cytokine and chemokine-inducing pathways. Downregulating the production of transforming growth factor beta (TGF-I) has been put forward as a potential mechanism in slowing the development of kidney injury [33][34]. An alternative method of treatment under increased testing, however, is gene therapy, which has shown mixed to positive results in amelioration [35].

Additional research is required to fully understand the physiological and metabolic mechanisms by which DN takes shape; identification of biomarkers paves the way for more potential therapeutic treatments. Controlling the transcriptome with regards to certain pathways can only be done with more widespread knowledge of gene interactions in DN. Transcriptomic analysis of kidney RNA-seq samples allows for a cross-sectional comparison of gene expression between normal and DN patients, increasing the potential of identifying novel gene targets. Previous bioinformatics-based approaches have found TGFB1, TNF, RELA, AGT, NOS2, IL6, IL1B, FOS, HMOX, VEGFA, CCL2, AKT1, SERPINE1, CAT, and various other inflammatory response-related genes as highly correlated with nephropathy progression and could act as transcriptomic regulatory drug targets [36][37][38]. Similar meta-analyses identify signal transduction, metabolism of proteins, immune system, AGE/RAGE signaling pathway in diabetic complications, and homeostasis as primary pathways implicated in renal injury in DN [39]. Here, we analyzed transcriptomic data from the next-generation sequencing dataset GSE142025 and microarray dataset GSE142153 from the National Center for Biotechnology Information (NCBI)’s Gene Expression Omnibus (GEO) to identify developmental factors and differential gene expressions (DGEs) in diabetic nephropathy [40][41]. Principal component analysis was performed to group samples based on diagnosis. Pathway and network analysis followed through various gene set enrichment tools in order to map out larger connections in the selected transcriptome and identify functionalities of interest genes. Ontologies were likewise identified to mark relations between DGEs. In sum, we present our findings parallel to literature along with a novel gene target to further evaluate in DN treatment. Through this exploration, we hope to further our understanding of the transcriptomic landscape of the diabetic kidney and its implications in the pathogenesis of DN.

## 2. Methodology

### 2.1. Data Extraction and Cleaning

In this study, we obtained and analyzed two publicly available datasets from the GEO, GSE142025 and GSE142153, whose samples are here referred to as Cohort A and Cohort B, respectively (**Table 2**). In the prior, 27 DN samples categorized into EDN (n = 6) and ADN (n = 21) were compared to expression data from control samples (n = 9). The latter dataset consists of samples from DN patients (n = 23), ESRD patients (n = 7), and healthy controls (n = 10). Only data on DN was considered and ESRD samples were used solely for principal component analysis. For each dataset, a threshold filter of greater than 5 was placed on the cumulative read count of each gene to eliminate those with low expression. In addition to the gene expression data, the series matrix files were extracted to facilitate downstream analysis.

**Table 2:** Pivot table displaying the summary sample statistics of Cohort A and Cohort B.

| Sample Type | GSE142025:<br>Cohort A | GSE142153:<br>Cohort B |
| --- | --- | --- |
| DN | 6 (EDN)<br>21 (ADN) | 23 |
| ESRD | -/- | 7 |
| Control | 9 | 10 |
| Total | 36 | 40 |

### 2.2. Principal Component Analysis

Exploratory data analysis via principal component analysis (PCA) was performed on the quantile normalized values from both sample sets to better interpret characteristics and distribution; the FactoMineR package was used in dimensionality reduction, visualizing data similarities along linear planes and cutting out noise [42]. Classification of samples was based on series matrix designations from the original study and a typical *k*-means machine learning algorithm was not utilized. The groupings used in the PCA for Cohort A were “EDN,” “ADN,” and “Control.” Cohort B was divided into “DN,” “ESRD,” and “Control.”

### 2.3. Differential Gene Expression Analysis

Differential gene expression (DGE) analysis was carried out across both datasets using the DESeq2 package in Bioconductor [43]. The DESeq2 package uses count data to estimate mean-variance relationships and identify differentially expressed genes (DEGs) based on a negative binomial distribution model [44]. DESeq2 objects were generated to provide statistical results for each gene’s expression, outputting mean *T*-values, *P*-values, and false discovery rate (FDR)-adjusted *Q*-values using the Benjamini-Hochberg correction method [45]. Log_2_ fold changes were calculated, allowing for robust interpretation of gene expression variation. DEGs were considered significant if they showed a log_2_ fold change (log_2_FC) of ±1.5 for Cohort A and

±1 for Cohort B, with statistical significance set at a *Q*-value threshold of α < 0.05. A volcano plot was used to visualize genes with significant up- or downregulation across the datasets.

### 2.4. PPI Network Construction and Hub Gene Identification

A protein-protein interaction (PPI) network was constructed to visualize predicted interactions between polypeptide products of differentially expressed genes in DN-positive individuals; three networks were formulated based on identified DGEs in EDN and ADN groups of Cohort A in addition to the DN group of Cohort B using the STRING database [46]. Proteins of interest were marked as nodes, while interactivity was shown through edges, or the links. Evidence of interaction was based on text mining, experimentation, databases, co-expression, gene neighborhoods, gene fusion, and co-occurrences, while a minimum threshold interaction score of 0.900 was set to increase validational strength of the connections. A *k*-means clustering algorithm was used in an effort to minimize the interaction distance between proteins and cluster centroids, revealing functional linkages between closer proteins. In turn, larger themes of interaction were discovered. The NetworkAnalzyer app was utilized in Cytoscape 3.10.2 to understand interactive characteristics, including degrees of nodes, neighborhood connectivity, and centrality [47][48]. The results of the topological landscape of the networks created are not described here. Cytoscape also generated the visual representations of the PPI networks. Top hub genes - that is, transcripts that produce highly connected proteins in their respective PPI networks - were identified for their potential importance in regulating the web of proteomic connections. Here, the cytoHubba plug-in employed the Maximal Clique Centrality (MCC) algorithm to display the 15 most significant regulatory nodes in each network [49]. Hub genes were then present in further analyses of differential regulation of the transcriptome in nephropathy. Hub gene nodes were colored based on the strength of their clustering coefficient.

### 2.5. Gene Ontology and Pathway Analysis

Gene set enrichment analysis was performed on nephropathic samples to annotate larger trends in differential expression; functional enrichment was performed on the identified differentially expressed genes. Specific comparisons were made using both the Gene Ontology (GO) and Kyoto Encyclopedia of Genes and Genomes (KEGG) databases [50][51]. GO is a major push in the bioinformatics field to unify functions of gene products and model representations of interactions. Analysis of DGEs through the ToppGene suite and validation using the clusterProfiler package in R revealed ontologies in biological processes (BP), cellular components (CC), and molecular function (MF) [52][53]. A FDR correction with *Q* < 0.05 was used as the cutoff. KEGG 2021 Human was accessed through Enrichr online to find varied protein expressions within specific pathways and the overall strength of specific pathway expressions [54]. Ranking of significance was based on a combined score model and a filter of *Q* < 0.05 was applied to only consider statistically significant pathways.

### 2.6. Identification of Novel Biomarkers

This study aimed to identify potential biomarkers with diagnostic or prognostic value in diabetic nephropathy (DN) by examining differentially expressed genes (DEGs) across disease stages. The DEGs from each analysis group were cross-referenced with curated gene-disease associations in the Comparative Toxicogenomics Database (CTD) to explore their relevance to DN [55]. Additionally, a literature review was conducted to assess the novelty of specific genes in the context of DN. Some transcripts had previously been associated with non-nephropathic diabetes-related conditions or DN in non-human species. Due to limited understanding of their mechanisms and effects on the human kidney, these genes were treated as novel for this analysis.

Only protein-coding genes identified as novel were selected for further analysis. Functional enrichment analysis of these genes was performed using Enrichr and ToppFun, with a significance threshold of *Q* < 0.1, to uncover their potential involvement in metabolic pathways and biological processes relevant to the progression of DN. Co-expression analysis was conducted using the ARCHS4 RNA-seq gene-gene co-expression matrix, expanding the gene set with the top 100 co-mentioned genes. This comprehensive approach helped identify a gene with progressive fold change from early to advanced stages of DN.

## 3. Results

### 3.1. PCA Visualization

Graphical outputs of the PCA library function were analyzed to identify clear similarities in intra-group distributions while differentiating samples of different groups. The 3D representations of principal component analysis across three principal components are shown for samples from both Cohort A (**Figure 2A**) and Cohort B (**Figure 2B**). In Cohort A, a strong sample self-correlation was observed in the both EDN and ADN sample groups.

**Figure 2:**
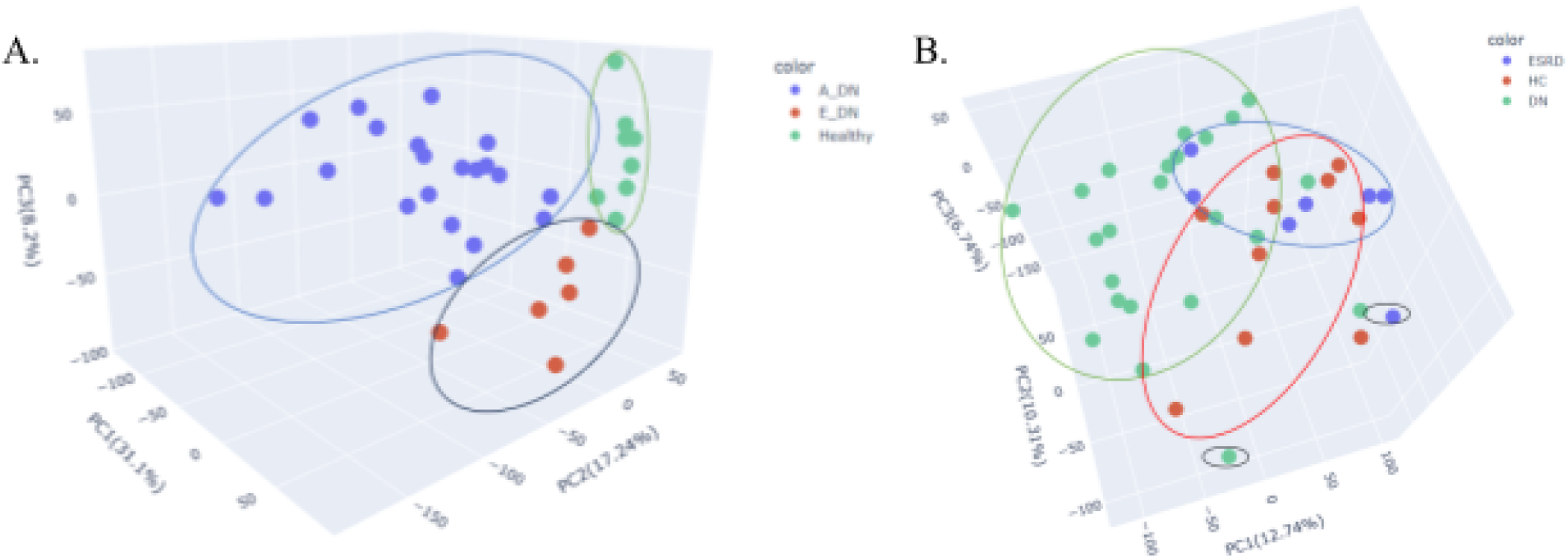
3D PCA visualizations used to identify clusters in transcriptomic similarity between defined sample groups; **A.** Cohort A (green: control; red: EDN; blue: ADN) against PC1, PC2, and PC3; **B.** Cohort B (red: control; green: DN; blue: ESRD) against the same principal components.

### 3.2. Analysis of Cohort A

#### 3.2.1. Differentially Expressed Genes

In samples diagnosed with early stage nephropathy, a total of 119 differential expressions were noted in the transcriptome when compared to the control samples (**Supplementary Table S1**); 28 upregulations and 91 downregulations comprised the altered expression (**Figures 3C and 3D**). CIDEC, MYBPC1, MYH7, MYL2, and TUSC5 displayed the largest increase in transcription, while FOSB, FOS, EGR1, NR4A1, and RGS1 maintained the greatest opposite shift in regulation (**Table 3**).

**Figure 3:**
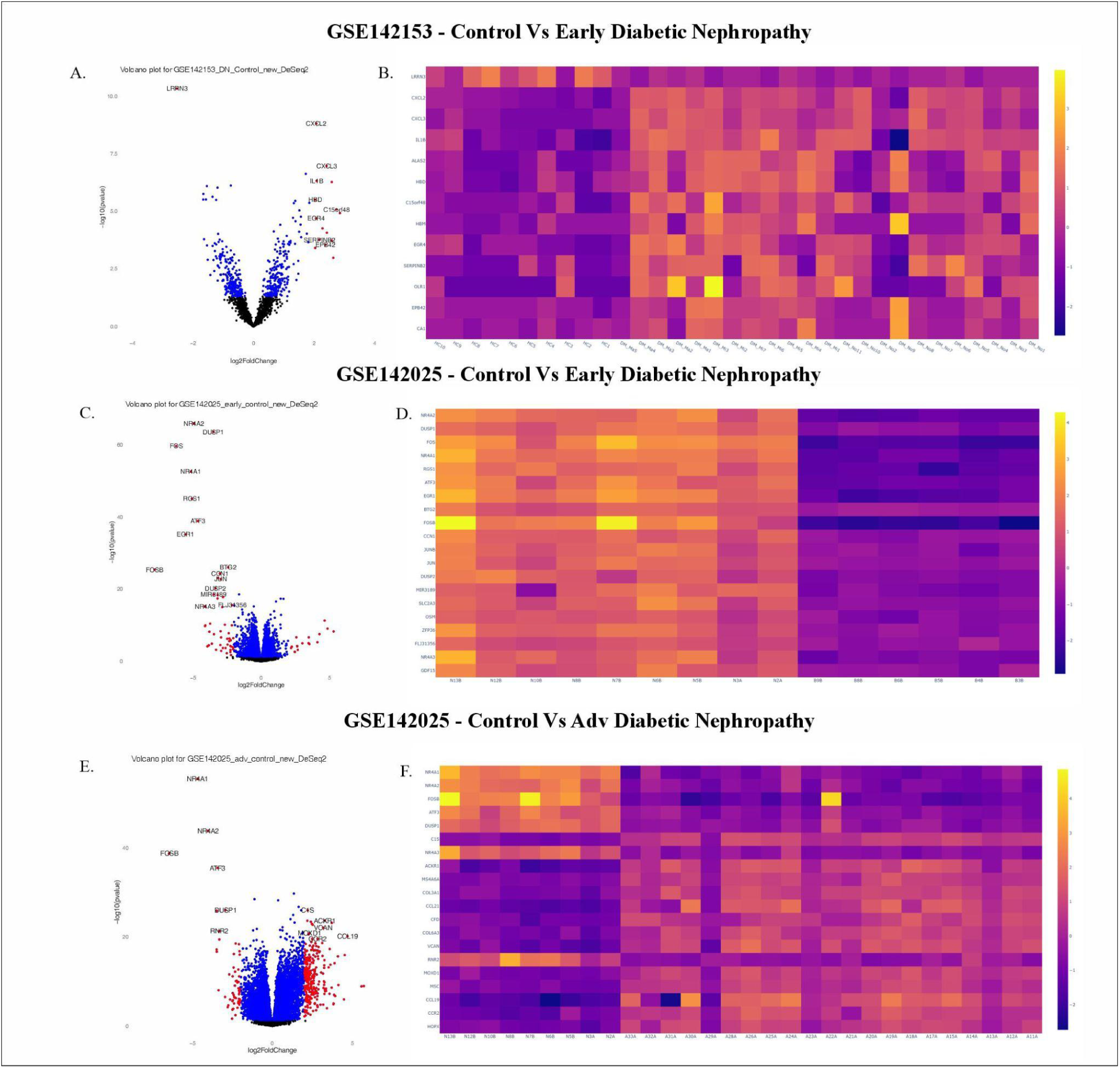
Differential gene expression volcano plot and heatmap analysis across DN-positive groups; **A.** Volcano plot visualization of DGEs from the DN-positive group of Cohort B; **B.** Heatmap representing change in key gene expressions in Cohort B; **C.** Volcano plot from Cohort A: EDN; **D.** Heatmap of DGEs from Cohort A: EDN; **E.** Volcano plot from Cohort A: ADN; **F.** Heatmap of DGEs from Cohort A: ADN.

**Table 3:**
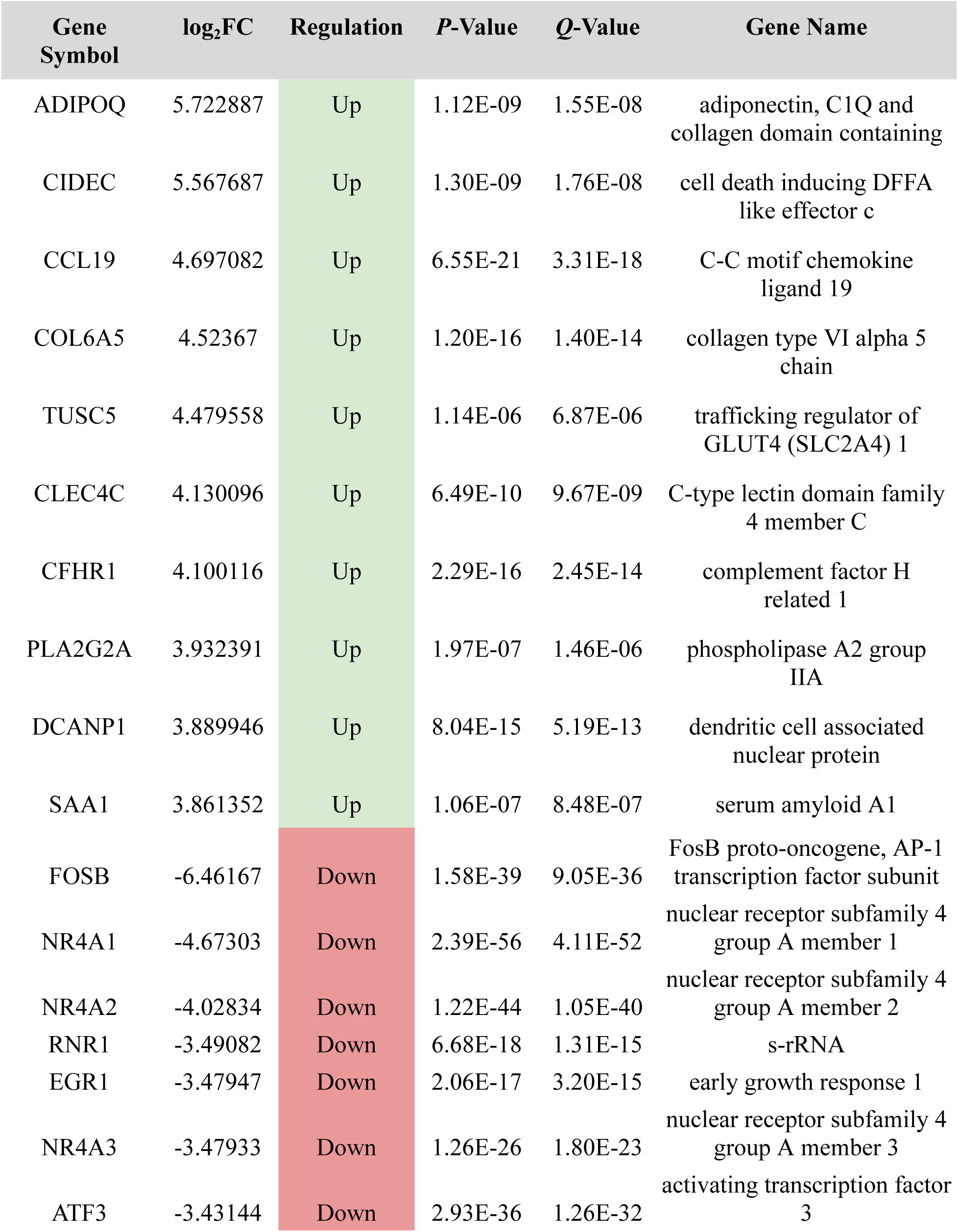
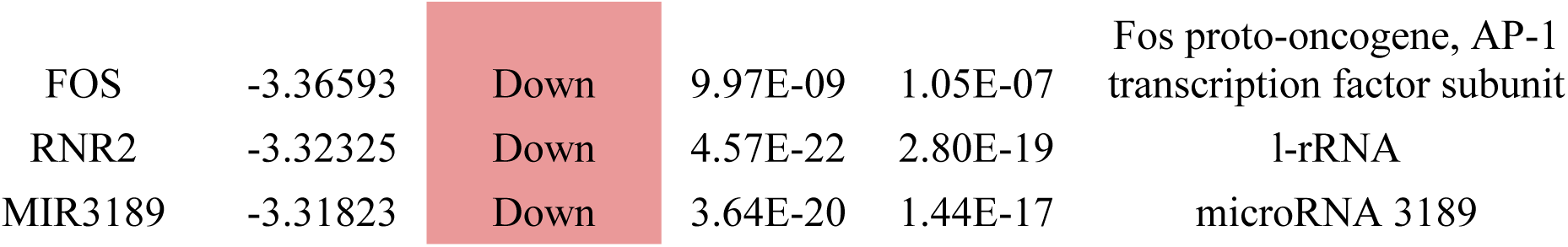
Key statistical metrics for the top 10 up- and downregulated genes in samples with EDN.

However, the ADN samples exhibited far greater variation in expression levels of protein-coding segments . The regulation of 784 genes showed a minimum absolute change in log_2_FC of 1.5, with 578 upregulated and 206 down (**Supplementary Table S2, Figures 3E and 3F**). In addition to the dysregulations in EDN samples, ADIPOQ, CCL19, and COL6A5 had log_2_FCs > 4.5, indicating severe upregulation. NR4A2, RNR1, and NR4A3 were present in the top five downregulated genes in ADN samples (**Table 4**). RNA-Seq analysis identified that drastic changes in the presence of various transcriptome RNA molecules underlied the progression from early to advanced nephropathy. In the latter stage, 39 and 10 genes showed an eight-fold change in either up- or downregulation, respectively. 279 upregulated and 58 downregulated genes possessed a minimum |log_2_FC > 2|. Therefore, a significant change in the transcriptome is noted as DN progresses from the early to advanced stage.

**Table 4:**
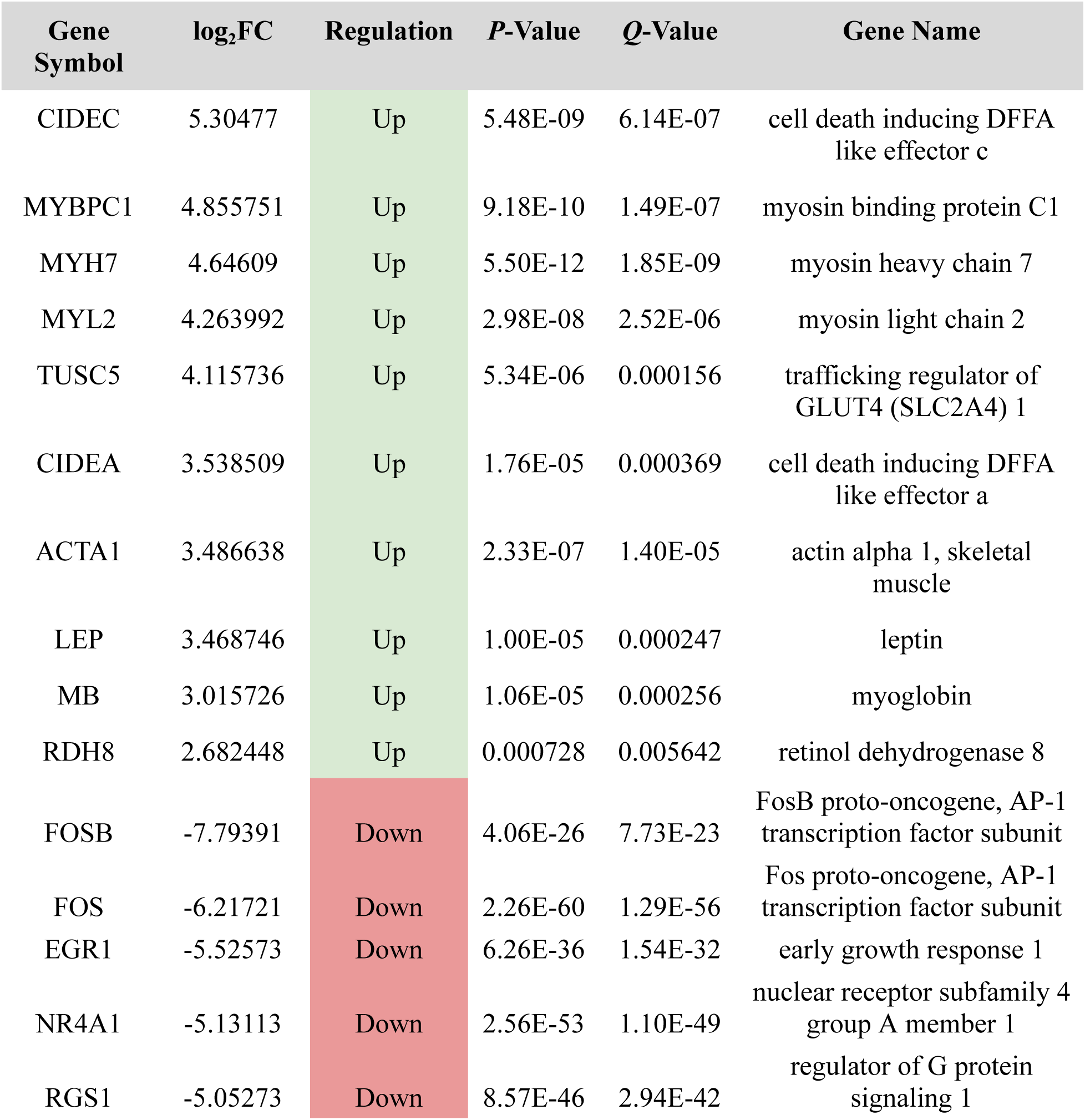
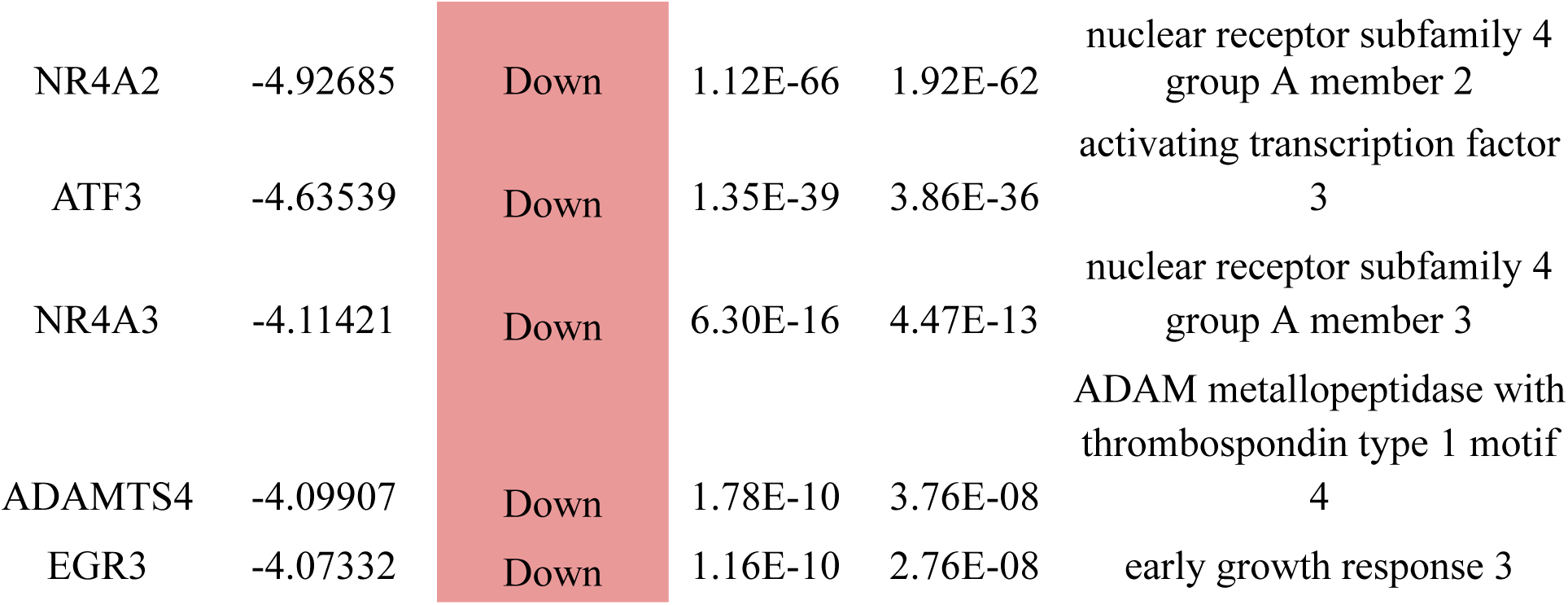
Key statistical metrics for the top 10 up- and downregulated genes in samples with ADN.

*In toto*, analyses of these two groups yielded 839 unique DGEs. Comparative analysis showed that 55 genes were differentially expressed in EDN samples only, while 720 were found to be DGEs in only ADN samples. 64 were marked statistically significant in both the early and advanced groups; CIDEC, TUSC5, LEP, ABCA13, and PLIN1 were the top five upregulated genes present in both DGE analyses, while FOSB, NR4A1, NR4A2, RNR1, and EGR1 comprised the top common downregulated genes.

#### 3.2.2. Interactome Analysis

Following compilation of DGEs, expression profiles were formed through the construction of protein networks. The interactome was built in STRING at a highest confidence of 0.900 alongside the removal of disconnected nodes.

With an enrichment *P* < 1.0e-16 to establish authority in results, the Cytoscape network formed with DGEs present in EDN possessed 66 nodes and 400 edges. Two main connected components were observed along with an average local clustering coefficient of 0.306 and node degree of 1.52. Regulation of polynucleotide adenylyltransferase activity and immediate early response, lipodystrophy, early growth response, and transcription factor AP-1 complex were the clusters with greatest strength - a measure of observed vs expected interactions. Of note, the results of a *k*-means clustering algorithm (cluster count of n = 5) showed that 27 network DGEs were involved in interleukin-10 signaling and four in lipodystrophy. The cytoHubba plug-in of Cytoscape employed the MCC algorithm to identify key regulatory nodes; hub genes present in the EDN network namely include CXCL2, FOSB, IL1B, CXCL1, ATF3, JUNB, NR4A1, FOS, IL6, CXCL8, EGR3, EGR1, EGR2, JUN, and PTGS2 (**Figure 4A**). The 15 genes possessed 62 interactions and a clustering coefficient of 0.843.

**Figure 4:**
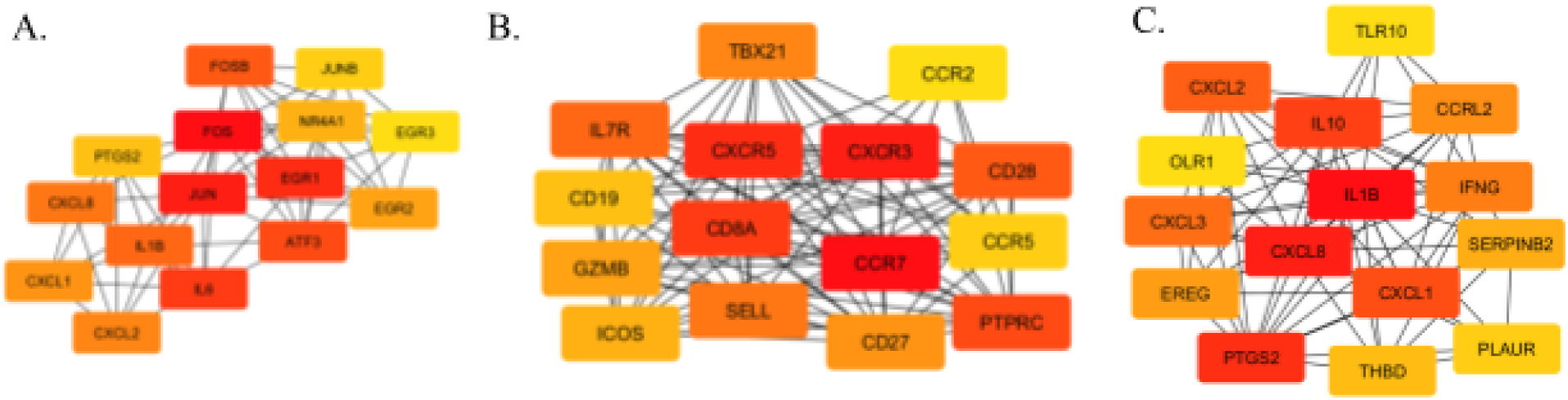
Protein-protein interaction networks were constructed in String and visualized in Cytoscape along with the networks of top 15 regulatory hub genes. They were used to identify interactions between DGEs; **A)** 66 nodes and 400 edges were observed in EDN samples in Cohort A; **B)** Hub genes from EDN network possessed 62 edges; **C)** 422 nodes and 3682 edges were seen in ADN samples in Cohort A; **D)** Hub genes from ADN network had 95 interactions between themselves; **E)** 34 nodes and 208 edges found in DN samples in Cohort B; **F)** Hub genes from DN network saw 71 edges.

In the ADN samples, the web of PPIs possessed 422 nodes and 3682 edges. A total of 13 connected components were present with an average local clustering coefficient of 0.3 and node degree of 1.79. At *Q* < 0.05, Alpha-beta T cell receptor complex, dysgammaglobulinemia & immunoglobulin domain, IgE binding, and complement component C1 complex showed the highest relative clustering strengths. A *k*-means clustering algorithm showed that 136 DGEs were associated with activation of the immune response, with CD247, CD3D, CD3E, CD3G, LCK, CD8A, ZAP70, and ITK showing prominent connectivity. In addition, 39 DGEs were present in integrin cell surface interactions, 26 in basic region leucin zipper and early growth response, 21 in chemokine-mediated signaling pathway, nine in RNA polymerase I promoter opening, and seven in retinoic acid metabolism. In Cytoscape, cytoHubba found CXCR3, CD27, GZMB, CCR5, CXCR5, CD8A, CCR2, TBX21, PTPRC, ICOS, SELL, IL7R, CD19, CD28, CCR7 as regulatory hub genes in the PPI network of differentially expressed genes in ADN samples (**Figure 4B**). Between these 15 nodes, 95 edges were present with an extremely significant clustering coefficient of 0.947.

#### 3.2.3. Functional Enrichment

Functional enrichment of both EDN and ADN DGEs consisted of pathway and gene ontology analysis. KEGG pathway analysis of differentially expressed genes from EDN samples found that IL-17 signaling, TNF signaling, viral protein interaction with cytokine and cytokine receptor, and cytokine-cytokine receptor interaction possessed the greatest combined scores at *Q* < 0.05 (**Supplementary Table S4 and Figure 5B**). As far as MF gene annotations GO, it was found that gene products of DGEs were involved in IL-8 receptor binding, MAP kinase tyrosine phosphatase activity, protein tyrosine/threonine phosphatase activity, and CXCR chemokine receptor binding. With respect to BP, chemotaxis of neutrophil, leukocyte, and cell, along with fat cell differentiation and myeloid leukocyte migration had the greatest gene ratio. In terms of CC, the top five ontologies were transcription factor AP-1 complex, cardiac myofibril, aggresome, specific granule membrane, and caveola (**Supplementary Table S5 and Figure 5A**).

**Figure 5:**
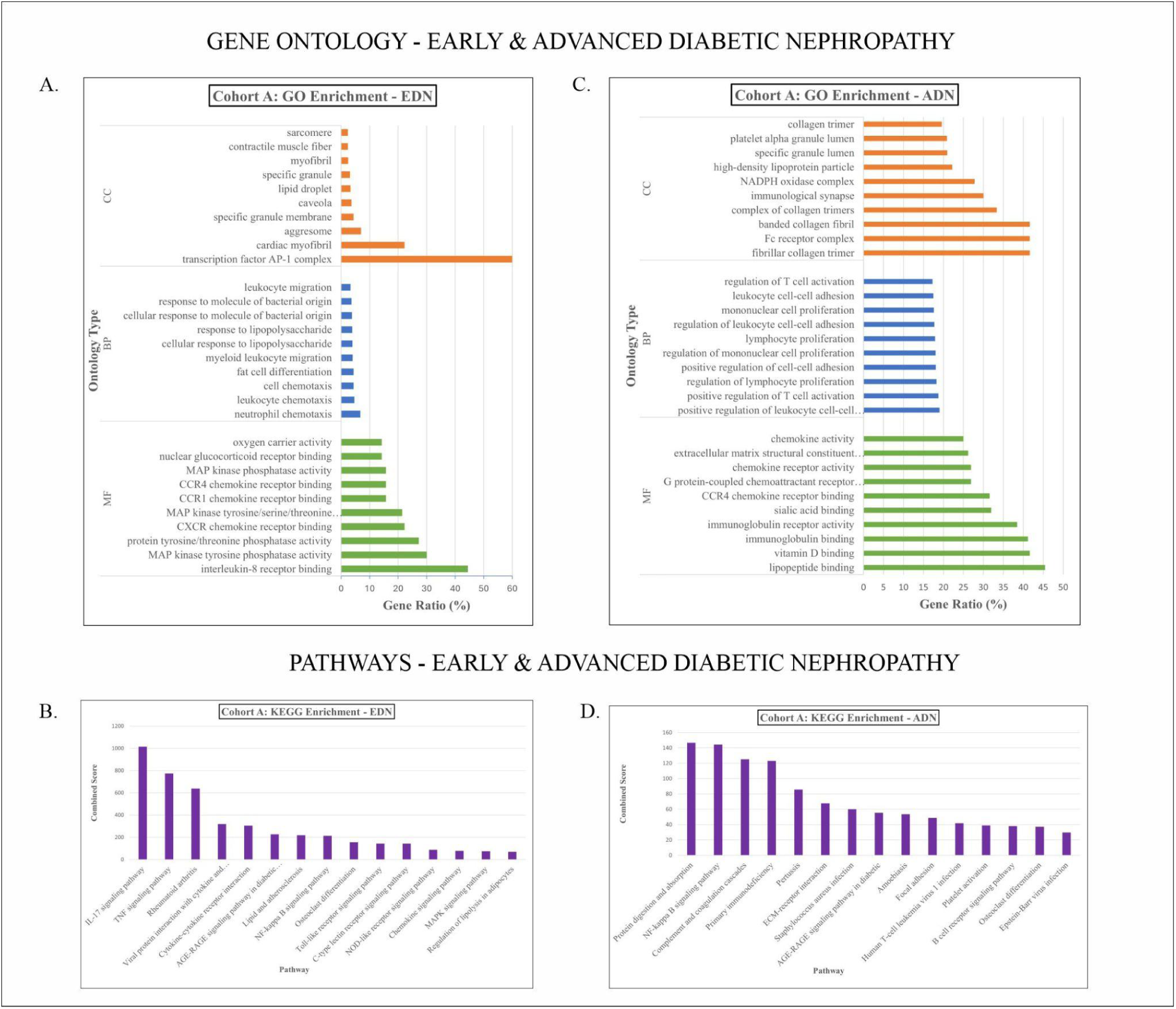
Functional enrichment results of Cohort A in terms of GO (CC, BP, and MF) and pathway modulation; **A.** Top 10 ontologies in EDN; **B.** Pathways enriched in EDN ranked by combined score; **C.** Top 10 ontologies from ADN; **D.** Pathways enriched in ADN.

In the ADN samples, pathway analysis of DGEs revealed that protein digestion and absorption, NF-kappa B signaling, complement and coagulation cascades, primary immunodeficiency, and pertussis had the highest combined scores of statistically significant KEGG pathways (**Supplementary Table S6 and Figure 5D**). Molecular function GO analysis showed C-C chemokine binding, lipopeptide binding, vitamin D binding, immunoglobulin binding, and immunoglobulin receptor activity to possess the largest share of DGEs relative to the total number of genes in the annotations (**Supplementary Table S7 and Figure 5C**). For BP, the most impactful ontologies were positive regulation of leukocyte cell-cell adhesion, positive regulation of T cell activation, regulation of lymphocyte proliferation, positive regulation of cell-cell adhesion, and regulation of mononuclear cell proliferation. In terms of CC, significant ontologies were fibrillar collagen trimer, Fc receptor complex, banded collagen fibril, complex of collagen trimmers, and immunological synapse.

### 3.3. Analysis of Cohort B

#### 3.3.1. Differentially Expressed Genes

In DN-positive samples in Cohort B, 80 differentially expressed genes were noted at fold change |Log2FC| > 1 and statistical significance *Q* < 0.05 (**Supplementary Table S3**). 46 genes were upregulated in DN transcriptomes and 34 were downregulated (**Figures 3A and 3B**). The top five upregulated transcripts were HBM, C15Orf48, HLA-DRB6, ALAS2, and OLR1, while the inverse consisted of LRRN3, SLC4A10, R11_327P2.4, UTS2, and IGKV2D-30 (**Table 5**). Compilation of these genes reveals a large portion of lncRNA-coding genes in addition to the protein-coding variants.

**Table 5:** Key statistical metrics for the top 10 up- and downregulated genes in samples with DN in Cohort B.

| Gene Symbol | log<sub>2</sub>FC | Regulation | P-Value | Q-Value | Gene Name |
| --- | --- | --- | --- | --- | --- |
| HBM | 2.854358 | Up | 1.20E-05 | 0.0017 | Hemoglobin Subunit Mu |
| C15orf48 | 2.736677 | Up | 8.34E-06 | 0.0013 | Normal Mucosa Of<br>Esophagus-Specific Gene 1<br>Protein |
| HLA-DRB6 | 2.637187 | Up | 0.001057 | 0.034054 | Major Histocompatibility<br>Complex, Class II, DR Beta<br>6 (Pseudogene) |
| ALAS2 | 2.586015 | Up | 5.41E-07 | 0.000267 | 5'-Aminolevulinate<br>Synthase 2 |
| OLR1 | 2.580787 | Up | 0.000191 | 0.013882 | Oxidized Low Density<br>Lipoprotein Receptor 1 |
| A_32_P471<br>66 | 2.42549 | Up | 8.57E-05 | 0.008196 | Long noncoding RNA |
| CXCL3 | 2.415666 | Up | 1.08E-07 | 0.000107 | C-X-C Motif Chemokine<br>Ligand 3 |
| EPB42 | 2.37307 | Up | 0.000289 | 0.01503 | Erythrocyte Membrane<br>Protein Band 4.2 |
| A_32_P488<br>2 | 2.279254 | Up | 5.56E-05 | 0.005885 | Long noncoding RNA |
| SERPINB2 | 2.176467 | Up | 0.000174 | 0.013207 | SERPINB2 |
| LRRN3 | -2.53057 | Down | 4.74E-11 | 1.40E-07 | Leucine-Rich Repeat<br>Neuronal Protein 3 |
| SLC4A10 | -1.66138 | Down | 3.11E-06 | 0.00064 | Solute Carrier Family 4<br>Member 10 |
| RP11_327P<br>2.4 | -1.65529 | Down | 1.78E-06 | 0.000527 | Clone RP11-327P2 |
| UTS2 | -1.63242 | Down | 0.000164 | 0.013162 | Urotensin 2 |
| A_32_P903<br>46 | -1.56933 | Down | 3.09E-06 | 0.00064 | Long noncoding RNA |
| ENST00000<br>474213 | -1.53822 | Down | 8.11E-07 | 0.0003 | Immunoglobulin Kappa<br>Variable 2D-30 |
| A_24_P868<br>313 | -1.53375 | Down | 0.000305 | 0.01559 | Long noncoding RNA |
| ENST00000<br>390625 | -1.51018 | Down | 0.001277 | 0.037093 | Immunoglobulin Heavy<br>Variable 3-49 |
| C2orf40 | -1.35273 | Down | 2.36E-06 | 0.000636 | ECRG4 Augurin Precursor |
| GPR34 | -1.35048 | Down | 0.000321 | 0.016112 | G Protein-Coupled Receptor<br>34 |

#### 3.3.2. Interactome Analysis

An interactome using the DGEs was modeled in STRING at a high confidence interval of 0.900. A total of 34 nodes were observed with 208 edges acting as interactions between different protein elements; two main components were present in the PPI network. Summary statistics revealed an average clustering coefficient of 0.241 and average node degree of 0.745. Primary local clusters (noted without categorization algorithm) were hemoglobin complex and ammonium transporter family, haptoglobin-hemoglobin complex, CXC chemokine, and chemokine receptors, each with a strength value of over two. Given the presence of two disconnected clusters, a k-means clustering algorithm with a number of clusters n = 3 found that nine genes were involved in IL-10 signaling, namely CXCL1, CXCL2, CXCL3, CXCL8, IL1B, IL10, IL12RB2, PTGS2, and ALOX12.

After Cytoscape was used to properly visualize the PPI network, the cytoHubba plug-in identified IL10, CXCL3, THBD, SERPINB2, CXCL8, EREG, TLR10, CCRL2, CXCL2, IFNG, PLAUR, CXCL1, IL1B, and PTGS2 as the 15 regulatory hub genes (**Figure 4C**). Between these 15 nodes, 71 connective edges existed with a clustering coefficient of 0.839.

#### 3.3.3. Functional Enrichment Analysis

Functional enrichment analysis of DGEs from the DN-positive samples of Cohort B consisted of analyzing both pathway and ontology implications. KEGG 2021 Human showed that IL-17 signaling, NF-kappa B signaling, TNF signaling, malaria, and rheumatoid arthritis had the highest combined scores in analysis (**Supplementary Table S8**). IL-8 receptor binding, CXCR chemokine receptor binding, haptoglobin binding, oxygen carrier activity, and oxidoreductase activity were the primary molecular function annotations from GO (**Supplementary Table S9**). ToppFun identified that the gene ratio of response to polysaccharide, response to molecules of bacterial origin, response to bacterium, regulation of chronic inflammatory response, and neutrophil activation were highest with regard to biological processes. The only two CC ontologies marked statistically significant were hemoglobin complex (GO:0005833) and haptoglobin-hemoglobin complex (GO:0031838).

### 3.4. Comparative Analysis

A comparison between differentially expressed genes in both groups of Cohort A and the DN-positive group of Cohort B revealed that the meta-analysis of both datasets produced a total of 911 unique DGEs (**Figure 6A**). 50 genes were present solely in the EDN analysis of the GSE142025 dataset. Likewise, 720 DGEs were present only in the analysis of ADN samples from the same dataset. Between the two sample groups from Cohort A, it was seen that 61 genes were statistically significant and of importance in both. Common to both groups, CIDEC, MYBPC1, and MYH7 were most upregulated, while FOSB, FOS, and EGR1 were most downregulated. 72 DGEs were unique to only the DN sample group of Cohort B, not appearing in either of the other two analyses of Cohort A. Five genes were found in analyses of both this group and the EDN sampling group of Cohort A: PLAUR, PTGS2, CXCL1, CXCL3, and IL1B. In all three groups, CXCL2, NR4A3, and RGS1 were present as DGEs.

**Figure 6:**
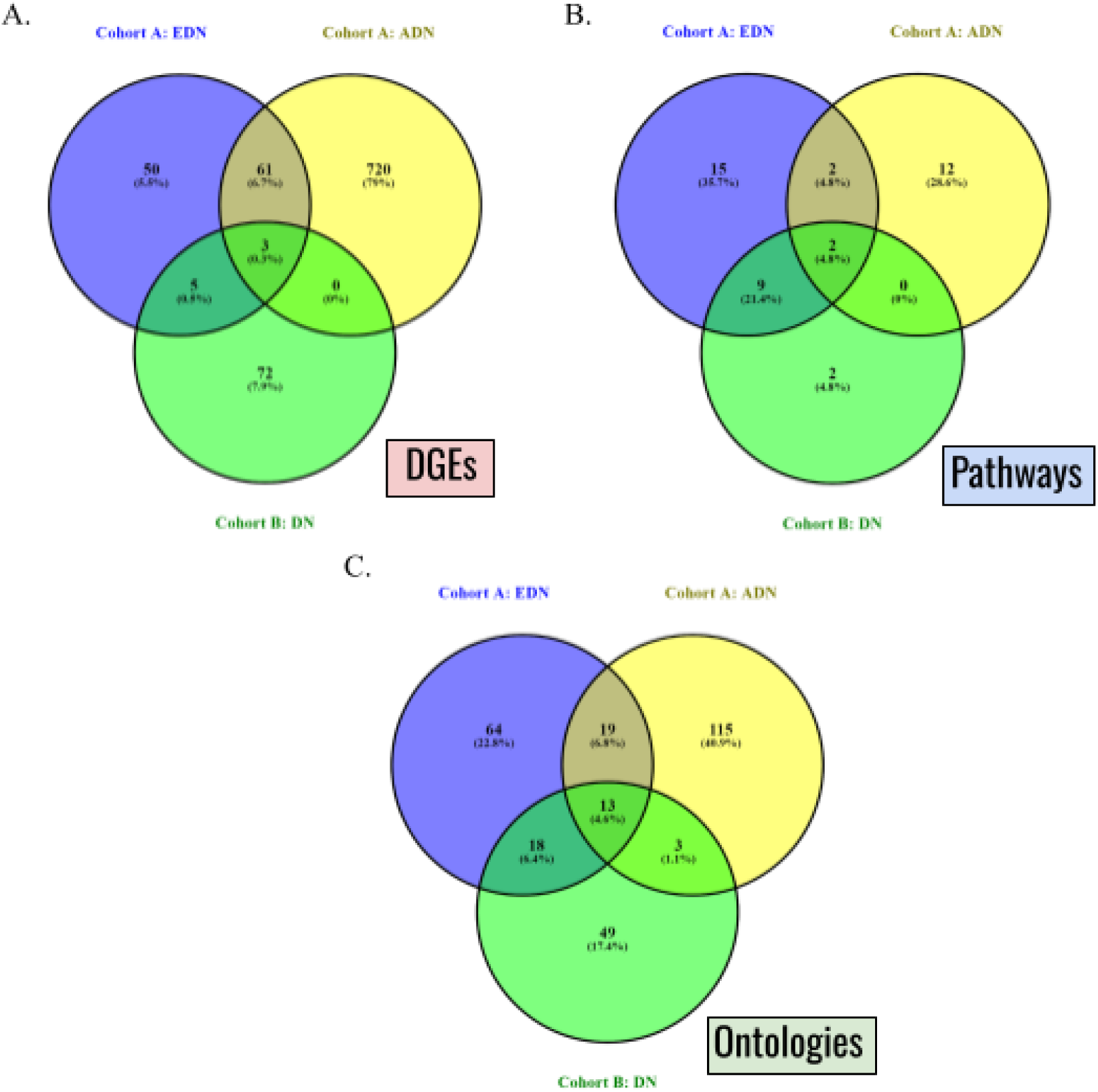
Venn diagrams representing commonalities identified between the different analysis groups; **A.** DGEs; **B.** KEGG Pathways; **C.** GO (MF, BP, and CC).

A comparison of enrichments displayed overlap between the different differential groups in terms of pathways affected (**Figure 6B**). Unique to the EDN group of the first dataset were the viral protein interaction with cytokine and cytokine receptor, toll-like receptor signaling, MAPK signaling, regulation of lipolysis in adipocytes, hypertrophic cardiomyopathy, fluid shear stress and atherosclerosis, parathyroid hormone synthesis, secretion and action, Th17 cell differentiation, apelin signaling, estrogen signaling, transcriptional misregulation in cancer, dilated cardiomyopathy, circadian entrainment, cellular senescence, and JAK-STAT signaling pathways. Differentially regulated pathways found only in the ADN samples of the same dataset were protein digestion and absorption, primary immunodeficiency, pertussis, ECM-receptor interaction, Staphylococcus aureus infection, amoebiasis, focal adhesion, human T-cell leukemia virus 1 infection, platelet activation, B cell receptor signaling, Epstein-Barr virus infection, and diabetic cardiomyopathy. DGEs from both EDN and ADN comparisons here revealed that the AGE-RAGE signaling pathway in diabetic complications and osteoclast differentiation pathway were common. Malaria and arachidonic acid metabolism were found only in the analysis of Cohort B. Nine pathways were of note in both the EDN and DN groups from Cohort A and B, respectively: IL-17 signaling, TNF signaling, rheumatoid arthritis, cytokine-cytokine receptor interaction, lipid and atherosclerosis, C-type lectin receptor signaling, NOD-like receptor signaling, chemokine signaling, and inflammatory bowel disease. Present in the functional enrichment results of all three groups were the NF-kappa B signaling pathway and complement and coagulation cascades pathway.

Similarly, commonalities between GO (all MF, BP, and CC) were identified (**Figure 6C**). 64 and 115 ontologies were found only in the EDN and ADN groups of Cohort A, respectively. Between the two groups, 19 ontologies were identified as being common. 49 ontologies were found only in the DN-positive group of Cohort B, while 18 and three ontologies in this analysis group were common with EDN and ADN groups of the other cohort, respectively. It was found that 13 ontologies were of note in the EDN, ADN, and DN-positive groups: cytokine activity, receptor ligand activity, signaling receptor regulator activity, chemokine activity, cytokine receptor binding, chemokine binding activity, signaling receptor activity, G protein-coupled binding, response to other organism, response to external biotic stimulus, response to biotic stimulus, regulation of cell adhesion, and regulation of cell-cell adhesion.

In sum, the greatest correspondence between analysis groups was observed between the EDN and ADN groups of the primary dataset. The DN-positive group of the second cohort was more closely linked to its EDN counterpart in Cohort A rather than the ADN group.

In Cohort A, specifically, pathway analysis using the pathview library in R revealed changes to pathway regulation between EDN and ADN, specifically in those involving metabolic disruption, inflammatory signaling, cell adhesion, and structural remodeling. EDN is marked by disturbances in oxidative phosphorylation and ribosomal function, which indicate mitochondrial stress and altered protein synthesis, alongside initial immune modulation and changes in cell adhesion molecules and amino acid metabolism. ADN is characterized by extracellular matrix (ECM) remodeling and oxidative stress responses, with altered ECM-receptor interactions and peroxisome pathways. Cell adhesion molecules, crucial in maintaining tissue integrity, were upregulated in ADN (**Figures 7A and 7B**). As DN advances, inflammation further intensifies with heightened activity in the JAK-STAT pathway (**Figures 7C and 7D**). Enhanced chemokine signaling and changes in hematopoietic lineage pathways imply increased recruitment and activation of immune cells within renal tissue, while significant metabolic dysregulation disrupts branched-chain amino acid and pyrimidine metabolism, impacting cellular repair and DNA synthesis.

**Figure 7:**
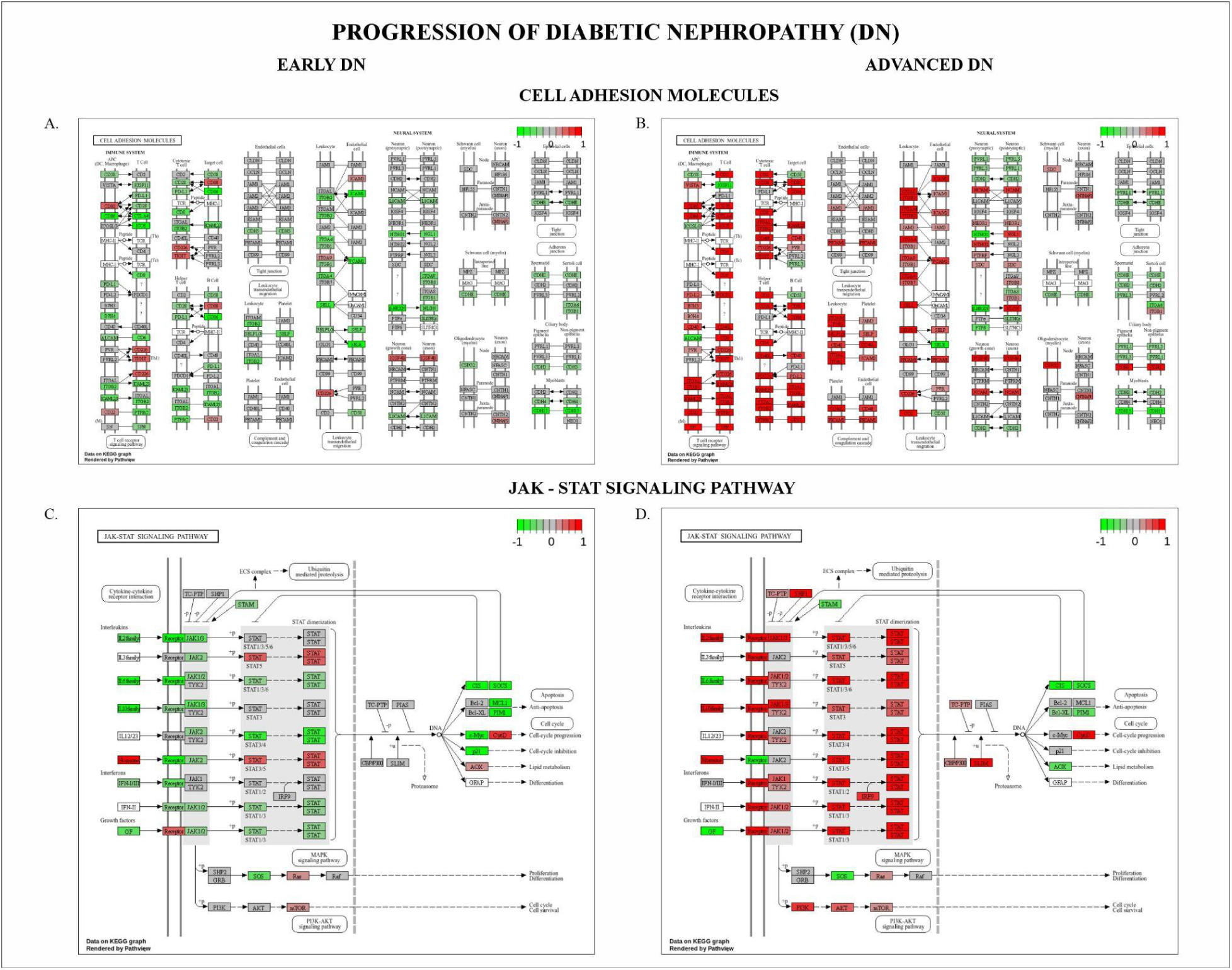
Pathview visualizations of change in pathway regulation between EDN and ADN in Cohort A; **A.** Homeostatic regulation of cell adhesion molecules is shown in EDN; **B.** Immune system is upregulated in the cell adhesion molecules pathway in ADN; **C.** Normal regulation of the JAK-STAT signaling pathway in EDN; **D.** Upregulation of cytokine-cytokine receptor interactions and STAT dimerization in the JAK-STAT signaling pathway in ADN.

### 3.5. Novel Differentially Expressed Genes in Diabetic Nephropathy

Through comparative analysis with CTD and cross-references with literature, several novel differentially expressed genes were noted in the meta-analysis; a total of 12 protein-coding genes were identified as being novel (**Table 6**). Pseudogenes were removed from the analysis given a lack of functional purpose. One protein-coding novel gene from the ADN grouping of Cohort A in particular - FCRL6 - was further investigated as candidate biomarkers with relevance to DN.

**Table 6:**
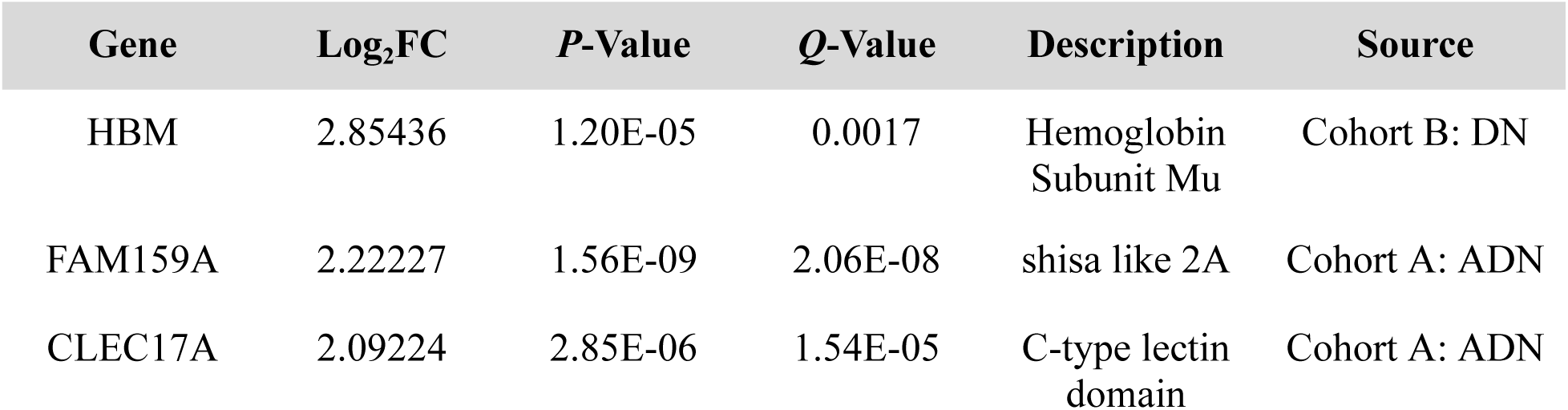
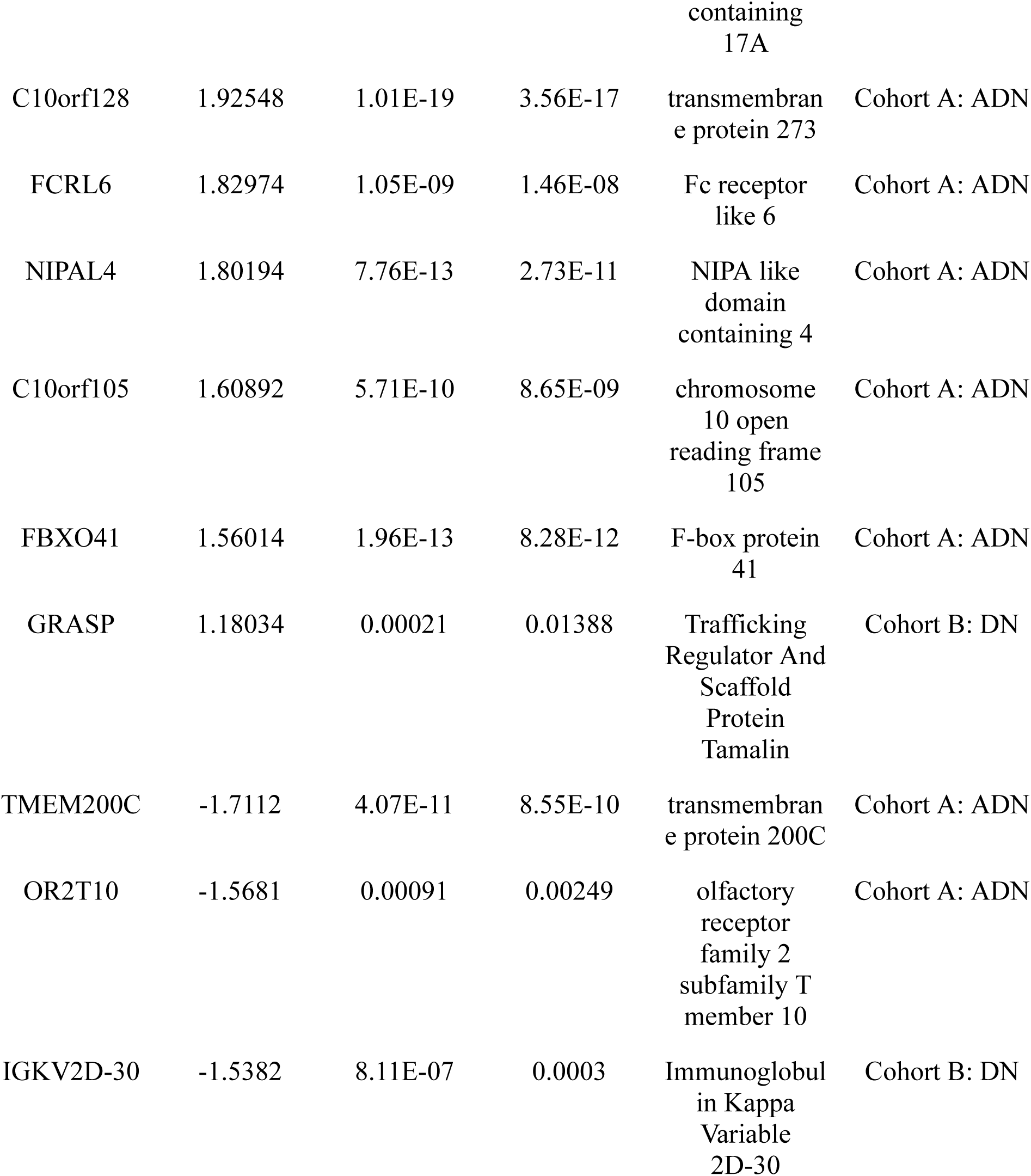
Novel protein-coding DGEs identified in ADN samples of Cohort A and DN samples of Cohort B sorted based on greatest log_2_FC.

| Gene | Log<sub>2</sub>FC | P-Value | Q-Value | Description | Source |
| --- | --- | --- | --- | --- | --- |
| HBM | 2.85436 | 1.20E-05 | 0.0017 | Hemoglobin<br>Subunit Mu | Cohort B: DN |
| FAM159A | 2.22227 | 1.56E-09 | 2.06E-08 | shisa like 2A | Cohort A: ADN |
| CLEC17A | 2.09224 | 2.85E-06 | 1.54E-05 | C-type lectin<br>domain containing<br>17A | Cohort A: ADN |
| C10orf128 | 1.92548 | 1.01E-19 | 3.56E-17 | transmembran<br>e protein 273 | Cohort A: ADN |
| FCRL6 | 1.82974 | 1.05E-09 | 1.46E-08 | Fc receptor<br>like 6 | Cohort A: ADN |
| NIPAL4 | 1.80194 | 7.76E-13 | 2.73E-11 | NIPA like<br>domain<br>containing 4 | Cohort A: ADN |
| C10orf105 | 1.60892 | 5.71E-10 | 8.65E-09 | chromosome<br>10 open<br>reading frame<br>105 | Cohort A: ADN |
| FBXO41 | 1.56014 | 1.96E-13 | 8.28E-12 | F-box protein<br>41 | Cohort A: ADN |
| GRASP | 1.18034 | 0.00021 | 0.01388 | Trafficking<br>Regulator And<br>Scaffold<br>Protein<br>Tamalin | Cohort B: DN |
| TMEM200C | -1.7112 | 4.07E-11 | 8.55E-10 | transmembran<br>e protein 200C | Cohort A: ADN |
| OR2T10 | -1.5681 | 0.00091 | 0.00249 | olfactory<br>receptor<br>family 2<br>subfamily T<br>member 10 | Cohort A: ADN |
| IGKV2D-30 | -1.5382 | 8.11E-07 | 0.0003 | Immunoglobul<br>in Kappa<br>Variable<br>2D-30 | Cohort B: DN |

### 3.6. Functional Enrichment of FCRL6

Fc Receptor Like 6 (FCRL6) is a type I transmembrane receptor and member of the Fc Receptor Like (FCRL) family of proteins with a homologous structural resemblance to leukocyte Fc receptors. The gene FCRL6 was marked differentially expressed in the ADN grouping of Cohort A solely with a log_2_FC of 1.82974 and *Q*-value of 1.22E-08. Functional enrichment revealed that FCRL6 was involved in 22 statistically significant KEGG pathways; natural killer cell mediated toxicity, primary immunodeficiency, allograft rejection, cell adhesion molecules, T cell receptor signaling pathway, and cell adhesion molecules pathways were most enriched when sorted by a combined score model (**Figure 8**).

**Figure 8:**
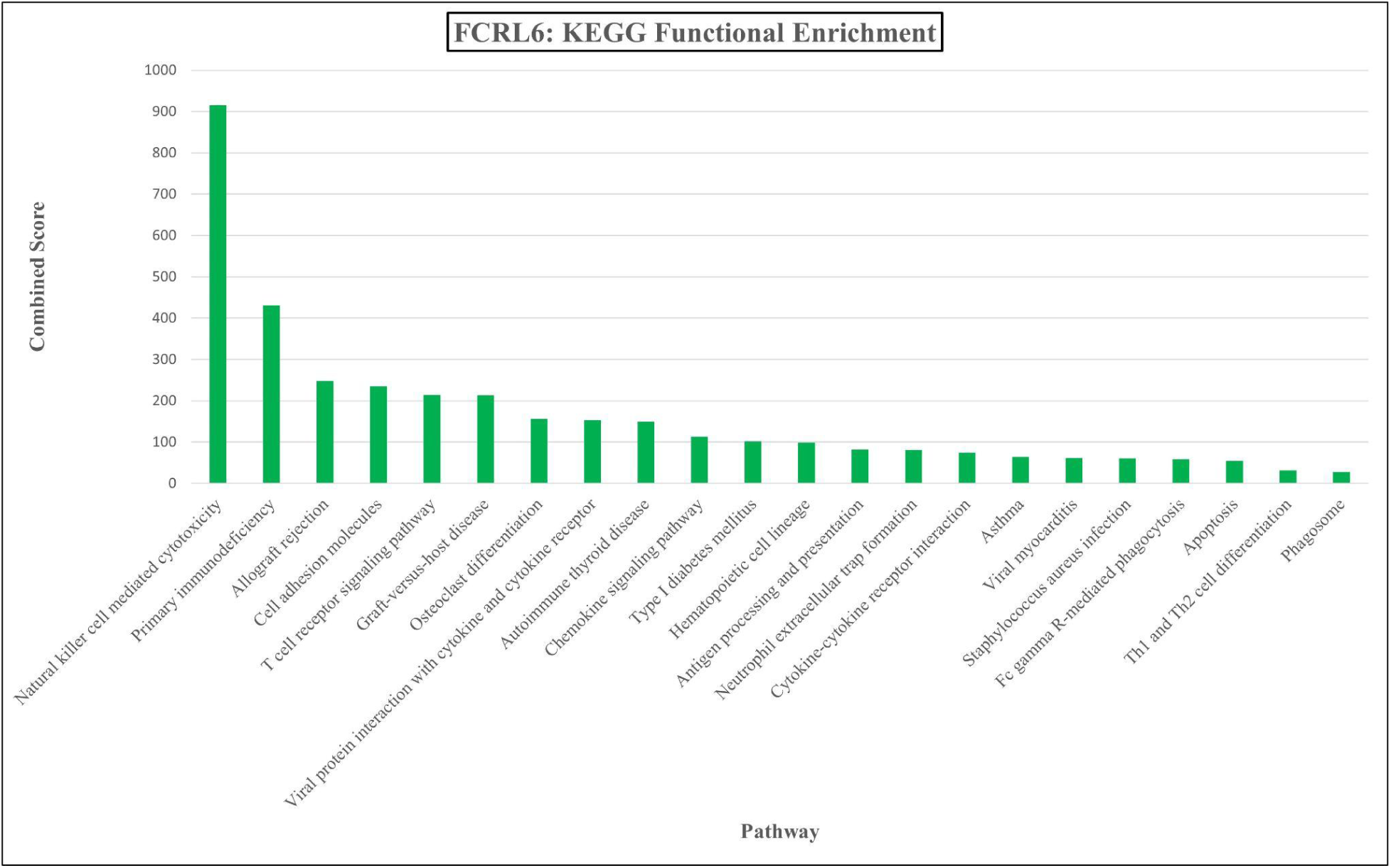
Pathway functional enrichment results of FCRL6 based on an ARCHS4 RNA-seq gene-gene co-expression matrix.

A visual comparison of these pathways with those of the analysis groups unearthed overlaps (**Figure 9**). Specifically, between FCRL6 and EDN samples from Cohort A, viral protein interaction with cytokine and cytokine receptor was affected. Between the receptor gene and ADN samples from the same dataset, both primary immunodeficiency and Staphylococcus aureus infection were found. Additionally, pathview generations of this group found FCRL6 to be involved in the overall upregulation of the cell adhesion molecules pathway. Between the three of these groups, osteoclast differentiation was common. FCRL6 and DGEs from EDN and DN samples in Cohorts A and B, respectively, were found to be involved in the chemokine signaling pathway and cytokine-cytokine receptor interaction.

**Figure 9:**
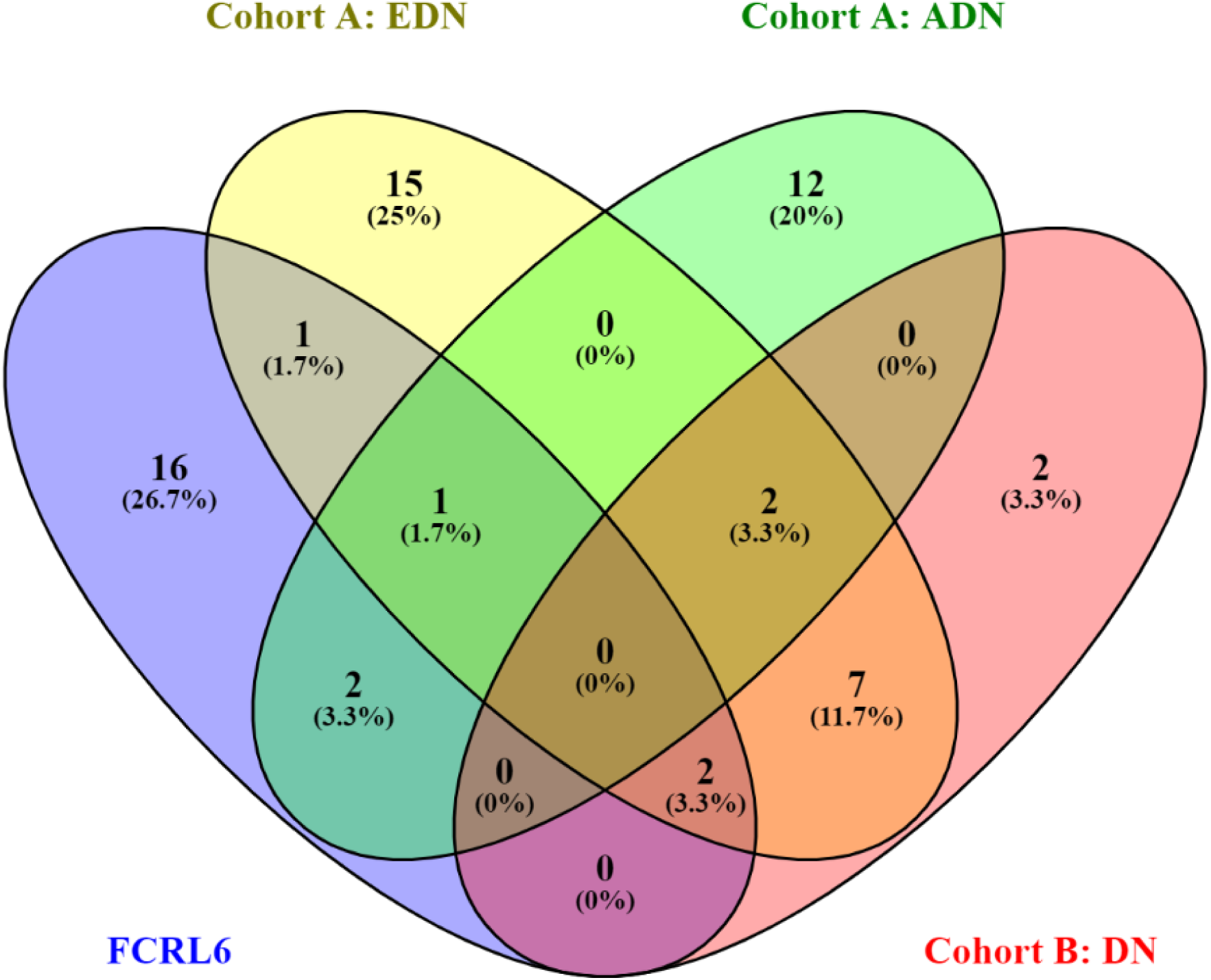
A comparison of KEGG pathways enriched in FCRL6 and the three DGE analysis groups.

In terms of molecular function gene ontologies, FCRL6 was found to be involved in MHC class II protein binding (GO:0042289), MHC protein binding (GO:0042287), protein phosphatase binding (GO:0019903), phosphatase binding (GO:0019902), transmembrane signaling receptor binding (GO:000488), and signaling receptor binding (GO:0005102). Furthermore, external side of plasma membrane (GO:0005102), side of membrane (GO:0098552), and cell surface (GO:0009986) were the enriched CC ontologies.

### 3.7. Uncovering the Role of FCRL6 in Diabetic Nephropathy

While all previous members of the FCRL family have possessed biased expression in the B cell lineage, FCRL6 is expressed preferentially - if not exclusively - in cytolytic natural killer (NK) cells, CD8+ T cells, and some populations of CD4+ T Cells [56]. Located in the plasma membrane, the external domain of this protein consists of spliced variations of five Ig-like domains [57]. Although binding between the receptor and immunoglobulins has not been noted to date, direct binding between FCRL6 and MHC II molecule human leukocyte antigen HLA-DR has been identified [58]. In both the ADN and DN analysis groups of each cohort, respectively, HLA-DR was upregulated significantly, suggesting a parallel nature in regulation between the two proteins. Given similarities in the C-regions of both Ig and MHC II molecules, a potential mechanism of action similar to Fc receptors has been linked [59]. It is predicted that cytoplasmic signal transduction is conducted through a consensus immunoreceptor tyrosine-based inhibition motif (ITIM) under homeostatic conditions [60]. Human FCRL6 proteins possess two tyrosine residues, Y356 and Y371. The prior is a canonical ITIM sequence, while the latter has been put forward as a non-canonical immunoreceptor tyrosine-based activation motif (ITAM) sequence in the cytoplasmic small tail. Intracellular tyrosyl-phosphorylation of the FCRL6 protein recruits SHP-1, SHP-2, SHIP1, and SHIP2 phosphatases, which in turn attenuate immunoreactive responses by NK cells [61][62]. Hyperactivation of SHP-1 results in the dephosphorylation of secondary signaling molecules, reversing NK cell activation [63]. It was similarly found that both SHP-1 and SHP-2 kinases are primarily activated through ITIMs and work to inhibit NK cell’s lymphocyte activity [64]. An uber-presence of SHIP-1 has previously been noted to down-regulate production of interferon gamma (IFN-*γ*) [65][66]. Members of both SHP and SHIP families have an inhibitory effect on the PI3K-Akt signaling pathway and phosphatidylinositol production. Both DM and DN are associated with the reduced presence and cytotoxicity of NK cells [67][68][69]. Various signaling pathways have been shown to inhibit the regular functioning of NK cells in hyperglycemic conditions. Therefore, specific niches filled by these innate immunity cells are left vulnerable to pathological changes. Dysregulation of necroptosis (mediated by NK cells) is a cornerstone of DN, as irregular functioning of mesangial and tubular cells leads to the dysregulation of further many pathways [70]. In addition, NK cells regulate the outflow of various cytokines such as interleukins [71]. Improper regulation of cytokines and chemokines have consistently proven to be a breeding ground for inflammatory responses in DN [72]. Thus, combined with the increased regulation of inflammatory markers and cells, a lack of NK functioning via a FCLR6/SHP/SHIP signaling upregulation could explain certain DN physiologies (**Figure 10**).

**Figure 10:**
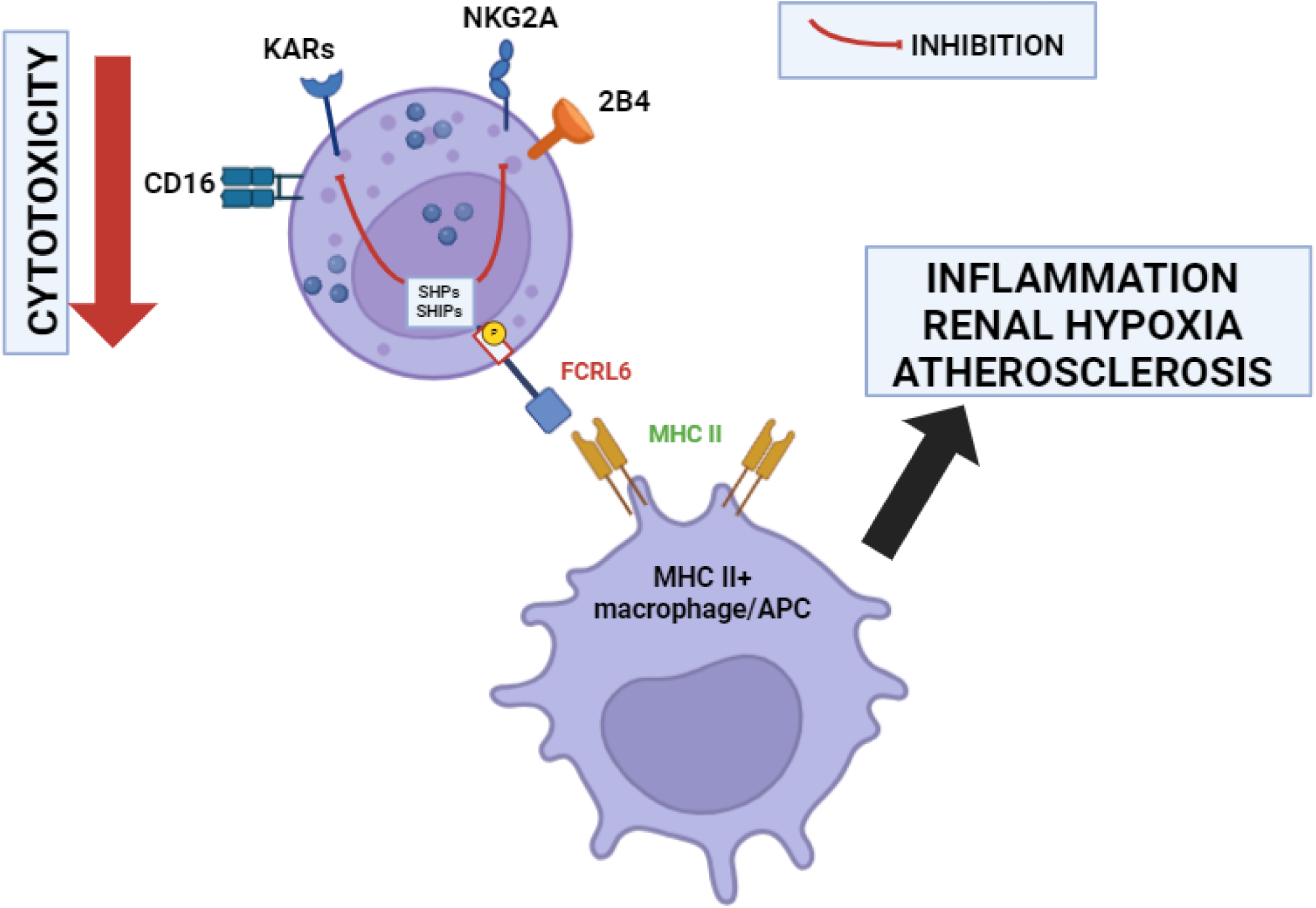
Visual representation of the mechanism by which increased FCRL6 interaction with MHC II molecules attenuates NK cell activity and cytotoxicity. The increased recruitment of SHPs and SHIPs from the FCRL6 cytoplasmic tail ITIM results in inhibition of activating receptors that mediate NK cell cytotoxicity, including CD16, NKG2A, 2B4, and KARs.

FCRL6 has been especially prominent in the discussion of immune-checkpoint therapy of various cancers, as previous research suggests its potential to act in cancer defense and tolerance via evasion of therapies [73]. Specifically, abnormal regulation of the receptor has been linked in HLA-DR+ samples with breast, lung, colorectal, and melanoma cancers following PD-1 blockade treatment [74]; from an ontogenic perspective, FCRL6 receptors biased presence in NK and cytotoxic T cells in adult human blood rather than in developmental sites such as bone marrow or the thymus [75]. This suggests a more generic mode of action in immunoregulation that is not necessarily specific beyond lymphoid tissue, aligning with the metastatic nature of diabetes. In fact, there is great similarity in the inflammatory reaction in both tumor clusters and renal structures in DN. With similar blood vessel structures in both tumors and renal hypoxia, the formation of reactive oxygen species is commonplace [76]. Inflammatory markers such as IL-1, IL-2, IL-6, IL-18, and TNF-ɑ are found to be overexpressed in both conditions [77]. Similar pathways, including the prominent JAK-STAT signaling pathway, lead to high levels of oxidative stress and inflammation [78]. Thus, the impacts of FCRL6 on DN through amelioration of NK cell-mediated cytotoxicity have been previously noted in other conditions and could help devise potential applications of protein regulation in immunotherapeutics.

## 4. Discussion

Diabetes mellitus has grown to be a global burden, with 25% of the population set to be affected by T1D, T2D, or glucose homeostasis impairment by 2030 [79][80]. DN is a microvascular renal complication that arises in a sizable percentage of individuals with DM. As a result of hyperglycemia and the upregulation of pro-inflammatory markers and pathways, physiological changes to the kidney structure are observed [81]. Proteinuria via hyperfiltration is used as a diagnostic tool in kidney disorders such as DN and has been the main method used to classify DN by stages [82]. In finding key disease-causing genes, bioinformatics-based approaches have recently grown massively in their usage given their potential to identify possible biomarkers based on a fold-change approach [83].

In this investigation, we found 911 unique differentially expressed genes, with 635 being upregulated and 276 downregulated. A majority of these genes have been previously linked to DN as either prognostic or diagnostic biomarkers. ADIPOQ exists in polymorphic states and is the primary gene associated with heightened serum plasma adiponectin levels [84]. CIDEC is affected by the attenuation of EGR1-1 proliferation, inhibiting the decomposition of lipids in podocytes present in the glomerular filtration barrier, increasing albuminuria [85]. CCL19 promotes fibrogenesis and inflammation in DN [86]. LEP is involved in its namesake pathway that increases the secretion of TGF-I1 in vascular endothelial cells of the kidney [87]. Both DUSP1 and FOSB were found to be hub genes in pro-inflammatory gene networks of samples with immunoglobulin nephropathy [88]. NR4A1 regulation impacts DN via mitochondrial fission activation and mitophagy repression [89]. Significant downregulation of NR4A2 stimulates myocardial injury in DN via the HDAC11 pathway [90]. FOS has been implicated in MAPK signaling and other inflammatory pathways when downregulated in DN, potentially serving as a candidate therapeutic target [91]. It has been proposed that elevated urinary levels of CCL21 mRNA could serve as an alternative diagnostic method in comparison with uACR and eGFR calculations [92]. With a potential link to FCRL6 in NK cells, HLA-DR antigens have been noted to be upregulated in DN [93]. SERPINE1 expands the effects of renal injury after diabetic insult through intracellular communication and paracrine signaling [94].

In the PPI network of DGEs from EDN samples in Cohort A, CXCL2, FOSB, IL1B, CXCL1, ATF3, JUNB, NR4A1, FOS, IL6, CXCL8, EGR3, EGR1, EGR2, JUN, and PTGS2 were hub genes, In the network of DGEs from ADN samples of the same dataset, CXCR3, CD27, GZMB, CCR5, CXC5, CD8A, CCR2, TBX21, PTPRC, ICOS, SELL, IL7R, CD19, CD28, and CCR7 were key regulatory genes. In the network of DN-positive samples of Cohort B, IL10, CXCL3, THBD, SERPINB2, CXCL8, EREG, TLR10, CCRL2, CXCL2, IFNG, PLAUR, CXCL1, IL1B, and PTGS2 were identified as hub genes. C-X-C chemokine motifs are able to orchestrate significant chemotaxis of adaptive immune cells [95]. In addition, they are found to be upregulated in both glomeruli and proximal tubules - the sites of maximum inflammation in DN [96]. JUN, ATF3, and JUNB have been seen as significant genes in monocyte infiltration in early DN tissue [97]. Co-stimulatory molecules CD19, CD27, and CD28 are found on T cells and are essential for activation and inflammatory responses in nephropathy [98][99]. PTGS2 has been found to be downregulated in DN, but involved in the inflammatory response [100].

A series of functional studies and bioinformatics-based functional enrichments have implicated various pathways in DN found here as well. The formation of advanced glycation end products stimulates RAGE activation, in turn stimulating the formation of reactive oxygen species and other damaging products [101]. Several interleukins have been displayed to take a part in the inflammatory response of DN; IL-6 receptors and gp130 form active dimer complexes involved in trans-signaling pathways on leukocytes and hepatocytes, IL-18 is involved in the release of serum cytokines, and IL-17 is responsible for cytokine secretion in Th17 cells [102][103][104]. Chemokine CXC receptor motifs recruit neutrophils and cause inflammation of renal tissue [105]. Protein digestion and absorption has downstream effects of apoptosis in renal tubules [106]. Activated in the glomeruli and tubules of DN patients, NF-κB pathways induce increased transcription of chemotactic protein-coding genes, drawing monocyte and neutrophil infiltrations [107]. Complement and coagulation cascades due to hyperglycemia often lead to increased autoreactivity and accelerated protein glycation, increasing immune deposition in the kidney [108]. In addition to these key pathways, progression of DN is built upon the interaction of many more as significant genetic and proteomic dysregulation has a cascading effect on the intracellular and intercellular landscape of the kidney.

Current treatments of DN primarily are linked with controlling blood sugar and electrolyte intake [109]. Individuals with more advanced stages of DN typically undergo weekly or bi-weekly hemodialysis procedures to extract excess water, salts, and toxins not filtered out by the inflamed kidney [110]. Likewise in later stages, nephrologists may suggest a kidney transplant to replace the dysfunctioning kidney [111]. In both dialysis and organ transplant treatment options, there are significant downsides associated. Quality of life gradually reduces over the years-long span of hemodialysis for DN-positive patients [112]. In some cases of kidney transplants, the arisal of graft-vs-host disease complicates matters; the condition possesses a near 75% mortality rate [113]. Therefore, a promising new potential method of treating nephropathy in diabetes (in addition to diabetes mellitus itself) is immunotherapy [114]. The treatment refers to the deployment of site-specific molecules to reduce inflammatory responses that have a cyclic effect on pathogenic progression. In DM type I, immunotherapy has been put forward as a treatment option in DN that uses receptor-blocking molecules to suppress both adaptive and innate immune responses. Thus far, several immuno-modulating molecules have been tested in DN. GLP-1 receptor agonists have been described to regulate blood glucose concentrations and have been selectively deployed in renal tissue recovered from nephrectomy samples [115]. By activating pathways that downregulate RAGE, this treatment mechanism has shown to possess potential in future treatment avenues. A particular study found that finerenone has statistically significant results in reducing kidney failure and chronic kidney disease progression in clinical trials, both of which follow similar inflammatory patterns to DN [116]. Sodium-glucose cotransporter 2 (SGLT2) inhibitors have been put forward as an add-on therapy to DM and DN treatments given their glycosuric action [117]. Angiotensin-converting enzyme (ACE) inhibitors were found to slow nephropathy onset and progression in diabetes independent of blood pressure effects [118]. While additional research is required to fully understand the side effects of immunosuppressive treatments over a long term in nephropathy treatment, the immuno-modulating potential presents a significant opportunity to control lymphocytic attacks on the kidney.

Fc Receptor Like 6 (FCRL6) is a receptor found exclusively in natural killer and cytotoxic T cells (along with rare appearances in other cytotoxic T cell variations). Differential expression of this protein was found in advanced DN samples of Cohort A during analysis and was functionally enriched in natural kill cell-mediated cytotoxicity, primary immunodeficiency, cell adhesion molecules, and T cell receptor signaling pathways. Its selective dysregulation in ADN suggests it could have potential as a diagnostic biomarker in DN. Additionally, immunotherapy treatments could introduce FCRL antagonists to the blood system, reactivating migrating NK cells’ cytotoxicity and reestablish their role as immunoregulators. To further probe the potential of FCRL6 - and other FCRL proteins for that matter - in treatments requires *in vivo* experimentation with possible clinical trials.

## 5. Limitations and Recommendations

This investigation used solely a bioinformatics-based approach in analyzing two datasets with the aim of identifying potential prognostic or diagnostic biomarkers in diabetic nephropathy. The study lacks *in vitro* experimentation and the biological evidence achieved through functional studies. Similarly, the implication that FCRL6 could serve as a biomarker in DN was done through functional enrichment and literature-bases analyses. To further corroborate these results, there is a need to perform clinical studies; animal studies with a FCRL6 gene knockout would elucidate the validity of this receptor as a biomarker and make headway into the idea of FCRL proteins as potential targets of immunotherapy in DN. Equally, further investigation is required to fully understand the mechanisms by which FCRL6 and downstream pathways impact NK cells. The approach used here provides a comprehensive integrated perspective on different aspects of DN physiology using meta-analysis techniques and provides many areas in which further exploration could take place. Thus, this research builds on a web of existing studies using bioinformatics, furthering our knowledge of the transcriptome of the diabetic kidney.

## Acknowledgements

We would like to thank the authors who deposited their transcriptomic read count data on the NCBI GEO. They additionally acknowledge the research community for their interconnected exchange of ideas and analyses upon which this study is predicated. We further acknowledge OmicsLogic and the Tauber Bioinformatics Research Center for providing the server on which the differential gene expression pipeline was run.

## Footnotes

### Ethical Statement

This study does not contain any studies with human subjects or animals performed by any of the authors.

### Conflicts of Interest

The authors declare that they have no conflicts of interest to this work.

## Data Availability Statement

The present study utilized data from two datasets on the National Center for Biotechnology Information’s Gene Expression Omnibus for analysis: GSE142025 (https://www.ncbi.nlm.nih.gov/geo/query/acc.cgi?acc=GSE142025) and GSE142153 (https://www.ncbi.nlm.nih.gov/geo/query/acc.cgi?acc=GSE142153).

## Author Contribution Statement

**Adarsh Marinedrive**: Conceptualization, formal analysis, investigation, methodology, visualization, writing - original draft. **Sonalika Ray**: Software, validation, visualization, writing - review & editing. **Mohit Mazumder**: Validation, writing - review & editing.

## Notes

https://www.ncbi.nlm.nih.gov/geo/query/acc.cgi?acc=GSE142025

https://www.ncbi.nlm.nih.gov/geo/query/acc.cgi?acc=GSE142153

### Competing Interest Statement

The authors have declared no competing interest.

