## Supplementary Tables S1-S3 for "Transcriptomic-Based Identification of FCRL6 as a Novel Diagnostic Biomarker in Diabetic Nephropathy"

**Table S1**: Differentially expressed genes from Cohort A: EDN. A total of 119 dysregulations were observed – 28 upregulations and 91 downregulations.

| Gene Symbol | $log_2FC$ | *P*-Value | *Q*-Value | Gene Name |
|---|---|---|---|---|
| CIDEC | 5.30477 | 5.48E-09 | 6.14E-07 | cell death inducing DFFA like effector c |
| MYBPC1 | 4.855751 | 9.18E-10 | 1.49E-07 | myosin binding protein C1 |
| MYH7 | 4.64609 | 5.50E-12 | 1.85E-09 | myosin heavy chain 7 |
| MYL2 | 4.263992 | 2.98E-08 | 2.52E-06 | myosin light chain 2 |
| TUSC5 | 4.115736 | 5.34E-06 | 0.000156 | trafficking regulator of GLUT4 (SLC2A4) 1 |
| CIDEA | 3.538509 | 1.76E-05 | 0.000369 | cell death inducing DFFA like effector a |
| ACTA1 | 3.486638 | 2.33E-07 | 1.40E-05 | actin alpha 1, skeletal muscle |
| LEP | 3.468746 | 1.00E-05 | 0.000247 | leptin |
| MB | 3.015726 | 1.06E-05 | 0.000256 | myoglobin |
| RDH8 | 2.682448 | 0.000728 | 0.005642 | retinol dehydrogenase 8 |
| WNT7B | 2.647617 | 7.57E-05 | 0.001053 | Wnt family member 7B |
| XIRP2 | 2.636626 | 6.00E-05 | 0.000895 | xin actin binding repeat containing 2 |
| PLIN1 | 2.581025 | 2.27E-07 | 1.37E-05 | perilipin 1 |
| TNNT1 | 2.471501 | 0.00274 | 0.015332 | troponin T1, slow skeletal type |
| HBG2 | 2.464596 | 0.002421 | 0.013939 | hemoglobin subunit gamma 2 |
| SCARA5 | 2.243669 | 9.12E-05 | 0.001205 | scavenger receptor class A member 5 |
| ABCA13 | 2.191475 | 0.000215 | 0.00228 | ATP binding cassette subfamily A member 13 |
| FABP4 | 1.968079 | 2.28E-06 | 8.18E-05 | fatty acid binding protein 4 |

| | | | | |
|---|---|---|---|---|
| TMC1 | 1.914014 | 0.011706 | 0.045592 | transmembrane channel like 1 |
| MROH2B | 1.870692 | 0.002552 | 0.014498 | maestro heat like repeat family member 2B |
| CKM | 1.85799 | 3.56E-05 | 0.000607 | creatine kinase, M-type |
| DES | 1.851118 | 1.42E-05 | 0.000318 | desmin |
| EEF1A2 | 1.808861 | 4.93E-05 | 0.00078 | eukaryotic translation elongation factor 1 alpha 2 |
| ADCYAP1R1 | 1.808569 | 0.000307 | 0.002963 | ADCYAP receptor type I |
| WISP2 | 1.735987 | 0.008874 | 0.037041 | cellular communication network factor 5 |
| CFD | 1.666365 | 3.09E-06 | 0.000104 | complement factor D |
| MIR27B | 1.63988 | 4.99E-18 | 4.28E-15 | microRNA 27b |
| AGXT | 1.526709 | 1.16E-05 | 0.000271 | alanine--glyoxylate and serine--pyruvate aminotransferase |
| CSRNP1 | -1.51235 | 0.000105 | 0.001339 | cysteine and serine rich nuclear protein 1 |
| CLEC4E | -1.5132 | 0.010157 | 0.041115 | C-type lectin domain family 4 member E |
| HBEGF | -1.54117 | 0.001341 | 0.008877 | heparin binding EGF like growth factor |
| SLC28A2 | -1.55275 | 0.00021 | 0.002245 | solute carrier family 28 member 2 |
| IGF2 | -1.55857 | 0.011634 | 0.045435 | insulin like growth factor 2 |
| CR2 | -1.56122 | 0.012375 | 0.047559 | complement C3d receptor 2 |
| NFKBIZ | -1.56445 | 5.28E-07 | 2.68E-05 | NFKB inhibitor zeta |
| MYC | -1.56509 | 0.00206 | 0.012325 | MYC proto-oncogene, bHLH transcription factor |

| | | | | |
|---|---|---|---|---|
| PLAUR | -1.58484 | 1.43E-06 | 5.85E-05 | plasminogen activator, urokinase receptor |
| SLCO4A1_AS1 | -1.58915 | 0.002353 | 0.013638 | NA |
| RGS2 | -1.61064 | 2.84E-19 | 3.34E-16 | regulator of G protein signaling 2 |
| FCGR3B | -1.6187 | 8.44E-05 | 0.00114 | Fc fragment of IgG receptor IIIb |
| PTGS2 | -1.64928 | 9.59E-13 | 4.01E-10 | prostaglandin-endoperoxide synthase 2 |
| CEACAM3 | -1.65141 | 0.002809 | 0.015639 | CEA cell adhesion molecule 3 |
| BRE_AS1 | -1.65993 | 2.00E-05 | 0.000402 | NA |
| MROH5 | -1.67869 | 8.90E-06 | 0.000226 | maestro heat like repeat family member 5 (gene/pseudogene) |
| VNN3 | -1.68017 | 0.004771 | 0.023055 | vanin 3 |
| C11orf96 | -1.70572 | 2.18E-10 | 4.51E-08 | chromosome 11 open reading frame 96 |
| RNR2 | -1.75021 | 0.003401 | 0.018124 | l-rRNA |
| ICOS | -1.75244 | 5.11E-05 | 0.000796 | inducible T cell costimulator |
| RNR1 | -1.75965 | 0.003603 | 0.01888 | s-rRNA |
| S100A8 | -1.76792 | 5.22E-08 | 3.98E-06 | S100 calcium binding protein A8 |
| F2RL3 | -1.76906 | 0.002794 | 0.015571 | F2R like thrombin or trypsin receptor 3 |
| CXCL1 | -1.78318 | 4.07E-06 | 0.000127 | C-X-C motif chemokine ligand 1 |
| LOC284454 | -1.80221 | 9.02E-06 | 0.000228 | NA |
| DUSP5 | -1.81372 | 1.62E-07 | 1.04E-05 | dual specificity phosphatase 5 |
| PDK4 | -1.8162 | 1.55E-05 | 0.000337 | pyruvate dehydrogenase kinase 4 |
| IL1RN | -1.83674 | 7.96E-05 | 0.001093 | interleukin 1 receptor antagonist |
| APOLD1 | -1.84437 | 0.001181 | 0.008167 | apolipoprotein L domain containing 1 |
| CTLA4 | -1.84677 | 0.002503 | 0.014279 | cytotoxic T-lymphocyte |

| | | | | |
|---|---|---|---|---|
| | | | | associated protein 4 |
| CCL8 | -1.87247 | 1.16E-05 | 0.000271 | C-C motif chemokine ligand 8 |
| CXCL3 | -1.9088 | 0.000113 | 0.001407 | C-X-C motif chemokine ligand 3 |
| LOC101927472 | -1.91388 | 9.66E-11 | 2.40E-08 | NA |
| KLF6 | -1.92779 | 1.29E-05 | 0.000294 | Kruppel like factor 6 |
| FCAR | -1.95279 | 0.001779 | 0.011033 | Fc fragment of IgA receptor |
| MIR23A | -1.9605 | 3.04E-05 | 0.000542 | microRNA 23a |
| RGS16 | -1.9624 | 3.54E-16 | 2.64E-13 | regulator of G protein signaling 16 |
| GUCY2EP | -1.96543 | 0.000961 | 0.006972 | guanylate cyclase 2E, pseudogene |
| CD69 | -1.98155 | 1.26E-11 | 3.92E-09 | CD69 molecule |
| TNFSF9 | -1.9948 | 2.29E-05 | 0.000445 | TNF superfamily member 9 |
| HSPA1B | -2.03646 | 4.15E-07 | 2.21E-05 | heat shock protein family A (Hsp70) member 1B |
| PADI4 | -2.04092 | 0.000475 | 0.004118 | peptidyl arginine deiminase 4 |
| INHBB | -2.04666 | 0.000141 | 0.001667 | inhibin subunit beta B |
| FLJ31356 | -2.11888 | 2.83E-16 | 2.21E-13 | uncharacterized protein FLJ31356 |
| GADD45B | -2.13977 | 1.10E-10 | 2.66E-08 | growth arrest and DNA damage inducible beta |
| CCDC144A | -2.18852 | 8.75E-05 | 0.001172 | coiled-coil domain containing 144A |
| LINC00473 | -2.2158 | 1.59E-06 | 6.34E-05 | phosphodiesterase 10A |
| LUCAT1 | -2.23568 | 0.000791 | 0.006002 | lung cancer associated transcript 1 |
| PLA2G2D | -2.24337 | 0.002882 | 0.015938 | phospholipase A2 group IID |
| SERPINE1 | -2.24671 | 5.72E-05 | 0.000864 | serpin family E member 1 |
| LOC221946 | -2.3097 | 5.00E-07 | 2.57E-05 | uncharacterized LOC221946 |

| | | | | |
|---|---|---|---|---|
| CXCL8 | -2.3293 | 2.66E-06 | 9.24E-05 | C-X-C motif chemokine ligand 8 |
| TREM1 | -2.36897 | 1.68E-08 | 1.55E-06 | triggering receptor expressed on myeloid cells 1 |
| BTG2 | -2.40989 | 9.03E-27 | 1.94E-23 | BTG anti-proliferation factor 2 |
| CH25H | -2.45334 | 2.45E-05 | 0.000471 | cholesterol 25-hydroxylase |
| IL1B | -2.4758 | 6.11E-07 | 3.00E-05 | interleukin 1 beta |
| ADAMDEC1 | -2.50616 | 0.00607 | 0.027756 | ADAM like decysin 1 |
| PROK2 | -2.59845 | 0.000218 | 0.0023 | prokineticin 2 |
| RASD1 | -2.65081 | 7.60E-11 | 1.98E-08 | ras related dexamethasone induced 1 |
| OSM | -2.78854 | 1.76E-18 | 1.77E-15 | oncostatin M |
| SELE | -2.79514 | 0.000212 | 0.002253 | selectin E |
| GDF15 | -2.84273 | 9.94E-16 | 6.56E-13 | growth differentiation factor 15 |
| UBD | -2.91773 | 0.001384 | 0.009108 | ubiquitin D |
| MAFF | -2.92624 | 2.40E-07 | 1.44E-05 | MAF bZIP transcription factor F |
| JUN | -2.94774 | 1.69E-23 | 2.41E-20 | Jun proto-oncogene, AP-1 transcription factor subunit |
| SOCS3 | -2.96257 | 4.91E-09 | 5.70E-07 | suppressor of cytokine signaling 3 |
| CCL20 | -2.9786 | 3.20E-05 | 0.000561 | C-C motif chemokine ligand 20 |
| CYR61 | -3.0179 | 4.79E-25 | 8.23E-22 | cellular communication network factor 1 |
| JUNB | -3.14378 | 7.89E-24 | 1.23E-20 | JunB proto-oncogene, AP-1 transcription factor subunit |
| SLC2A3 | -3.18338 | 4.54E-19 | 4.87E-16 | solute carrier family 2 member 3 |
| ZFP36 | -3.18407 | 3.89E-18 | 3.71E-15 | ZFP36 ring finger protein |

| | | | | |
|---|---|---|---|---|
| RNF17 | -3.29118 | 0.000706 | 0.005519 | ring finger protein 17 |
| DUSP2 | -3.3359 | 5.68E-21 | 7.50E-18 | dual specificity phosphatase 2 |
| CXCL2 | -3.44554 | 3.14E-08 | 2.58E-06 | C-X-C motif chemokine ligand 2 |
| MIR3189 | -3.47088 | 2.92E-19 | 3.34E-16 | microRNA 3189 |
| DUSP1 | -3.49315 | 2.51E-64 | 2.15E-60 | dual specificity phosphatase 1 |
| AKR1B10 | -3.59953 | 1.37E-05 | 0.000308 | aldo-keto reductase family 1 member B10 |
| CCL3 | -3.72404 | 3.84E-11 | 1.08E-08 | C-C motif chemokine ligand 3 |
| EGR2 | -3.78033 | 6.66E-09 | 7.18E-07 | early growth response 2 |
| IL6 | -3.8201 | 3.75E-05 | 0.00063 | interleukin 6 |
| USP32P1 | -3.93013 | 7.43E-05 | 0.001038 | ubiquitin specific peptidase 32 pseudogene 1 |
| EGR3 | -4.07332 | 1.16E-10 | 2.76E-08 | early growth response 3 |
| ADAMTS4 | -4.09907 | 1.78E-10 | 3.76E-08 | ADAM metallopeptidase with thrombospondin type 1 motif 4 |
| NR4A3 | -4.11421 | 6.30E-16 | 4.47E-13 | nuclear receptor subfamily 4 group A member 3 |
| ATF3 | -4.63539 | 1.35E-39 | 3.86E-36 | activating transcription factor 3 |
| NR4A2 | -4.92685 | 1.12E-66 | 1.92E-62 | nuclear receptor subfamily 4 group A member 2 |
| RGS1 | -5.05273 | 8.57E-46 | 2.94E-42 | regulator of G protein signaling 1 |
| NR4A1 | -5.13113 | 2.56E-53 | 1.10E-49 | nuclear receptor subfamily 4 group A member 1 |
| EGR1 | -5.52573 | 6.26E-36 | 1.54E-32 | early growth response 1 |
| FOS | -6.21721 | 2.26E-60 | 1.29E-56 | Fos proto-oncogene, AP-1 transcription factor subunit |

| | | | | |
|---|---|---|---|---|
| FOSB | -7.79391 | 4.06E-26 | 7.73E-23 | FosB proto-oncogene, AP-1 transcription factor subunit |

**Table S2**: Differentially expressed genes from Cohort A: ADN. A total of 784 dysregulations were observed – 578 upregulations and 206 downregulations.

| Gene Symbol | $log_2FC$ | *P*-Value | *Q*-Value | Gene Name |
|---|---|---|---|---|
| ADIPOQ | 5.722887 | 1.12E-09 | 1.55E-08 | adiponectin, C1Q and collagen domain containing |
| CIDEC | 5.567687 | 1.30E-09 | 1.76E-08 | cell death inducing DFFA like effector c |
| CCL19 | 4.697082 | 6.55E-21 | 3.31E-18 | C-C motif chemokine ligand 19 |
| COL6A5 | 4.52367 | 1.20E-16 | 1.40E-14 | collagen type VI alpha 5 chain |
| TUSC5 | 4.479558 | 1.14E-06 | 6.87E-06 | trafficking regulator of GLUT4 (SLC2A4) 1 |
| CLEC4C | 4.130096 | 6.49E-10 | 9.67E-09 | C-type lectin domain family 4 member C |
| CFHR1 | 4.100116 | 2.29E-16 | 2.45E-14 | complement factor H related 1 |
| PLA2G2A | 3.932391 | 1.97E-07 | 1.46E-06 | phospholipase A2 group IIA |
| DCANP1 | 3.889946 | 8.04E-15 | 5.19E-13 | dendritic cell associated nuclear protein |
| SAA1 | 3.861352 | 1.06E-07 | 8.48E-07 | serum amyloid A1 |
| GABBR1 | 3.830382 | 9.27E-06 | 4.36E-05 | gamma-aminobutyric acid type B receptor subunit 1 |
| LEP | 3.818211 | 1.41E-05 | 6.35E-05 | leptin |
| CCL21 | 3.714891 | 6.86E-24 | 6.20E-21 | C-C motif chemokine ligand 21 |
| LTF | 3.707305 | 2.60E-09 | 3.24E-08 | lactotransferrin |
| TIFAB | 3.63457 | 1.76E-14 | 1.03E-12 | TIFA inhibitor |
| IRF4 | 3.58925 | 5.17E-18 | 1.06E-15 | interferon regulatory factor 4 |
| STMN2 | 3.575162 | 6.06E-08 | 5.18E-07 | stathmin 2 |
| TUBB3 | 3.517496 | 6.88E-14 | 3.37E-12 | tubulin beta 3 class III |
| FGG | 3.411908 | 0.000107 | 0.000376 | fibrinogen gamma chain |
| IGLL5 | 3.357206 | 1.33E-08 | 1.35E-07 | immunoglobulin lambda like polypeptide 5 |
| REG3G | 3.351887 | 4.10E-09 | 4.79E-08 | regenerating family member 3 gamma |

| | | | | |
|---|---|---|---|---|
| LILRA4 | 3.294428 | 1.65E-08 | 1.64E-07 | leukocyte immunoglobulin like receptor A4 |
| SLPI | 3.283682 | 8.66E-09 | 9.24E-08 | secretory leukocyte peptidase inhibitor |
| ACKR1 | 3.281036 | 2.10E-24 | 2.40E-21 | atypical chemokine receptor 1 (Duffy blood group) |
| REG1A | 3.26105 | 1.67E-10 | 2.95E-09 | regenerating family member 1 alpha |
| LINC01426 | 3.255812 | 1.65E-15 | 1.32E-13 | long intergenic non-protein coding RNA 1426 |
| TCL1A | 3.25572 | 6.58E-08 | 5.58E-07 | TCL1 family AKT coactivator A |
| SPIB | 3.24641 | 2.37E-14 | 1.34E-12 | Spi-B transcription factor |
| ABCA13 | 3.239983 | 8.85E-14 | 4.12E-12 | ATP binding cassette subfamily A member 13 |
| SIRPG_AS1 | 3.238015 | 6.13E-10 | 9.20E-09 | NA |
| MZB1 | 3.225446 | 9.51E-11 | 1.79E-09 | marginal zone B and B1 cell specific protein |
| GREM1 | 3.20067 | 2.97E-09 | 3.64E-08 | gremlin 1, DAN family BMP antagonist |
| CLC | 3.182474 | 1.71E-05 | 7.50E-05 | Charcot-Leyden crystal galectin |
| VCAN | 3.178103 | 7.66E-23 | 5.73E-20 | versican |
| CXCL6 | 3.16848 | 1.70E-12 | 5.32E-11 | C-X-C motif chemokine ligand 6 |
| SERPINA3 | 3.134758 | 1.83E-16 | 2.02E-14 | serpin family A member 3 |
| CADM3 | 3.1186 | 2.30E-19 | 7.06E-17 | cell adhesion molecule 3 |
| CD79A | 3.049118 | 9.96E-13 | 3.37E-11 | CD79a molecule |
| CD5L | 3.00701 | 1.78E-05 | 7.76E-05 | CD5 molecule like |
| CD3D | 2.959815 | 1.70E-17 | 2.75E-15 | CD3d molecule |
| CD1C | 2.933625 | 2.63E-17 | 3.93E-15 | CD1c molecule |
| FCRL3 | 2.925927 | 1.08E-10 | 2.00E-09 | Fc receptor like 3 |
| CD48 | 2.890295 | 1.79E-18 | 4.22E-16 | CD48 molecule |
| CTSG | 2.88313 | 4.74E-10 | 7.37E-09 | cathepsin G |
| FCRL2 | 2.882932 | 5.97E-09 | 6.67E-08 | Fc receptor like 2 |

| | | | | |
|---|---|---|---|---|
| CHI3L2 | 2.869313 | 8.57E-16 | 7.71E-14 | chitinase 3 like 2 |
| FAP | 2.849324 | 2.14E-16 | 2.34E-14 | carboxyl ester lipase |
| PTCRA | 2.831822 | 5.69E-05 | 0.000215 | pre T cell antigen receptor alpha |
| FCRL5 | 2.831434 | 4.96E-08 | 4.35E-07 | Fc receptor like 5 |
| IKZF3 | 2.830608 | 1.10E-17 | 1.97E-15 | IKAROS family zinc finger 3 |
| CCR2 | 2.826163 | 2.78E-20 | 1.16E-17 | C-C motif chemokine receptor 2 |
| TMEM132D | 2.819389 | 2.03E-07 | 1.50E-06 | transmembrane protein 132D |
| PLIN1 | 2.80543 | 2.43E-05 | 0.000102 | perilipin 1 |
| LINC00402 | 2.790274 | 1.11E-06 | 6.73E-06 | long intergenic non-protein coding RNA 402 |
| MMP7 | 2.788449 | 1.08E-17 | 1.97E-15 | matrix metallopeptidase 7 |
| CLEC10A | 2.783311 | 4.92E-17 | 6.55E-15 | C-type lectin domain containing 10A |
| HIST1H4L | 2.773101 | 2.92E-12 | 8.62E-11 | H4 clustered histone 13 |
| FAM30A | 2.755208 | 1.65E-08 | 1.64E-07 | family with sequence similarity 30 member A |
| CNR2 | 2.750527 | 2.98E-09 | 3.65E-08 | cannabinoid receptor 2 |
| LCK | 2.74625 | 1.15E-18 | 2.82E-16 | LCK proto-oncogene, Src family tyrosine kinase |
| CD1E | 2.738771 | 7.31E-14 | 3.53E-12 | CD1e molecule |
| HLA_DRB5 | 2.736458 | 0.000309 | 0.00096 | NA |
| SCEL | 2.699238 | 2.81E-08 | 2.61E-07 | sciellin |
| CD2 | 2.698138 | 3.02E-19 | 8.78E-17 | CD2 molecule |
| FCER2 | 2.695326 | 1.13E-08 | 1.17E-07 | Fc fragment of IgE receptor II |
| FCRL1 | 2.691836 | 5.20E-08 | 4.53E-07 | Fc receptor like A |
| HAS2 | 2.688949 | 2.42E-19 | 7.28E-17 | hyaluronan synthase 2 |
| ANKRD36BP2 | 2.688034 | 2.05E-08 | 1.98E-07 | ankyrin repeat domain 36B pseudogene 2 |
| COL1A1 | 2.685309 | 2.59E-16 | 2.74E-14 | collagen type I alpha 1 chain |
| NTM | 2.672757 | 9.39E-18 | 1.77E-15 | neurotrimin |
| FCRLA | 2.665517 | 3.57E-07 | 2.47E-06 | Fc receptor like A |
| PAX5 | 2.662105 | 8.55E-06 | 4.05E-05 | paired box 5 |
| SYT16 | 2.658415 | 2.19E-13 | 9.01E-12 | synaptotagmin 16 |

| | | | | |
|---|---|---|---|---|
| LAX1 | 2.656076 | 4.21E-13 | 1.58E-11 | lymphocyte transmembrane adaptor 1 |
| ELK2AP | 2.645624 | 1.07E-05 | 4.96E-05 | ETS transcription factor ELK2A, pseudogene |
| SLAMF7 | 2.642521 | 1.74E-12 | 5.43E-11 | SLAM family member 7 |
| LYZ | 2.641993 | 6.36E-13 | 2.29E-11 | lysozyme |
| MMP9 | 2.63728 | 3.87E-09 | 4.58E-08 | matrix metallopeptidase 9 |
| SIRPG | 2.623754 | 6.78E-12 | 1.78E-10 | signal regulatory protein gamma |
| CST6 | 2.606131 | 3.84E-08 | 3.47E-07 | cystatin E/M |
| BLK | 2.595157 | 3.92E-09 | 4.61E-08 | BLK proto-oncogene, Src family tyrosine kinase |
| TDO2 | 2.587752 | 2.01E-09 | 2.60E-08 | tryptophan 2,3-dioxygenase |
| IL7R | 2.585986 | 1.86E-11 | 4.29E-10 | interleukin 7 receptor |
| SCARA5 | 2.580559 | 2.75E-10 | 4.56E-09 | scavenger receptor class A member 5 |
| HIST1H3I | 2.58055 | 2.11E-15 | 1.60E-13 | H3 clustered histone 11 |
| IRX6 | 2.580218 | 6.56E-11 | 1.30E-09 | iroquois homeobox 6 |
| SIT1 | 2.580067 | 1.11E-17 | 1.97E-15 | signaling threshold regulating transmembrane adaptor 1 |
| MYBPC2 | 2.56872 | 9.70E-08 | 7.83E-07 | myosin binding protein C2 |
| SIGLEC6 | 2.567624 | 3.38E-12 | 9.80E-11 | sialic acid binding Ig like lectin 6 |
| JAML | 2.566534 | 1.11E-16 | 1.33E-14 | junction adhesion molecule like |
| RHOH | 2.560364 | 1.44E-11 | 3.42E-10 | ras homolog family member H |
| WFDC2 | 2.55934 | 9.46E-19 | 2.43E-16 | WAP four-disulfide core domain 2 |
| GZMK | 2.553166 | 7.08E-15 | 4.68E-13 | granzyme K |
| FUT9 | 2.547784 | 0.000457 | 0.001354 | fucosyltransferase 9 |
| TNC | 2.540564 | 2.55E-13 | 1.02E-11 | tenascin C |
| SNX20 | 2.524901 | 6.20E-18 | 1.25E-15 | sorting nexin 20 |
| COL6A3 | 2.524518 | 1.77E-23 | 1.45E-20 | collagen type VI alpha 3 chain |
| DHRS9 | 2.522759 | 7.59E-08 | 6.34E-07 | dehydrogenase/reductase 9 |
| LCN2 | 2.520494 | 6.50E-12 | 1.72E-10 | lipocalin 2 |
| EOMES | 2.512277 | 7.49E-16 | 6.89E-14 | eomesodermin |

| | | | | |
|---|---|---|---|---|
| FAM129C | 2.511535 | 2.37E-08 | 2.26E-07 | niban apoptosis regulator 3 |
| SFRP2 | 2.507591 | 1.16E-08 | 1.20E-07 | secreted frizzled related protein 2 |
| PRRX1 | 2.506624 | 1.88E-18 | 4.36E-16 | paired related homeobox 1 |
| MEOX1 | 2.50621 | 1.68E-10 | 2.95E-09 | mesenchyme homeobox 1 |
| HOPX | 2.490368 | 2.86E-20 | 1.17E-17 | HOP homeobox |
| ERFE | 2.487976 | 7.07E-12 | 1.85E-10 | erythroferrone |
| CFD | 2.477163 | 1.21E-23 | 1.04E-20 | complement factor D |
| UBASH3A | 2.475454 | 5.40E-15 | 3.73E-13 | ubiquitin associated and SH3 domain containing A |
| P2RX5 | 2.470151 | 2.71E-08 | 2.54E-07 | purinergic receptor P2X 5 |
| CD52 | 2.457007 | 1.08E-10 | 2.00E-09 | CD52 molecule |
| FABP4 | 2.450192 | 6.87E-07 | 4.41E-06 | fatty acid binding protein 4 |
| TREML2 | 2.449713 | 9.13E-08 | 7.44E-07 | triggering receptor expressed on myeloid cells like 2 |
| CD180 | 2.436684 | 1.37E-19 | 4.52E-17 | CD180 molecule |
| LOC100507616 | 2.436656 | 7.54E-07 | 4.79E-06 | NA |
| CADM3_AS1 | 2.435051 | 4.47E-09 | 5.17E-08 | NA |
| ITGAD | 2.434388 | 2.45E-10 | 4.13E-09 | integrin subunit alpha D |
| NNMT | 2.433791 | 5.26E-10 | 8.05E-09 | nicotinamide N-methyltransferase |
| CORO1A | 2.430498 | 8.60E-18 | 1.64E-15 | coronin 1A |
| TRAT1 | 2.429906 | 3.78E-10 | 6.05E-09 | T cell receptor associated transmembrane adaptor 1 |
| ITGB6 | 2.429785 | 1.95E-09 | 2.52E-08 | integrin subunit beta 6 |
| SIGLEC8 | 2.428847 | 1.99E-09 | 2.57E-08 | sialic acid binding Ig like lectin 8 |
| FCER1A | 2.417072 | 2.23E-11 | 5.02E-10 | Fc fragment of IgE receptor Ia |
| FUT7 | 2.416169 | 1.21E-10 | 2.21E-09 | fucosyltransferase 7 |
| TPSAB1 | 2.411208 | 1.42E-09 | 1.90E-08 | tryptase alpha/beta 1 |
| CD79B | 2.406839 | 1.54E-12 | 4.91E-11 | CD79b molecule |
| COL3A1 | 2.406006 | 3.79E-24 | 3.83E-21 | collagen type III alpha 1 chain |
| ZNF831 | 2.404264 | 4.15E-13 | 1.56E-11 | zinc finger protein 831 |
| CPA3 | 2.397688 | 3.81E-10 | 6.08E-09 | carboxypeptidase A3 |
| BATF | 2.387304 | 2.87E-11 | 6.25E-10 | basic leucine zipper ATF-like transcription factor |

| | | | | |
|---|---|---|---|---|
| CCL17 | 2.384065 | 8.43E-05 | 0.000304 | C-C motif chemokine ligand 17 |
| RETN | 2.380259 | 0.003074 | 0.007277 | resistin |
| MS4A1 | 2.377349 | 7.15E-07 | 4.57E-06 | membrane spanning 4-domains A1 |
| C16orf54 | 2.373066 | 3.43E-16 | 3.45E-14 | chromosome 16 open reading frame 54 |
| CCL18 | 2.370868 | 4.88E-06 | 2.47E-05 | C-C motif chemokine ligand 18 |
| MIR4539 | 2.366887 | 7.10E-06 | 3.44E-05 | microRNA 4539 |
| SP140 | 2.361767 | 1.50E-14 | 8.81E-13 | SP140 nuclear body protein |
| DRP2 | 2.361524 | 2.93E-07 | 2.06E-06 | dihydropyrimidinase like 2 |
| CD3E | 2.357654 | 1.78E-15 | 1.41E-13 | CD3e molecule |
| MIR4537 | 2.345824 | 6.64E-06 | 3.25E-05 | microRNA 4537 |
| LOC100996286 | 2.345674 | 1.16E-05 | 5.29E-05 | NA |
| TIMP1 | 2.343941 | 8.80E-19 | 2.36E-16 | TIMP metallopeptidase inhibitor 1 |
| LINC01215 | 2.338882 | 2.04E-08 | 1.98E-07 | long intergenic non-protein coding RNA 1215 |
| IL2RB | 2.338775 | 1.20E-12 | 3.98E-11 | interleukin 2 receptor subunit beta |
| PLAC8 | 2.338473 | 3.59E-12 | 1.02E-10 | galectin 13 |
| C3 | 2.337564 | 3.54E-11 | 7.52E-10 | complement C3 |
| CPZ | 2.335907 | 6.43E-13 | 2.30E-11 | carboxypeptidase Z |
| IKZF1 | 2.335585 | 1.23E-15 | 1.03E-13 | IKAROS family zinc finger 1 |
| POU2AF1 | 2.334858 | 3.15E-07 | 2.19E-06 | POU class 2 homeobox associating factor 1 |
| MROH2B | 2.334755 | 2.31E-08 | 2.21E-07 | maestro heat like repeat family member 2B |
| LSP1 | 2.330345 | 2.60E-15 | 1.95E-13 | lymphocyte specific protein 1 |
| CD7 | 2.32907 | 5.03E-14 | 2.57E-12 | CD7 molecule |
| MOXD1 | 2.323852 | 1.36E-21 | 7.80E-19 | monooxygenase DBH like 1 |
| PIK3CD_AS1 | 2.321501 | 1.10E-15 | 9.44E-14 | NA |
| CD19 | 2.320526 | 3.17E-09 | 3.86E-08 | CD19 molecule |
| TPSD1 | 2.319625 | 1.94E-06 | 1.10E-05 | tryptase delta 1 |
| KCNA3 | 2.306475 | 1.85E-11 | 4.28E-10 | potassium voltage-gated channel subfamily A member 3 |
| C11orf21 | 2.303748 | 1.54E-12 | 4.89E-11 | chromosome 11 open reading frame 21 |
| CARD11 | 2.302853 | 1.05E-16 | 1.29E-14 | caspase recruitment domain family member 11 |
| IL2RG | 2.294307 | 1.18E-13 | 5.24E-12 | interleukin 2 receptor subunit gamma |
| TMC8 | 2.289608 | 4.82E-13 | 1.78E-11 | transmembrane channel like 8 |
| WNT10A | 2.286045 | 6.44E-12 | 1.71E-10 | Wnt family member 10A |
| WNT7B | 2.283838 | 5.11E-06 | 2.57E-05 | Wnt family member 7B |
| FCMR | 2.277232 | 3.55E-09 | 4.27E-08 | Fc fragment of IgM receptor |
| TPSB2 | 2.275901 | 7.79E-09 | 8.42E-08 | tryptase alpha/beta 1 |
| CRLF1 | 2.271871 | 4.15E-11 | 8.69E-10 | cytokine receptor like factor 1 |
| TNFSF14 | 2.268629 | 8.82E-16 | 7.86E-14 | TNF superfamily member 14 |

| | | | | |
|---|---|---|---|---|
| TLR10 | 2.266561 | 5.88E-09 | 6.59E-08 | toll like receptor 10 |
| TESPA1 | 2.261263 | 3.20E-12 | 9.33E-11 | thymocyte expressed, positive selection associated 1 |
| GRAP2 | 2.260505 | 4.88E-13 | 1.79E-11 | GRB2 related adaptor protein 2 |
| MSC | 2.259517 | 2.66E-21 | 1.38E-18 | musculin |
| CUX2 | 2.25043 | 5.07E-09 | 5.80E-08 | cut like homeobox 2 |
| PNOC | 2.249918 | 2.45E-10 | 4.12E-09 | prepronociceptin |
| HIST1H3J | 2.245874 | 1.78E-07 | 1.34E-06 | H3 clustered histone 12 |
| LY9 | 2.245749 | 1.78E-11 | 4.13E-10 | lymphocyte antigen 9 |
| HIST1H1B | 2.241786 | 8.11E-13 | 2.84E-11 | H1.5 linker histone, cluster member |
| CCR7 | 2.241764 | 6.64E-10 | 9.84E-09 | C-C motif chemokine receptor 7 |
| TIGIT | 2.239327 | 1.24E-12 | 4.08E-11 | T cell immunoreceptor with Ig and ITIM domains |
| KCNQ2 | 2.238222 | 0.001663 | 0.004215 | potassium voltage-gated channel subfamily Q member 2 |
| SELL | 2.234363 | 8.59E-11 | 1.64E-09 | selectin L |
| CPA4 | 2.231431 | 4.74E-06 | 2.41E-05 | carboxypeptidase A4 |
| SLAMF1 | 2.22948 | 1.76E-13 | 7.50E-12 | signaling lymphocytic activation molecule family member 1 |
| RAC2 | 2.228643 | 4.09E-16 | 4.04E-14 | Rac family small GTPase 2 |
| FAM159A | 2.222273 | 1.56E-09 | 2.06E-08 | shisa like 2A |
| WNT7A | 2.221857 | 6.63E-09 | 7.30E-08 | Wnt family member 7A |
| PRKCB | 2.219123 | 1.22E-13 | 5.38E-12 | protein kinase C beta |
| LOC105370697 | 2.21798 | 7.27E-05 | 0.000267 | uncharacterized LOC105370697 |
| WISP2 | 2.211358 | 3.18E-07 | 2.22E-06 | cellular communication network factor 5 |
| CD6 | 2.209738 | 6.67E-12 | 1.75E-10 | CD6 molecule |
| C1S | 2.206268 | 1.03E-26 | 1.61E-23 | complement C1s |
| ADCYAP1R1 | 2.205051 | 1.06E-06 | 6.44E-06 | ADCYAP receptor type I |
| JSRP1 | 2.2009 | 2.52E-05 | 0.000106 | junctional sarcoplasmic reticulum protein 1 |
| JCHAIN | 2.197463 | 5.00E-06 | 2.52E-05 | joining chain of multimeric IgA and IgM |
| THBS2 | 2.193199 | 2.21E-15 | 1.67E-13 | thrombospondin 2 |
| ITGBL1 | 2.191434 | 1.04E-14 | 6.37E-13 | integrin subunit beta like 1 |
| THEMIS | 2.188934 | 1.45E-14 | 8.55E-13 | thymocyte selection associated |
| MS4A6A | 2.187736 | 2.26E-24 | 2.43E-21 | membrane spanning 4-domains A6A |
| LOC100129697 | 2.183769 | 4.53E-11 | 9.38E-10 | uncharacterized LOC100129697 |
| GDF5 | 2.183631 | 3.93E-06 | 2.04E-05 | growth differentiation factor 5 |
| CCL22 | 2.179564 | 1.83E-06 | 1.04E-05 | C-C motif chemokine ligand 22 |
| SYT12 | 2.17673 | 1.32E-10 | 2.38E-09 | synaptotagmin 11 |
| CD3G | 2.175834 | 1.21E-15 | 1.02E-13 | CD3g molecule |

| | | | | |
|---|---|---|---|---|
| POM121L9P | 2.173638 | 2.07E-09 | 2.67E-08 | POM121 transmembrane nucleoporin like 9, pseudogene |
| MYBL2 | 2.173579 | 1.28E-10 | 2.33E-09 | MYB proto-oncogene like 2 |
| GPR55 | 2.173025 | 5.15E-07 | 3.42E-06 | G protein-coupled receptor 55 |
| AHNAK2 | 2.170377 | 1.22E-17 | 2.10E-15 | AHNAK nucleoprotein 2 |
| GALNT5 | 2.170282 | 6.20E-11 | 1.23E-09 | polypeptide N-acetylgalactosaminyltransferase 5 |
| LOC101926964 | 2.163528 | 3.54E-09 | 4.25E-08 | uncharacterized LOC101926964 |
| CARMIL2 | 2.159721 | 3.20E-12 | 9.33E-11 | capping protein regulator and myosin 1 linker 2 |
| GZMB | 2.157508 | 2.44E-06 | 1.34E-05 | granzyme B |
| S1PR4 | 2.157273 | 2.88E-13 | 1.14E-11 | sphingosine-1-phosphate receptor 4 |
| TTC24 | 2.152569 | 1.40E-06 | 8.24E-06 | tetratricopeptide repeat domain 24 |
| LIX1 | 2.151189 | 1.21E-13 | 5.37E-12 | limb and CNS expressed 1 |
| DOCK2 | 2.147709 | 6.91E-16 | 6.45E-14 | dedicator of cytokinesis 2 |
| SAA2 | 2.147617 | 0.00036 | 0.0011 | serum amyloid A1 |
| SCML4 | 2.139608 | 1.57E-13 | 6.80E-12 | Scm polycomb group protein like 4 |
| CD244 | 2.137575 | 8.69E-14 | 4.06E-12 | CD244 molecule |
| CLEC12A | 2.137305 | 1.25E-09 | 1.70E-08 | C-type lectin domain family 12 member A |
| CCL13 | 2.135858 | 1.90E-07 | 1.41E-06 | C-C motif chemokine ligand 13 |
| MIR650 | 2.135635 | 0.000973 | 0.00264 | microRNA 650 |
| HIST1H2AM | 2.135528 | 6.34E-12 | 1.69E-10 | H2A clustered histone 17 |
| MS4A2 | 2.13267 | 8.10E-08 | 6.71E-07 | membrane spanning 4-domains A1 |
| LCP1 | 2.132478 | 5.34E-15 | 3.72E-13 | lymphocyte cytosolic protein 1 |
| ITGAL | 2.128435 | 6.01E-14 | 3.00E-12 | integrin subunit alpha L |
| ZBP1 | 2.12495 | 9.90E-10 | 1.39E-08 | insulin like growth factor 2 mRNA binding protein 1 |
| CCR5 | 2.123949 | 1.05E-14 | 6.38E-13 | C-C motif chemokine receptor 5 |
| SLAMF6 | 2.123442 | 1.55E-12 | 4.91E-11 | SLAM family member 6 |
| HIST1H2AI | 2.123199 | 3.23E-09 | 3.93E-08 | H2A clustered histone 13 |
| MYO1G | 2.12231 | 1.33E-14 | 7.92E-13 | myosin IG |
| FCGR2B | 2.121183 | 3.41E-13 | 1.32E-11 | Fc fragment of IgG receptor IIb |
| MIR142 | 2.120985 | 2.67E-08 | 2.51E-07 | microRNA 142 |
| SASH3 | 2.117715 | 1.58E-17 | 2.63E-15 | SAM and SH3 domain containing 3 |
| MAP1LC3C | 2.116373 | 5.45E-07 | 3.59E-06 | microtubule associated protein 1 light chain 3 gamma |
| BCL11A | 2.114753 | 4.06E-09 | 4.76E-08 | BAF chromatin remodeling complex subunit BCL11A |
| ARHGAP40 | 2.112162 | 6.47E-11 | 1.28E-09 | Rho GTPase activating protein 40 |

| | | | | |
|---|---|---|---|---|
| ITK | 2.109448 | 1.60E-11 | 3.75E-10 | IL2 inducible T cell kinase |
| HIST1H2AG | 2.108774 | 2.54E-12 | 7.63E-11 | H2A clustered histone 11 |
| BTK | 2.103574 | 5.02E-16 | 4.85E-14 | Bruton tyrosine kinase |
| ARHGAP9 | 2.102266 | 2.40E-13 | 9.73E-12 | Rho GTPase activating protein 9 |
| PIK3R6 | 2.100464 | 2.55E-13 | 1.02E-11 | phosphoinositide-3-kinase regulatory subunit 6 |
| NCMAP | 2.092866 | 2.71E-13 | 1.08E-11 | non-compact myelin associated protein |
| CLEC17A | 2.092242 | 2.85E-06 | 1.54E-05 | C-type lectin domain containing 17A |
| IFNG_AS1 | 2.088266 | 5.35E-07 | 3.54E-06 | NA |
| CAPN6 | 2.087012 | 2.14E-15 | 1.62E-13 | calpain 6 |
| GZMM | 2.08622 | 7.44E-13 | 2.62E-11 | granzyme M |
| CP | 2.081584 | 3.77E-19 | 1.08E-16 | ceruloplasmin |
| XCL2 | 2.075665 | 1.23E-06 | 7.32E-06 | X-C motif chemokine ligand 2 |
| MYBPC1 | 2.0752 | 0.004676 | 0.010511 | myosin binding protein C1 |
| CACNA1I | 2.068955 | 5.10E-05 | 0.000196 | calcium voltage-gated channel subunit alpha1 I |
| CD1D | 2.067168 | 6.75E-13 | 2.40E-11 | CD1d molecule |
| FGD2 | 2.066768 | 6.95E-16 | 6.46E-14 | melanocortin 2 receptor accessory protein |
| CD53 | 2.059395 | 3.48E-15 | 2.51E-13 | CD53 molecule |
| LIMD2 | 2.058725 | 9.90E-17 | 1.22E-14 | LIM domain containing 2 |
| LAMC2 | 2.058456 | 3.96E-13 | 1.51E-11 | laminin subunit gamma 2 |
| HIST1H3G | 2.057463 | 1.10E-08 | 1.14E-07 | H3 clustered histone 8 |
| FAIM2 | 2.05268 | 2.15E-13 | 8.96E-12 | Fas apoptotic inhibitory molecule 2 |
| DAPP1 | 2.043455 | 5.56E-12 | 1.51E-10 | dual adaptor of phosphotyrosine and 3-phosphoinositides 1 |
| PRAMENP | 2.039535 | 0.000875 | 0.002408 | PRAME N-terminal like, pseudogene |
| SPN | 2.039153 | 7.45E-12 | 1.95E-10 | sialophorin |
| FCHO1 | 2.036633 | 2.86E-12 | 8.51E-11 | FCH and mu domain containing endocytic adaptor 1 |
| ALOX15 | 2.03619 | 0.000308 | 0.000957 | arachidonate 15-lipoxygenase |
| CD247 | 2.034557 | 7.00E-15 | 4.66E-13 | CD247 molecule |
| HIST1H2AH | 2.028669 | 3.73E-09 | 4.43E-08 | H2A clustered histone 12 |
| C1QA | 2.026956 | 6.70E-18 | 1.31E-15 | complement C1q A chain |
| GZMA | 2.024779 | 9.98E-14 | 4.56E-12 | granzyme A |
| PARVG | 2.024266 | 3.72E-15 | 2.66E-13 | parvin gamma |
| CD96 | 2.021724 | 2.68E-11 | 5.89E-10 | CD96 molecule |
| PLCB2 | 2.018041 | 6.97E-15 | 4.66E-13 | phospholipase C beta 2 |
| PIK3R5 | 2.018036 | 6.84E-13 | 2.43E-11 | phosphoinositide-3-kinase regulatory subunit 5 |
| IGDCC4 | 2.017409 | 1.03E-08 | 1.08E-07 | immunoglobulin superfamily DCC subclass member 4 |

| | | | | |
|---|---|---|---|---|
| CD84 | 2.014167 | 3.44E-17 | 4.97E-15 | CD84 molecule |
| COL5A1 | 2.01238 | 3.73E-13 | 1.43E-11 | collagen type V alpha 1 chain |
| RARRES1 | 2.010926 | 1.09E-12 | 3.66E-11 | retinoic acid receptor responder 1 |
| AGAP2 | 2.010576 | 1.52E-12 | 4.87E-11 | ArfGAP with GTPase domain, ankyrin repeat and PH domain 2 |
| SH2D1A | 2.009641 | 1.20E-09 | 1.64E-08 | SH2 domain containing 1A |
| DLGAP1 | 2.007671 | 1.41E-12 | 4.54E-11 | DLG associated protein 1 |
| NCKAP1L | 2.006692 | 3.39E-16 | 3.43E-14 | NCK associated protein 1 like |
| CST1 | 2.005397 | 0.007533 | 0.016044 | cystatin SN |
| ALX1 | 2.003859 | 1.56E-12 | 4.93E-11 | ALX homeobox 1 |
| CHIT1 | 2.001069 | 0.00031 | 0.000962 | chitinase 1 |
| ALOX5 | 1.991771 | 8.44E-16 | 7.64E-14 | arachidonate 5-lipoxygenase |
| TYROBP | 1.986824 | 8.70E-16 | 7.79E-14 | transmembrane immune signaling adaptor TYROBP |
| C3orf80 | 1.985073 | 9.54E-08 | 7.72E-07 | chromosome 3 open reading frame 80 |
| CD70 | 1.984578 | 1.44E-06 | 8.45E-06 | CD70 molecule |
| TMSB10 | 1.984566 | 2.13E-19 | 6.65E-17 | thymosin beta 10 |
| SLCO5A1 | 1.981611 | 8.52E-11 | 1.64E-09 | solute carrier organic anion transporter family member 5A1 |
| DEF6 | 1.977905 | 4.84E-16 | 4.72E-14 | defensin alpha 6 |
| JAK3 | 1.977076 | 2.60E-11 | 5.78E-10 | Janus kinase 3 |
| APCDD1 | 1.975864 | 2.89E-16 | 2.95E-14 | APC down-regulated 1 |
| HIST1H1D | 1.96874 | 7.47E-11 | 1.46E-09 | H1.3 linker histone, cluster member |
| ADRA2A | 1.968476 | 1.03E-11 | 2.57E-10 | adrenoceptor alpha 2A |
| CCR4 | 1.96825 | 4.32E-12 | 1.21E-10 | C-C motif chemokine receptor 4 |
| CSF2RB | 1.967246 | 5.43E-11 | 1.10E-09 | colony stimulating factor 2 receptor subunit beta |
| CXCR3 | 1.966958 | 3.07E-15 | 2.25E-13 | C-X-C motif chemokine receptor 3 |
| KCNS1 | 1.965294 | 9.61E-14 | 4.44E-12 | potassium voltage-gated channel modifier subfamily S member 1 |
| CD200R1 | 1.964254 | 5.07E-13 | 1.86E-11 | CD200 receptor 1 |
| P2RY10 | 1.960908 | 3.81E-08 | 3.44E-07 | P2Y receptor family member 10 |
| CD8A | 1.956539 | 2.57E-10 | 4.32E-09 | CD8a molecule |
| FNDC4 | 1.956151 | 1.06E-13 | 4.79E-12 | fibronectin type III domain containing 4 |
| IL10RA | 1.955579 | 4.70E-13 | 1.75E-11 | interleukin 10 receptor subunit alpha |
| GAPT | 1.95163 | 1.51E-09 | 2.00E-08 | GRB2 binding adaptor protein, transmembrane |
| CD163L1 | 1.950672 | 3.89E-11 | 8.21E-10 | CD163 molecule like 1 |

| | | | | |
|---|---|---|---|---|
| ARL11 | 1.948912 | 1.03E-14 | 6.35E-13 | ADP ribosylation factor like GTPase 11 |
| PTPRC | 1.94774 | 4.22E-12 | 1.18E-10 | protein tyrosine phosphatase receptor type C |
| MAP4K1 | 1.947602 | 8.84E-13 | 3.06E-11 | mitogen-activated protein kinase kinase kinase kinase 1 |
| MDK | 1.938251 | 9.61E-19 | 2.43E-16 | midkine |
| CXCR2P1 | 1.937994 | 0.000236 | 0.000755 | C-X-C motif chemokine receptor 2 pseudogene 1 |
| GABRP | 1.93482 | 2.56E-08 | 2.42E-07 | gamma-aminobutyric acid type A receptor subunit pi |
| IRF8 | 1.931957 | 2.31E-12 | 7.00E-11 | interferon regulatory factor 8 |
| TNFRSF18 | 1.928325 | 3.22E-06 | 1.71E-05 | TNF receptor superfamily member 18 |
| C10orf128 | 1.925483 | 1.01E-19 | 3.56E-17 | transmembrane protein 273 |
| RASSF2 | 1.92315 | 1.43E-13 | 6.24E-12 | Ras association domain family member 2 |
| LINC01279 | 1.921653 | 7.82E-15 | 5.09E-13 | coiled-coil domain containing 80 |
| VAV1 | 1.92123 | 5.29E-15 | 3.69E-13 | vav guanine nucleotide exchange factor 1 |
| ADAMDEC1 | 1.918951 | 0.000601 | 0.001726 | ADAM like decysin 1 |
| CCND2 | 1.917927 | 1.72E-20 | 7.57E-18 | cyclin D2 |
| LAIR1 | 1.91774 | 1.50E-18 | 3.58E-16 | leukocyte associated immunoglobulin like receptor 1 |
| HIST1H2BH | 1.912388 | 1.72E-10 | 3.02E-09 | H2B clustered histone 9 |
| SLA2 | 1.91043 | 6.36E-14 | 3.15E-12 | Src like adaptor 2 |
| COL14A1 | 1.904741 | 1.30E-16 | 1.51E-14 | collagen type XIV alpha 1 chain |
| HCST | 1.904489 | 8.23E-12 | 2.10E-10 | hematopoietic cell signal transducer |
| ARNTL2 | 1.902258 | 3.26E-13 | 1.27E-11 | aryl hydrocarbon receptor nuclear translocator like 2 |
| C1R | 1.9006 | 9.70E-22 | 5.75E-19 | complement C1r |
| E2F2 | 1.899263 | 6.86E-06 | 3.34E-05 | E2F transcription factor 2 |
| VWF | 1.897756 | 7.80E-12 | 2.01E-10 | von Willebrand factor |
| CCDC88B | 1.896364 | 1.93E-11 | 4.42E-10 | coiled-coil domain containing 88B |
| ARHGAP30 | 1.894436 | 5.41E-14 | 2.73E-12 | Rho GTPase activating protein 30 |
| BCL11B | 1.893809 | 1.53E-12 | 4.88E-11 | BAF chromatin remodeling complex subunit BCL11B |
| PYCARD | 1.892731 | 3.53E-16 | 3.53E-14 | PYD and CARD domain containing |
| CTSS | 1.891711 | 4.06E-15 | 2.86E-13 | cathepsin S |
| ZAP70 | 1.889351 | 6.41E-11 | 1.27E-09 | zeta chain of T cell receptor associated protein kinase 70 |

| | | | | |
|---|---|---|---|---|
| TNFRSF13B | 1.888201 | 0.000572 | 0.001652 | TNF receptor superfamily member 13B |
| RASAL3 | 1.88716 | 1.15E-12 | 3.83E-11 | RAS protein activator like 3 |
| ISLR2 | 1.886916 | 4.28E-09 | 4.98E-08 | immunoglobulin superfamily containing leucine rich repeat 2 |
| CD209 | 1.886314 | 2.38E-12 | 7.17E-11 | CD209 molecule |
| COL1A2 | 1.884279 | 4.05E-18 | 8.58E-16 | collagen type I alpha 2 chain |
| INHBA | 1.884073 | 2.17E-10 | 3.73E-09 | inhibin subunit beta A |
| PVT1 | 1.884009 | 3.09E-17 | 4.50E-15 | Pvt1 oncogene |
| KIF21B | 1.881928 | 9.70E-13 | 3.30E-11 | kinesin family member 21B |
| FAM78A | 1.876093 | 7.82E-17 | 9.95E-15 | family with sequence similarity 78 member A |
| CCL5 | 1.874313 | 1.77E-10 | 3.11E-09 | C-C motif chemokine ligand 5 |
| GLIPR2 | 1.868567 | 1.58E-21 | 8.78E-19 | GLI pathogenesis related 2 |
| FCN1 | 1.867434 | 5.13E-07 | 3.40E-06 | ficolin 1 |
| AIM2 | 1.864595 | 1.67E-05 | 7.34E-05 | absent in melanoma 2 |
| NCF4 | 1.863747 | 3.97E-18 | 8.58E-16 | neutrophil cytosolic factor 4 |
| PTGDR | 1.862414 | 8.70E-12 | 2.21E-10 | prostaglandin D2 receptor |
| MPEG1 | 1.860788 | 7.71E-12 | 2.00E-10 | macrophage expressed 1 |
| NOD2 | 1.859823 | 2.05E-11 | 4.69E-10 | nucleotide binding oligomerization domain containing 2 |
| TMSB4X | 1.85448 | 1.34E-19 | 4.51E-17 | thymosin beta 4 X-linked |
| C1QC | 1.852623 | 5.29E-14 | 2.68E-12 | complement C1q C chain |
| SELPLG | 1.851797 | 1.13E-14 | 6.83E-13 | selectin P ligand |
| NRSN1 | 1.849152 | 2.10E-11 | 4.76E-10 | neurensin 1 |
| ITGB2_AS1 | 1.846898 | 1.04E-11 | 2.60E-10 | NA |
| COL7A1 | 1.846728 | 2.02E-05 | 8.71E-05 | collagen type VII alpha 1 chain |
| PLEK | 1.841972 | 4.32E-12 | 1.21E-10 | pleckstrin |
| HIST1H3F | 1.834972 | 8.69E-10 | 1.25E-08 | H3 clustered histone 7 |
| LRG1 | 1.834634 | 2.19E-06 | 1.22E-05 | leucine rich alpha-2-glycoprotein 1 |
| CXCR5 | 1.833693 | 5.26E-08 | 4.58E-07 | C-X-C motif chemokine receptor 5 |
| TRG_AS1 | 1.83159 | 5.87E-10 | 8.85E-09 | NA |
| B3GALT5 | 1.831324 | 3.57E-11 | 7.56E-10 | beta-1,3-galactosyltransferase 5 |
| IL16 | 1.831193 | 1.06E-11 | 2.63E-10 | interleukin 16 |
| DOK2 | 1.830568 | 8.49E-15 | 5.40E-13 | docking protein 2 |
| FCRL6 | 1.829736 | 1.05E-09 | 1.46E-08 | Fc receptor like 6 |
| HIST1H3B | 1.827298 | 6.31E-10 | 9.43E-09 | H3 clustered histone 2 |
| RUNX3 | 1.827009 | 8.47E-10 | 1.22E-08 | RUNX family transcription factor 3 |
| CD28 | 1.825249 | 3.08E-11 | 6.63E-10 | CD28 molecule |
| PLD4 | 1.825159 | 1.40E-08 | 1.41E-07 | phospholipase D family member 4 |
| S100B | 1.824394 | 1.23E-06 | 7.35E-06 | S100 calcium binding protein B |

| | | | | |
|---|---|---|---|---|
| OSR2 | 1.82167 | 7.87E-09 | 8.48E-08 | odd-skipped related transciption factor 2 |
| TMEM59L | 1.819522 | 3.71E-09 | 4.41E-08 | transmembrane protein 59 like |
| CD40LG | 1.81833 | 1.22E-08 | 1.25E-07 | CD40 ligand |
| ITPRIPL1 | 1.817058 | 1.10E-17 | 1.97E-15 | ITPRIP like 1 |
| HSH2D | 1.813824 | 3.28E-08 | 3.00E-07 | hematopoietic SH2 domain containing |
| MKI67 | 1.813035 | 9.95E-10 | 1.40E-08 | marker of proliferation Ki-67 |
| DERL3 | 1.810062 | 2.40E-08 | 2.29E-07 | derlin 3 |
| CFP | 1.805984 | 7.27E-10 | 1.06E-08 | complement factor properdin |
| SVEP1 | 1.803605 | 9.77E-27 | 1.61E-23 | sushi, von Willebrand factor type A, EGF and pentraxin domain containing 1 |
| CD300LF | 1.803298 | 4.63E-11 | 9.56E-10 | CD300 molecule like family member f |
| NIPAL4 | 1.801941 | 7.76E-13 | 2.73E-11 | NIPA like domain containing 4 |
| PARP15 | 1.798443 | 1.14E-08 | 1.18E-07 | poly(ADP-ribose) polymerase family member 15 |
| HIST2H3D | 1.795924 | 1.58E-11 | 3.71E-10 | H3 clustered histone 13 |
| CD33 | 1.793261 | 6.64E-16 | 6.23E-14 | CD33 molecule |
| MGP | 1.792855 | 2.64E-15 | 1.96E-13 | matrix Gla protein |
| LAPTM5 | 1.790942 | 2.02E-13 | 8.43E-12 | lysosomal protein transmembrane 5 |
| CLIC6 | 1.784509 | 2.21E-09 | 2.81E-08 | chloride intracellular channel 6 |
| ARHGAP15 | 1.782873 | 1.82E-15 | 1.43E-13 | Rho GTPase activating protein 15 |
| LAMP5 | 1.78195 | 1.14E-07 | 9.07E-07 | lysosomal associated membrane protein family member 5 |
| NKG7 | 1.781278 | 6.62E-09 | 7.30E-08 | natural killer cell granule protein 7 |
| MDFI | 1.780792 | 1.11E-14 | 6.73E-13 | MyoD family inhibitor |
| STEAP4 | 1.778349 | 1.91E-13 | 8.11E-12 | STEAP4 metalloreductase |
| CCDC80 | 1.776061 | 1.70E-16 | 1.91E-14 | coiled-coil domain containing 80 |
| C1QB | 1.77551 | 7.75E-12 | 2.00E-10 | complement C1q B chain |
| JAKMIP1 | 1.775191 | 1.65E-07 | 1.25E-06 | janus kinase and microtubule interacting protein 1 |
| CD27 | 1.771567 | 3.56E-10 | 5.74E-09 | CD27 molecule |
| GFRA2 | 1.770387 | 9.35E-19 | 2.43E-16 | GDNF family receptor alpha 2 |
| GRIN2A | 1.769249 | 1.11E-11 | 2.72E-10 | glutamate ionotropic receptor NMDA type subunit 2A |
| HIST1H2BO | 1.768669 | 1.31E-05 | 5.94E-05 | H2B clustered histone 17 |
| LINC00861 | 1.767158 | 2.70E-10 | 4.49E-09 | long intergenic non-protein coding RNA 861 |
| HIST1H4F | 1.766445 | 2.11E-09 | 2.70E-08 | H4 clustered histone 6 |
| DGKA | 1.758877 | 8.20E-11 | 1.58E-09 | diacylglycerol kinase alpha |
| AKNA | 1.753808 | 4.70E-11 | 9.69E-10 | AT-hook transcription factor |

| | | | | |
|---|---|---|---|---|
| MMRN1 | 1.751617 | 1.97E-05 | 8.49E-05 | multimerin 1 |
| KLRB1 | 1.750492 | 2.82E-08 | 2.62E-07 | killer cell lectin like receptor B1 |
| CRTAM | 1.744516 | 1.15E-07 | 9.13E-07 | cytotoxic and regulatory T cell molecule |
| APOC1 | 1.743757 | 2.53E-05 | 0.000106 | apolipoprotein C1 |
| FLJ16779 | 1.742603 | 1.16E-06 | 7.00E-06 | uncharacterized LOC100192386 |
| MEGF11 | 1.738439 | 2.19E-11 | 4.96E-10 | multiple EGF like domains 11 |
| FASLG | 1.736599 | 2.34E-09 | 2.96E-08 | Fas ligand |
| ITGAX | 1.73555 | 1.14E-07 | 9.07E-07 | integrin subunit alpha X |
| LILRB1 | 1.735338 | 2.01E-12 | 6.14E-11 | leukocyte immunoglobulin like receptor B1 |
| SIGLEC1 | 1.7351 | 1.10E-08 | 1.15E-07 | sialic acid binding Ig like lectin 1 |
| CYBB | 1.729594 | 8.34E-15 | 5.32E-13 | cytochrome b-245 beta chain |
| NCF2 | 1.729348 | 2.20E-12 | 6.68E-11 | neutrophil cytosolic factor 2 |
| COMP | 1.729151 | 4.83E-10 | 7.49E-09 | cartilage oligomeric matrix protein |
| CEACAM21 | 1.728401 | 1.38E-12 | 4.47E-11 | CEA cell adhesion molecule 21 |
| ADCY7 | 1.727497 | 5.92E-13 | 2.15E-11 | adenylate cyclase 7 |
| NAPSB | 1.724855 | 1.94E-12 | 5.99E-11 | napsin B aspartic peptidase, pseudogene |
| LOC101927751 | 1.723645 | 5.38E-19 | 1.49E-16 | uncharacterized LOC101927751 |
| TMEM179 | 1.722104 | 4.18E-06 | 2.16E-05 | transmembrane protein 179 |
| LUM | 1.721879 | 1.17E-20 | 5.44E-18 | lumican |
| TXLNB | 1.718018 | 1.65E-05 | 7.27E-05 | taxilin beta |
| CCDC141 | 1.717972 | 2.41E-13 | 9.75E-12 | coiled-coil domain containing 141 |
| LINC00639 | 1.717937 | 1.87E-08 | 1.83E-07 | long intergenic non-protein coding RNA 639 |
| RNASE6 | 1.71749 | 2.97E-13 | 1.17E-11 | ribonuclease A family member k6 |
| TNFRSF13C | 1.714939 | 3.49E-05 | 0.00014 | TNF receptor superfamily member 13C |
| MYH7 | 1.713586 | 0.001567 | 0.003999 | myosin heavy chain 7 |
| MYCL | 1.713148 | 3.86E-10 | 6.15E-09 | MYCL proto-oncogene, bHLH transcription factor |
| NCF1 | 1.710304 | 2.32E-09 | 2.94E-08 | neutrophil cytosolic factor 1 |
| HCK | 1.708818 | 2.30E-13 | 9.45E-12 | HCK proto-oncogene, Src family tyrosine kinase |
| RGS19 | 1.706666 | 1.20E-17 | 2.08E-15 | regulator of G protein signaling 19 |
| EPHB2 | 1.704967 | 3.73E-09 | 4.43E-08 | EPH receptor B2 |
| BIN2 | 1.704645 | 1.03E-11 | 2.57E-10 | bridging integrator 2 |
| SLC24A4 | 1.699943 | 2.14E-09 | 2.74E-08 | solute carrier family 24 member 4 |

| | | | | |
|---|---|---|---|---|
| CXCL13 | 1.699928 | 0.023256 | 0.042691 | C-X-C motif chemokine ligand 13 |
| HAVCR1 | 1.699436 | 9.02E-08 | 7.35E-07 | hepatitis A virus cellular receptor 1 |
| WNT4 | 1.695899 | 4.45E-12 | 1.24E-10 | Wnt family member 4 |
| CD300A | 1.692277 | 1.18E-10 | 2.17E-09 | CD300a molecule |
| IL24 | 1.688098 | 0.001122 | 0.002988 | interleukin 24 |
| HBB | 1.687688 | 0.003499 | 0.008145 | hemoglobin subunit beta |
| MS4A4A | 1.687542 | 2.18E-13 | 9.01E-12 | membrane spanning 4-domains A4A |
| IL12RB1 | 1.686877 | 1.58E-12 | 4.97E-11 | interleukin 12 receptor subunit beta 1 |
| GBP5 | 1.686635 | 4.37E-08 | 3.89E-07 | guanylate binding protein 5 |
| HTR7 | 1.68573 | 1.75E-08 | 1.73E-07 | 5-hydroxytryptamine receptor 7 |
| PLXNA4 | 1.685163 | 1.13E-13 | 5.07E-12 | plexin A4 |
| COL16A1 | 1.682798 | 2.73E-11 | 5.99E-10 | collagen type XVI alpha 1 chain |
| FOLR2 | 1.681881 | 2.13E-19 | 6.65E-17 | folate receptor beta |
| CAMK4 | 1.681161 | 4.52E-11 | 9.37E-10 | calcium/calmodulin dependent protein kinase IV |
| PPP1R1B | 1.680047 | 3.95E-10 | 6.27E-09 | protein phosphatase 1 regulatory inhibitor subunit 1B |
| SIGLEC12 | 1.679688 | 0.000328 | 0.001011 | sialic acid binding Ig like lectin 12 |
| CEACAM4 | 1.679686 | 2.14E-06 | 1.19E-05 | CEA cell adhesion molecule 4 |
| ADAM12 | 1.677485 | 2.21E-10 | 3.79E-09 | ADAM metallopeptidase domain 12 |
| TMEM92 | 1.675827 | 2.95E-05 | 0.000121 | transmembrane protein 92 |
| LAT2 | 1.674318 | 1.07E-15 | 9.20E-14 | linker for activation of T cells family member 2 |
| PIK3CG | 1.674249 | 9.45E-14 | 4.38E-12 | phosphatidylinositol-4,5-bisphosphate 3-kinase catalytic subunit gamma |
| GFRA1 | 1.670627 | 1.25E-14 | 7.50E-13 | GDNF family receptor alpha 1 |
| ADAM19 | 1.669787 | 2.06E-12 | 6.30E-11 | ADAM metallopeptidase domain 19 |
| GFI1 | 1.668256 | 5.47E-11 | 1.11E-09 | growth factor independent 1 transcriptional repressor |
| TLR8 | 1.659844 | 5.22E-09 | 5.94E-08 | toll like receptor 8 |
| TRAF3IP3 | 1.65977 | 8.92E-11 | 1.70E-09 | TRAF3 interacting protein 3 |
| FAM65B | 1.658868 | 8.23E-10 | 1.19E-08 | RHO family interacting cell polarization regulator 2 |
| PROM1 | 1.657952 | 6.11E-09 | 6.81E-08 | prominin 1 |
| TGFBI | 1.656764 | 3.67E-16 | 3.64E-14 | transforming growth factor beta induced |
| LAMP3 | 1.655204 | 8.58E-06 | 4.07E-05 | lysosomal associated membrane protein 3 |
| TMC3 | 1.652724 | 7.95E-08 | 6.61E-07 | transmembrane channel like 3 |

| | | | | |
|---|---|---|---|---|
| ARHGAP22 | 1.652716 | 2.78E-17 | 4.12E-15 | Rho GTPase activating protein 22 |
| COL8A2 | 1.652303 | 9.48E-15 | 5.88E-13 | collagen type VIII alpha 2 chain |
| HIST1H2AL | 1.651997 | 3.92E-05 | 0.000156 | H2A clustered histone 16 |
| ALDH1A3 | 1.651647 | 2.73E-18 | 6.26E-16 | aldehyde dehydrogenase 1 family member A3 |
| CIDEA | 1.651029 | 0.025313 | 0.045902 | cell death inducing DFFA like effector a |
| PODN | 1.646208 | 1.97E-08 | 1.91E-07 | podocan |
| APOBEC3G | 1.644781 | 1.08E-16 | 1.32E-14 | apolipoprotein B mRNA editing enzyme catalytic subunit 3G |
| POU2F2 | 1.644346 | 1.11E-09 | 1.53E-08 | POU class 2 homeobox 2 |
| FBN1 | 1.643019 | 1.69E-27 | 3.63E-24 | fibrillin 1 |
| GPR174 | 1.641328 | 6.97E-06 | 3.39E-05 | G protein-coupled receptor 174 |
| KCNN4 | 1.64115 | 6.37E-09 | 7.06E-08 | potassium calcium-activated channel subfamily N member 4 |
| RIMS1 | 1.640897 | 0.000818 | 0.00227 | regulating synaptic membrane exocytosis 1 |
| PLA2G7 | 1.640511 | 7.62E-06 | 3.66E-05 | phospholipase A2 group VII |
| BEND4 | 1.638782 | 4.48E-05 | 0.000175 | BEN domain containing 4 |
| HSPA7 | 1.637666 | 3.88E-09 | 4.58E-08 | heat shock protein family A (Hsp70) member 7 (pseudogene) |
| MFAP4 | 1.635827 | 1.79E-19 | 5.80E-17 | microfibril associated protein 4 |
| SPON2 | 1.633788 | 1.12E-15 | 9.62E-14 | spondin 2 |
| SLFN12L | 1.633586 | 1.61E-09 | 2.13E-08 | schlafen family member 12 like |
| TMEM154 | 1.631946 | 1.24E-08 | 1.26E-07 | transmembrane protein 154 |
| CSTA | 1.631726 | 0.000126 | 0.000434 | cystatin A |
| FCGR2C | 1.63172 | 2.84E-10 | 4.70E-09 | Fc fragment of IgG receptor IIb |
| CAPN8 | 1.631242 | 1.18E-06 | 7.09E-06 | calpain 8 |
| LRRN4 | 1.629364 | 3.06E-09 | 3.75E-08 | leucine rich repeats and calponin homology domain containing 4 |
| FOXJ1 | 1.626518 | 2.10E-10 | 3.61E-09 | forkhead box J1 |
| LRRK1 | 1.625951 | 8.60E-14 | 4.03E-12 | leucine rich repeat kinase 1 |
| HIST1H4I | 1.62131 | 4.60E-09 | 5.31E-08 | H4 clustered histone 9 |
| SCARNA21 | 1.621227 | 4.85E-10 | 7.52E-09 | small Cajal body-specific RNA 21 |
| RIMS2 | 1.620369 | 5.12E-07 | 3.40E-06 | regulating synaptic membrane exocytosis 2 |
| ARL4C | 1.613937 | 5.79E-08 | 4.98E-07 | ADP ribosylation factor like GTPase 4C |
| MYO1F | 1.612261 | 1.08E-09 | 1.50E-08 | myosin IF |
| SLC34A2 | 1.611466 | 9.70E-07 | 5.96E-06 | solute carrier family 34 member 2 |
| RUFY4 | 1.611355 | 0.002337 | 0.005706 | RUN and FYVE domain containing 4 |

| | | | | |
|---|---|---|---|---|
| PRDM8 | 1.611215 | 3.55E-12 | 1.01E-10 | PR/SET domain 8 |
| TREML1 | 1.610409 | 1.15E-05 | 5.27E-05 | triggering receptor expressed on myeloid cells like 1 |
| C10orf105 | 1.608917 | 5.71E-10 | 8.65E-09 | chromosome 10 open reading frame 105 |
| AMPD1 | 1.608543 | 0.012094 | 0.02413 | adenosine monophosphate deaminase 1 |
| ADAMTS2 | 1.60687 | 8.38E-13 | 2.91E-11 | ADAM metallopeptidase with thrombospondin type 1 motif 2 |
| CILP | 1.606821 | 4.34E-08 | 3.87E-07 | cartilage intermediate layer protein |
| HLA_DPB1 | 1.604647 | 0.006525 | 0.014121 | NA |
| HAPLN3 | 1.60421 | 8.85E-10 | 1.26E-08 | hyaluronan and proteoglycan link protein 3 |
| GMIP | 1.602475 | 2.35E-11 | 5.25E-10 | GEM interacting protein |
| LILRA2 | 1.59974 | 7.56E-09 | 8.20E-08 | leukocyte immunoglobulin like receptor A2 |
| FMOD | 1.599163 | 5.05E-17 | 6.68E-15 | fibromodulin |
| ADGRE2 | 1.597787 | 1.68E-07 | 1.27E-06 | adhesion G protein-coupled receptor E2 |
| S100A4 | 1.595255 | 2.76E-15 | 2.03E-13 | S100 calcium binding protein A4 |
| PTAFR | 1.59301 | 4.61E-14 | 2.39E-12 | platelet activating factor receptor |
| XCR1 | 1.592996 | 1.34E-05 | 6.05E-05 | X-C motif chemokine receptor 1 |
| MIR31HG | 1.591104 | 8.22E-10 | 1.19E-08 | MIR31 host gene |
| UBE2C | 1.590547 | 8.54E-06 | 4.05E-05 | ubiquitin conjugating enzyme E2 C |
| CPXM1 | 1.590248 | 6.62E-06 | 3.24E-05 | carboxypeptidase X, M14 family member 1 |
| PLPP2 | 1.585529 | 7.56E-10 | 1.10E-08 | phospholipid phosphatase 2 |
| ZBED2 | 1.58327 | 3.70E-06 | 1.93E-05 | zinc finger BED-type containing 2 |
| PSTPIP1 | 1.579827 | 5.15E-09 | 5.87E-08 | proline-serine-threonine phosphatase interacting protein 1 |
| NFAM1 | 1.579393 | 1.33E-08 | 1.35E-07 | NFAT activating protein with ITAM motif 1 |
| SRPX | 1.578526 | 2.06E-15 | 1.58E-13 | sushi repeat containing protein X-linked |
| SHD | 1.577341 | 0.011138 | 0.022511 | Src homology 2 domain containing transforming protein D |
| RET | 1.575493 | 2.20E-12 | 6.68E-11 | ret proto-oncogene |
| VSIG4 | 1.574862 | 1.31E-15 | 1.09E-13 | V-set and immunoglobulin domain containing 4 |
| LINGO3 | 1.574491 | 4.77E-08 | 4.21E-07 | leucine rich repeat and Ig domain containing 3 |

| | | | | |
|---|---|---|---|---|
| QPCT | 1.574316 | 2.71E-14 | 1.52E-12 | glutaminyl-peptide cyclotransferase |
| GAS7 | 1.574273 | 2.18E-17 | 3.35E-15 | growth arrest specific 7 |
| TBX21 | 1.571829 | 3.73E-08 | 3.38E-07 | T-box transcription factor 21 |
| PYHIN1 | 1.571597 | 1.08E-06 | 6.59E-06 | pyrin and HIN domain family member 1 |
| IFI27L2 | 1.570472 | 4.98E-09 | 5.72E-08 | interferon alpha inducible protein 27 like 2 |
| ADGRE1 | 1.570383 | 2.23E-05 | 9.48E-05 | adhesion G protein-coupled receptor E1 |
| FAM131B | 1.569303 | 4.39E-08 | 3.91E-07 | family with sequence similarity 131 member B |
| PAPPA2 | 1.565907 | 3.00E-10 | 4.94E-09 | pappalysin 2 |
| FBXO41 | 1.560137 | 1.96E-13 | 8.28E-12 | F-box protein 41 |
| HAPLN1 | 1.559688 | 0.018258 | 0.034523 | hyaluronan and proteoglycan link protein 1 |
| KRT80 | 1.558233 | 3.53E-06 | 1.86E-05 | keratin 80 |
| RUNX1 | 1.557211 | 8.43E-09 | 9.02E-08 | RUNX family transcription factor 1 |
| HDC | 1.55367 | 2.17E-05 | 9.27E-05 | histidine decarboxylase |
| RUBCNL | 1.552124 | 6.14E-11 | 1.22E-09 | rubicon like autophagy enhancer |
| SPON1 | 1.55195 | 4.15E-15 | 2.91E-13 | spondin 1 |
| TLR6 | 1.551632 | 6.18E-12 | 1.66E-10 | toll like receptor 6 |
| SH3BP1 | 1.550876 | 1.26E-10 | 2.30E-09 | SH3 domain binding protein 1 |
| CSF1R | 1.549537 | 8.28E-14 | 3.91E-12 | colony stimulating factor 1 receptor |
| HIST1H2AJ | 1.547958 | 0.000219 | 0.000706 | H2A clustered histone 14 |
| FYB | 1.547596 | 2.25E-10 | 3.85E-09 | FYN binding protein 1 |
| TESC | 1.547476 | 2.40E-12 | 7.22E-11 | tescalcin |
| CRACR2A | 1.547223 | 8.34E-14 | 3.93E-12 | calcium release activated channel regulator 2A |
| BTLA | 1.547147 | 8.81E-07 | 5.50E-06 | B and T lymphocyte associated |
| MEI1 | 1.54673 | 1.12E-06 | 6.80E-06 | meiotic double-stranded break formation protein 1 |
| TREM2 | 1.546157 | 4.44E-07 | 2.99E-06 | triggering receptor expressed on myeloid cells 2 |
| PI15 | 1.545732 | 0.00051 | 0.001488 | peptidase inhibitor 15 |
| ACTG2 | 1.545425 | 0.005714 | 0.012581 | actin gamma 2, smooth muscle |
| AEBP1 | 1.544522 | 6.67E-12 | 1.75E-10 | AE binding protein 1 |
| EEF1A2 | 1.544459 | 0.011281 | 0.022758 | eukaryotic translation elongation factor 1 alpha 2 |
| HIST1H2BI | 1.54439 | 6.57E-09 | 7.25E-08 | H2B clustered histone 5 |
| SAMHD1 | 1.543951 | 2.39E-11 | 5.33E-10 | SAM and HD domain containing deoxynucleoside triphosphate triphosphohydrolase 1 |
| TMEM156 | 1.541952 | 1.27E-05 | 5.78E-05 | transmembrane protein 156 |

| | | | | |
|---|---|---|---|---|
| ADRB2 | 1.541162 | 3.74E-13 | 1.43E-11 | adrenoceptor beta 2 |
| HS3ST1 | 1.541041 | 4.06E-08 | 3.64E-07 | heparan sulfate-glucosamine 3-sulfotransferase 1 |
| TAGAP | 1.534237 | 6.16E-09 | 6.85E-08 | T cell activation RhoGTPase activating protein |
| PTPN22 | 1.532972 | 9.93E-09 | 1.05E-07 | protein tyrosine phosphatase non-receptor type 22 |
| VIM | 1.532815 | 3.00E-18 | 6.79E-16 | vimentin |
| ISLR | 1.531947 | 6.89E-15 | 4.64E-13 | immunoglobulin superfamily containing leucine rich repeat |
| CD22 | 1.529744 | 2.20E-06 | 1.22E-05 | CD22 molecule |
| TLR7 | 1.525888 | 7.18E-10 | 1.05E-08 | toll like receptor 7 |
| LOC100506585 | 1.524979 | 9.03E-10 | 1.28E-08 | NA |
| ADH1B | 1.524733 | 8.94E-13 | 3.09E-11 | alcohol dehydrogenase 1B (class I), beta polypeptide |
| TSHZ2 | 1.523795 | 5.13E-18 | 1.06E-15 | teashirt zinc finger homeobox 2 |
| TNNT1 | 1.520409 | 0.007226 | 0.015473 | troponin T1, slow skeletal type |
| CCL11 | 1.517901 | 2.14E-06 | 1.19E-05 | C-C motif chemokine ligand 11 |
| MUC12 | 1.513453 | 6.68E-07 | 4.30E-06 | mucin 12, cell surface associated |
| IL2RA | 1.513303 | 2.18E-06 | 1.21E-05 | interleukin 2 receptor subunit alpha |
| FERMT3 | 1.512367 | 3.97E-12 | 1.12E-10 | fermitin family member 3 |
| COL8A1 | 1.511667 | 1.98E-13 | 8.31E-12 | collagen type VIII alpha 1 chain |
| ITGB2 | 1.510952 | 1.85E-09 | 2.41E-08 | integrin subunit beta 2 |
| FN1 | 1.509097 | 5.06E-10 | 7.79E-09 | fibronectin 1 |
| LINC00426 | 1.504392 | 1.05E-09 | 1.46E-08 | long intergenic non-protein coding RNA 426 |
| HP | 1.503778 | 8.11E-05 | 0.000294 | haptoglobin |
| ICOS | 1.503616 | 0.000133 | 0.000455 | inducible T cell costimulator |
| ADD2 | 1.502032 | 9.23E-06 | 4.34E-05 | adducin 2 |
| GXYLT2 | 1.500192 | 1.78E-15 | 1.41E-13 | glucoside xylosyltransferase 2 |
| CNTNAP4 | -1.50098 | 0.002797 | 0.006681 | contactin associated protein family member 4 |
| ADAMTS19 | -1.50152 | 1.83E-05 | 7.98E-05 | ADAM metallopeptidase with thrombospondin type 1 motif 19 |
| LINC00645 | -1.5067 | 8.46E-11 | 1.63E-09 | long intergenic non-protein coding RNA 645 |
| LINC00970 | -1.51086 | 6.33E-09 | 7.02E-08 | long intergenic non-protein coding RNA 970 |
| DEPDC1B | -1.51373 | 4.87E-10 | 7.55E-09 | DEP domain containing 1B |
| CDH20 | -1.51738 | 4.67E-05 | 0.000181 | protocadherin gamma subfamily B, 4 |
| ALDH1L1_AS1 | -1.51801 | 6.50E-08 | 5.52E-07 | NA |
| APOB | -1.51859 | 2.38E-06 | 1.31E-05 | apolipoprotein B |

| | | | | |
|---|---|---|---|---|
| C2orf71 | -1.51932 | 0.000495 | 0.001452 | photoreceptor cilium actin regulator |
| PTGER3 | -1.52062 | 3.12E-14 | 1.70E-12 | prostaglandin E receptor 3 |
| NPIPB5 | -1.52208 | 3.39E-08 | 3.09E-07 | nuclear pore complex interacting protein family member B3 |
| ADAMTS17 | -1.52315 | 7.45E-14 | 3.57E-12 | ADAM metallopeptidase with thrombospondin type 1 motif 17 |
| KIRREL2 | -1.52639 | 9.09E-07 | 5.65E-06 | kirre like nephrin family adhesion molecule 2 |
| KCNN2 | -1.53517 | 1.74E-05 | 7.62E-05 | potassium calcium-activated channel subfamily N member 2 |
| ZCCHC16 | -1.5378 | 1.81E-07 | 1.36E-06 | retrotransposon Gag like 4 |
| PCDH15 | -1.54021 | 5.79E-05 | 0.000218 | protocadherin related 15 |
| IHH | -1.54027 | 6.35E-14 | 3.15E-12 | Indian hedgehog signaling molecule |
| MIR324 | -1.54166 | 3.55E-09 | 4.27E-08 | microRNA 324 |
| POM121L1P | -1.54329 | 0.000268 | 0.000847 | POM121 transmembrane nucleoporin like 1, pseudogene |
| CTD_3080P12.3 | -1.5469 | 2.56E-05 | 0.000107 | NA |
| GPR26 | -1.54735 | 1.33E-09 | 1.79E-08 | lysophosphatidic acid receptor 1 |
| SLC26A4 | -1.54921 | 7.29E-05 | 0.000267 | solute carrier family 26 member 4 |
| NELL1 | -1.55307 | 1.24E-06 | 7.40E-06 | neural EGFL like 1 |
| MIR27A | -1.55581 | 0.001725 | 0.004348 | microRNA 27a |
| C1QL1 | -1.55627 | 5.31E-05 | 0.000203 | complement C1q like 1 |
| LOC100507537 | -1.55686 | 0.008722 | 0.018254 | NA |
| GSTA7P | -1.5595 | 0.003103 | 0.007337 | glutathione S-transferase alpha 7, pseudogene |
| F2RL3 | -1.56571 | 7.07E-06 | 3.43E-05 | F2R like thrombin or trypsin receptor 3 |
| SLC13A5 | -1.5672 | 0.000283 | 0.000888 | solute carrier family 13 member 5 |
| LINC00482 | -1.568 | 5.39E-06 | 2.70E-05 | long intergenic non-protein coding RNA 482 |
| OR2T10 | -1.56814 | 0.000909 | 0.002489 | olfactory receptor family 2 subfamily T member 10 |
| LINC00853 | -1.57085 | 3.52E-05 | 0.000141 | long intergenic non-protein coding RNA 853 |
| SYT10 | -1.57653 | 2.74E-09 | 3.40E-08 | synaptotagmin 10 |
| PLEKHD1 | -1.58336 | 6.43E-05 | 0.00024 | pleckstrin homology and coiled-coil domain containing D1 |
| ATOH7 | -1.59048 | 0.000221 | 0.000711 | atonal bHLH transcription factor 7 |
| DLGAP2 | -1.5939 | 6.21E-05 | 0.000232 | DLG associated protein 2 |

| | | | | |
|---|---|---|---|---|
| GPAT3 | -1.59835 | 5.49E-09 | 6.21E-08 | glycerol-3-phosphate acyltransferase 3 |
| RHCG | -1.59935 | 5.49E-06 | 2.74E-05 | Rh family C glycoprotein |
| AFM | -1.60342 | 0.000524 | 0.001522 | afamin |
| OR2T35 | -1.60378 | 0.000522 | 0.001518 | olfactory receptor family 2 subfamily T member 35 |
| DRAIC | -1.6082 | 1.00E-05 | 4.67E-05 | downregulated RNA in cancer, inhibitor of cell invasion and migration |
| LMX1B | -1.60974 | 1.24E-07 | 9.72E-07 | LIM homeobox transcription factor 1 beta |
| FCN3 | -1.61235 | 2.85E-05 | 0.000117 | ficolin 3 |
| LOC105375787 | -1.61328 | 1.63E-15 | 1.31E-13 | uncharacterized LOC105375787 |
| TEX41 | -1.61659 | 4.04E-18 | 8.58E-16 | testis expressed 41 |
| CCDC129 | -1.6197 | 2.66E-10 | 4.44E-09 | ITPR interacting domain containing 1 |
| SNORD97 | -1.62035 | 4.58E-16 | 4.50E-14 | small nucleolar RNA, C/D box 97 |
| DEPDC7 | -1.62119 | 2.40E-13 | 9.73E-12 | DEP domain containing 7 |
| PROZ | -1.62175 | 4.91E-06 | 2.49E-05 | protein Z, vitamin K dependent plasma glycoprotein |
| LOC102723886 | -1.63383 | 2.95E-06 | 1.58E-05 | NA |
| CTSV | -1.63517 | 5.08E-06 | 2.56E-05 | cathepsin V |
| SALL3 | -1.63929 | 9.99E-06 | 4.67E-05 | spalt like transcription factor 3 |
| CYP2C8 | -1.63973 | 8.78E-10 | 1.26E-08 | cytochrome P450 family 2 subfamily C member 8 |
| PCK1 | -1.64076 | 2.21E-07 | 1.62E-06 | phosphoenolpyruvate carboxykinase 1 |
| PM20D1 | -1.64521 | 8.01E-06 | 3.83E-05 | peptidase M20 domain containing 1 |
| SLC10A5 | -1.64849 | 7.90E-09 | 8.51E-08 | solute carrier family 10 member 5 |
| CCL3 | -1.65275 | 6.30E-05 | 0.000235 | C-C motif chemokine ligand 3 |
| APOLD1 | -1.65398 | 2.15E-07 | 1.58E-06 | apolipoprotein L domain containing 1 |
| APOC3 | -1.65399 | 0.000578 | 0.001668 | apolipoprotein C3 |
| CXCL2 | -1.65585 | 1.00E-05 | 4.67E-05 | C-X-C motif chemokine ligand 2 |
| MUC13 | -1.65654 | 0.004763 | 0.010683 | mucin 13, cell surface associated |
| OR2T2 | -1.65714 | 3.53E-05 | 0.000142 | olfactory receptor family 2 subfamily T member 2 |
| SERPINA4 | -1.6588 | 1.84E-11 | 4.26E-10 | serpin family A member 4 |
| HPGD | -1.66307 | 3.72E-09 | 4.43E-08 | 15-hydroxyprostaglandin dehydrogenase |
| GSTA2 | -1.66327 | 0.000685 | 0.001942 | glutathione S-transferase alpha 2 |

| | | | | |
|---|---|---|---|---|
| MRPL23_AS1 | -1.66382 | 1.87E-08 | 1.83E-07 | NA |
| LINC00417 | -1.66627 | 0.003022 | 0.007164 | long intergenic non-protein coding RNA 417 |
| PYY | -1.66993 | 1.72E-09 | 2.25E-08 | peptide YY |
| LINC00484 | -1.67276 | 2.60E-13 | 1.04E-11 | long intergenic non-protein coding RNA 484 |
| TMEM207 | -1.67687 | 4.95E-06 | 2.50E-05 | transmembrane protein 207 |
| SLC6A17 | -1.68067 | 4.09E-11 | 8.58E-10 | solute carrier family 6 member 17 |
| LOC101927472 | -1.69372 | 1.33E-11 | 3.18E-10 | NA |
| MIR3183 | -1.69583 | 2.58E-06 | 1.41E-05 | microRNA 3183 |
| GPR3 | -1.69672 | 2.90E-06 | 1.56E-05 | G protein-coupled receptor 3 |
| LOC100129316 | -1.6999 | 2.64E-11 | 5.82E-10 | uncharacterized LOC100129316 |
| ESM1 | -1.70092 | 4.57E-07 | 3.06E-06 | endothelial cell specific molecule 1 |
| NPTX1 | -1.70657 | 1.24E-06 | 7.40E-06 | neuronal pentraxin 1 |
| EGR2 | -1.70759 | 2.52E-05 | 0.000106 | early growth response 2 |
| NPHS1 | -1.70946 | 4.49E-05 | 0.000175 | NPHS1 adhesion molecule, nephrin |
| TMEM200C | -1.71123 | 4.07E-11 | 8.55E-10 | transmembrane protein 200C |
| SLC12A3 | -1.71206 | 0.000301 | 0.000939 | solute carrier family 12 member 3 |
| LOC101926962 | -1.71466 | 5.85E-14 | 2.93E-12 | NA |
| DUSP2 | -1.71475 | 5.14E-09 | 5.86E-08 | dual specificity phosphatase 2 |
| LINC01108 | -1.71621 | 3.11E-10 | 5.08E-09 | long intergenic non-protein coding RNA 1108 |
| SNTG1 | -1.72162 | 6.01E-05 | 0.000226 | syntrophin gamma 1 |
| JUNB | -1.72743 | 1.27E-11 | 3.04E-10 | JunB proto-oncogene, AP-1 transcription factor subunit |
| C2orf54 | -1.72965 | 0.002846 | 0.006782 | mab-21 like 4 |
| LOC643623 | -1.73221 | 9.09E-06 | 4.28E-05 | NA |
| HCRTR2 | -1.73308 | 0.005176 | 0.011516 | hypocretin receptor 2 |
| GJA3 | -1.73464 | 1.14E-07 | 9.07E-07 | gap junction protein alpha 3 |
| TINCR | -1.73716 | 4.66E-09 | 5.37E-08 | TINCR ubiquitin domain containing |
| BRE_AS1 | -1.74006 | 3.05E-11 | 6.58E-10 | NA |
| MIR554 | -1.74266 | 1.21E-11 | 2.93E-10 | microRNA 554 |
| DPP6 | -1.74369 | 1.70E-11 | 3.97E-10 | dipeptidyl peptidase like 6 |
| LRRC9 | -1.74609 | 7.61E-08 | 6.35E-07 | leucine rich repeat containing 9 |
| CYP3A7 | -1.74669 | 3.40E-06 | 1.79E-05 | cytochrome P450 family 3 subfamily A member 7 |
| CRABP1 | -1.75187 | 7.18E-05 | 0.000264 | cellular retinoic acid binding protein 1 |
| JUN | -1.75534 | 2.36E-10 | 4.00E-09 | Jun proto-oncogene, AP-1 transcription factor subunit |
| OR2T3 | -1.75883 | 0.000115 | 0.000398 | olfactory receptor family 2 subfamily T member 3 |

| | | | | |
|---|---|---|---|---|
| FLJ31356 | -1.7639 | 6.97E-15 | 4.66E-13 | uncharacterized protein FLJ31356 |
| ALB | -1.76397 | 0.001426 | 0.00369 | albumin |
| LINC01517 | -1.76412 | 0.002535 | 0.006122 | long intergenic non-protein coding RNA 1517 |
| CR2 | -1.7659 | 0.000903 | 0.002475 | complement C3d receptor 2 |
| OLIG2 | -1.77 | 6.68E-10 | 9.89E-09 | oligodendrocyte transcription factor 2 |
| GADL1 | -1.77334 | 0.001621 | 0.004121 | glutamate decarboxylase like 1 |
| KCNG3 | -1.77653 | 1.21E-15 | 1.02E-13 | potassium voltage-gated channel modifier subfamily G member 3 |
| GPRC6A | -1.77724 | 0.000766 | 0.00214 | G protein-coupled receptor class C group 6 member A |
| ZNF804B | -1.77879 | 5.69E-08 | 4.91E-07 | zinc finger protein 804B |
| LINC01485 | -1.78297 | 2.73E-07 | 1.94E-06 | long intergenic non-protein coding RNA 1485 |
| DDN | -1.78639 | 2.83E-05 | 0.000117 | dendrin |
| CALB1 | -1.78661 | 2.40E-05 | 0.000101 | calbindin 1 |
| CHI3L1 | -1.78697 | 2.79E-08 | 2.61E-07 | chitinase 3 like 1 |
| CBARP | -1.79463 | 7.52E-09 | 8.17E-08 | CACN subunit beta associated regulatory protein |
| LINC00964 | -1.79698 | 6.26E-07 | 4.06E-06 | long intergenic non-protein coding RNA 964 |
| MYCNOS | -1.80655 | 2.97E-05 | 0.000122 | MYCN opposite strand |
| IP6K3 | -1.80821 | 5.90E-09 | 6.61E-08 | inositol hexakisphosphate kinase 3 |
| OR2T34 | -1.80968 | 8.15E-05 | 0.000295 | olfactory receptor family 2 subfamily T member 34 |
| ANKRD20A8P | -1.81648 | 0.002296 | 0.005618 | ankyrin repeat domain 20 family member A8, pseudogene |
| WDR46 | -1.82633 | 0.024713 | 0.044943 | WD repeat domain 46 |
| ETNPPL | -1.82659 | 4.61E-05 | 0.000179 | ethanolamine-phosphate phospho-lyase |
| SLC2A3 | -1.82844 | 2.50E-13 | 1.01E-11 | solute carrier family 2 member 3 |
| IGFBP1 | -1.82895 | 0.000114 | 0.000396 | insulin like growth factor binding protein 1 |
| FAM95A | -1.83636 | 0.008825 | 0.018427 | family with sequence similarity 95 member A |
| MAFF | -1.83794 | 7.24E-08 | 6.07E-07 | MAF bZIP transcription factor F |
| GRIA2 | -1.84316 | 1.67E-06 | 9.59E-06 | glutamate ionotropic receptor AMPA type subunit 2 |
| APOH | -1.84324 | 3.43E-05 | 0.000138 | apolipoprotein H |
| LOC200772 | -1.84408 | 4.24E-07 | 2.87E-06 | NA |

| | | | | |
|---|---|---|---|---|
| GADD45B | -1.84512 | 4.00E-18 | 8.58E-16 | growth arrest and DNA damage inducible beta |
| CXXC4_AS1 | -1.84588 | 2.13E-05 | 9.10E-05 | NA |
| SYCP2 | -1.86989 | 2.09E-15 | 1.59E-13 | synaptonemal complex protein 2 |
| PCAT29 | -1.87729 | 1.11E-05 | 5.13E-05 | prostate cancer associated transcript 29 |
| MIR23A | -1.87897 | 5.41E-08 | 4.69E-07 | microRNA 23a |
| LINC00473 | -1.8884 | 3.43E-08 | 3.12E-07 | phosphodiesterase 10A |
| GDF15 | -1.90234 | 4.34E-13 | 1.62E-11 | growth differentiation factor 15 |
| TMEM105 | -1.90362 | 0.000163 | 0.000548 | TMEM105 long non-coding RNA |
| LINC00323 | -1.90607 | 8.03E-08 | 6.67E-07 | long intergenic non-protein coding RNA 323 |
| KRTAP5_8 | -1.91461 | 1.74E-09 | 2.28E-08 | NA |
| MIR221 | -1.91685 | 2.96E-06 | 1.59E-05 | microRNA 221 |
| SLCO4A1_AS1 | -1.92649 | 2.93E-06 | 1.57E-05 | NA |
| ADAMTS4 | -1.93418 | 8.52E-05 | 0.000307 | ADAM metallopeptidase with thrombospondin type 1 motif 4 |
| IL6 | -1.9346 | 0.001358 | 0.003537 | interleukin 6 |
| C11orf87 | -1.943 | 0.001174 | 0.003114 | chromosome 11 open reading frame 87 |
| MRO | -1.95092 | 1.15E-08 | 1.20E-07 | maestro |
| ERRFI1 | -1.95966 | 3.66E-10 | 5.88E-09 | ERBB receptor feedback inhibitor 1 |
| TRIM72 | -1.96113 | 1.10E-11 | 2.71E-10 | tripartite motif containing 72 |
| PTHLH | -1.98708 | 4.46E-09 | 5.17E-08 | parathyroid hormone like hormone |
| HRG | -1.99531 | 1.97E-09 | 2.54E-08 | neuregulin 1 |
| LOC101928596 | -1.99544 | 5.09E-06 | 2.57E-05 | uncharacterized LOC101928596 |
| LINC01847 | -2.00345 | 2.90E-06 | 1.57E-05 | long intergenic non-protein coding RNA 1847 |
| CEL | -2.01976 | 5.37E-09 | 6.08E-08 | carboxyl ester lipase |
| FOXF1 | -2.04346 | 9.88E-17 | 1.22E-14 | forkhead box F1 |
| HELT | -2.04429 | 9.83E-07 | 6.04E-06 | helt bHLH transcription factor |
| G6PC | -2.05276 | 2.11E-08 | 2.03E-07 | glucose-6-phosphatase catalytic subunit |
| ASB15 | -2.05455 | 2.33E-06 | 1.29E-05 | ankyrin repeat and SOCS box containing 15 |
| NR0B2 | -2.05525 | 1.62E-17 | 2.65E-15 | nuclear receptor subfamily 0 group B member 2 |
| TCF24 | -2.05996 | 2.92E-08 | 2.70E-07 | transcription factor 24 |
| RGS1 | -2.06655 | 1.75E-05 | 7.65E-05 | regulator of G protein signaling 1 |
| CCDC144A | -2.06893 | 5.33E-06 | 2.67E-05 | coiled-coil domain containing 144A |
| GAD1 | -2.08069 | 7.35E-12 | 1.92E-10 | glutamate decarboxylase 1 |

| | | | | |
|---|---|---|---|---|
| HMGN5 | -2.08579 | 7.48E-08 | 6.26E-07 | high mobility group nucleosome binding domain 5 |
| ARC | -2.08858 | 1.20E-08 | 1.24E-07 | nucleolar protein 3 |
| CYR61 | -2.08961 | 4.29E-19 | 1.21E-16 | cellular communication network factor 1 |
| NDNF | -2.0949 | 8.14E-07 | 5.13E-06 | neuron derived neurotrophic factor |
| EGF | -2.11612 | 1.33E-10 | 2.40E-09 | epidermal growth factor |
| FMN2 | -2.11746 | 3.49E-09 | 4.20E-08 | formin 2 |
| IGF2 | -2.12528 | 3.27E-06 | 1.73E-05 | insulin like growth factor 2 |
| PDK4 | -2.1272 | 2.19E-13 | 9.01E-12 | pyruvate dehydrogenase kinase 4 |
| MIR6863 | -2.13374 | 7.61E-06 | 3.66E-05 | microRNA 6863 |
| CDH15 | -2.18522 | 8.71E-05 | 0.000313 | cadherin 15 |
| LINC01230 | -2.19425 | 1.72E-10 | 3.02E-09 | long intergenic non-protein coding RNA 1230 |
| CYP27B1 | -2.23793 | 7.17E-11 | 1.41E-09 | cytochrome P450 family 27 subfamily B member 1 |
| TNS4 | -2.28099 | 8.69E-11 | 1.66E-09 | tensin 4 |
| TRPM6 | -2.28518 | 5.63E-07 | 3.70E-06 | transient receptor potential cation channel subfamily M member 6 |
| ZFP36 | -2.29673 | 2.97E-19 | 8.78E-17 | ZFP36 ring finger protein |
| RNF17 | -2.30855 | 0.003811 | 0.008769 | ring finger protein 17 |
| LOC221946 | -2.33194 | 6.59E-13 | 2.35E-11 | uncharacterized LOC221946 |
| CYP4Z1 | -2.34652 | 8.95E-10 | 1.28E-08 | cytochrome P450 family 4 subfamily Z member 1 |
| FER1L6_AS1 | -2.37413 | 3.92E-07 | 2.68E-06 | NA |
| TRIM50 | -2.39264 | 2.86E-07 | 2.02E-06 | tripartite motif containing 50 |
| AKR1B10 | -2.39926 | 9.19E-06 | 4.33E-05 | aldo-keto reductase family 1 member B10 |
| CYP26B1 | -2.40955 | 6.83E-17 | 8.89E-15 | cytochrome P450 family 26 subfamily B member 1 |
| GUCY2EP | -2.4227 | 1.05E-07 | 8.44E-07 | guanylate cyclase 2E, pseudogene |
| MIR6723 | -2.42581 | 7.67E-14 | 3.65E-12 | NA |
| SPDYE7P | -2.44845 | 0.000144 | 0.00049 | speedy/RINGO cell cycle regulator family member E7, pseudogene |
| LOC101927136 | -2.46565 | 1.75E-06 | 1.00E-05 | uncharacterized LOC101927136 |
| HSPA1B | -2.48085 | 4.28E-08 | 3.82E-07 | heat shock protein family A (Hsp70) member 1B |
| MIR6883 | -2.48189 | 7.11E-12 | 1.86E-10 | microRNA 6883 |
| GC | -2.49748 | 0.000227 | 0.00073 | glucosylceramidase beta pseudogene 1 |
| FER1L6_AS2 | -2.50023 | 2.67E-06 | 1.45E-05 | NA |

| | | | | |
|---|---|---|---|---|
| HSPA1A | -2.51866 | 0.024475 | 0.044566 | heat shock protein family A (Hsp70) member 1A |
| MIR5690 | -2.64233 | 1.19E-06 | 7.12E-06 | microRNA 5690 |
| EGR3 | -2.69662 | 2.71E-12 | 8.08E-11 | early growth response 3 |
| KLK1 | -2.75454 | 2.42E-05 | 0.000102 | kallikrein 1 |
| USP32P1 | -2.79352 | 6.57E-05 | 0.000244 | ubiquitin specific peptidase 32 pseudogene 1 |
| DUSP1 | -2.94099 | 9.14E-27 | 1.61E-23 | dual specificity phosphatase 1 |
| SRRM4 | -2.95652 | 3.11E-07 | 2.18E-06 | serine/arginine repetitive matrix 4 |
| MIR3189 | -3.31823 | 3.64E-20 | 1.44E-17 | microRNA 3189 |
| RNR2 | -3.32325 | 4.57E-22 | 2.80E-19 | l-rRNA |
| FOS | -3.36593 | 9.97E-09 | 1.05E-07 | Fos proto-oncogene, AP-1 transcription factor subunit |
| ATF3 | -3.43144 | 2.93E-36 | 1.26E-32 | activating transcription factor 3 |
| NR4A3 | -3.47933 | 1.26E-26 | 1.80E-23 | nuclear receptor subfamily 4 group A member 3 |
| EGR1 | -3.47947 | 2.06E-17 | 3.20E-15 | early growth response 1 |
| RNR1 | -3.49082 | 6.68E-18 | 1.31E-15 | s-rRNA |
| NR4A2 | -4.02834 | 1.22E-44 | 1.05E-40 | nuclear receptor subfamily 4 group A member 2 |
| NR4A1 | -4.67303 | 2.39E-56 | 4.11E-52 | nuclear receptor subfamily 4 group A member 1 |
| FOSB | -6.46167 | 1.58E-39 | 9.05E-36 | FosB proto-oncogene, AP-1 transcription factor subunit |

**Table S3**: Differentially expressed genes from Cohort B: DN. A total of 80 dysregulations were observed – 46 upregulations and 34 downregulations.

| **Gene Symbol** | **$log_2$FC** | ***P*-Value** | ***Q*-Value** | **Gene Name** |
|---|---|---|---|---|
| HBM | 2.854358 | 1.20E-05 | 0.0017 | Hemoglobin Subunit Mu |
| C15orf48 | 2.736677 | 8.34E-06 | 0.0013 | Normal Mucosa Of Esophagus-Specific Gene 1 Protein |
| HLA_DRB6 | 2.637187 | 0.001057 | 0.034054 | Major Histocompatibility Complex, Class II, DR Beta 6 (Pseudogene) |
| ALAS2 | 2.586015 | 5.41E-07 | 0.000267 | 5'-Aminolevulinate Synthase 2 |

| | | | | |
|---|---|---|---|---|
| OLR1 | 2.580787 | 0.000191 | 0.013882 | Oxidized Low Density Lipoprotein Receptor 1 |
| A_32_P47166 | 2.42549 | 8.57E-05 | 0.013882 | Long noncoding RNA |
| CXCL3 | 2.415666 | 1.08E-07 | 0.000107 | C-X-C Motif Chemokine Ligand 3 |
| EPB42 | 2.37307 | 0.000289 | 0.01503 | Erythrocyte Membrane Protein Band 4.2 |
| A_32_P4882 | 2.279254 | 5.56E-05 | 0.005885 | Long noncoding RNA |
| SERPINB2 | 2.176467 | 0.000174 | 0.013207 | SERPINB2 |
| IL1B | 2.084519 | 4.79E-07 | 0.000267 | Interleukin 1 Beta |
| EGR4 | 2.076215 | 2.01E-05 | 0.002486 | Early Growth Response 4 |
| CXCL2 | 2.075299 | 1.55E-09 | 2.30E-06 | C-X-C Motif Chemokine Ligand 2 |
| HBD | 2.042676 | 3.16E-06 | 0.00064 | Hemoglobin Subunit Delta |
| CA1 | 2.039417 | 0.000391 | 0.018383 | Carbonic Anhydrase 1 |
| THBD | 1.845336 | 4.36E-06 | 0.000735 | Thrombomodulin |
| EREG | 1.812352 | 0.00022 | 0.013894 | Epiregulin |
| CXCL1 | 1.763524 | 9.42E-05 | 0.008647 | C-X-C Motif Chemokine Ligand 1 |
| RGS1 | 1.727592 | 2.39E-07 | 0.000177 | Regulator Of G Protein Signaling 1 |
| HBA2 | 1.575436 | 0.000134 | 0.011353 | Hemoglobin Subunit Alpha 2 |
| G0S2 | 1.566598 | 1.73E-05 | 0.002336 | G0/G1 Switch 2 |
| ZNF331 | 1.555593 | 3.81E-05 | 0.004251 | Zinc Finger Protein 331 |
| IL8 | 1.515433 | 9.25E-06 | 0.00137 | C-X-C Motif Chemokine Ligand 8 |
| A_32_P213948 | 1.508937 | 7.81E-05 | 0.007714 | Long noncoding RNA |
| A_32_P132393 | 1.479313 | 0.001428 | 0.03883 | Long noncoding RNA |

| | | | | |
|---|---|---|---|---|
| A_32_P74243 | 1.389347 | 2.72E-05 | 0.003226 | Long noncoding RNA |
| A_24_P185516 | 1.382731 | 3.56E-06 | 0.000659 | Long noncoding RNA |
| NR4A3 | 1.373492 | 0.00026 | 0.015019 | Nuclear Receptor Subfamily 4 Group A Member 3 |
| FAM115C | 1.333011 | 4.46E-06 | 0.000735 | TRPM8 Channel Associated Factor 2 |
| GABARAPL1 | 1.327218 | 0.001977 | 0.047543 | GABA Type A Receptor Associated Protein Like 1 |
| IL10 | 1.306255 | 1.93E-05 | 0.002486 | Interleukin 10 |
| ALOX12 | 1.225999 | 0.000194 | 0.013882 | Arachidonate 12-Lipoxygenase, 12S Type |
| PLAUR | 1.2234 | 0.000233 | 0.014409 | Plasminogen Activator, Urokinase Receptor |
| ENST00000437044 | 1.182648 | 0.000286 | 0.01503 | Triggering Receptor Expressed On Myeloid Cells Like 1 |
| GRASP | 1.180337 | 0.000214 | 0.013882 | Trafficking Regulator And Scaffold Protein Tamalin |
| PTGS2 | 1.156994 | 0.000559 | 0.023521 | Prostaglandin-Endoperoxide Synthase 2 |
| KCNJ15 | 1.141434 | 0.000264 | 0.015019 | Potassium Inwardly Rectifying Channel Subfamily J Member 15 |
| ENST00000419533 | 1.104823 | 0.001151 | 0.035174 | Ubiquitin Protein Ligase E3 Component N-Recognin 4 |
| JMY | 1.103278 | 0.000961 | 0.032347 | Junction Mediating And Regulatory |

| | | | | |
|---|---|---|---|---|
| | | | | Protein, P53 Cofactor |
| SAMSN1 | 1.099563 | 0.000118 | 0.010274 | SAM Domain, SH3 Domain And Nuclear Localization Signals 1 |
| ZNF571 | 1.091636 | 0.000733 | 0.027181 | Zinc Finger Protein 571 |
| CD365380 | 1.050094 | 0.000535 | 0.022966 | NA |
| PDE4D | 1.045838 | 0.000715 | 0.027181 | Phosphodiesterase 4D |
| A_24_P746044 | 1.036553 | 0.001451 | 0.038996 | Long noncoding RNA |
| SLC16A6 | 1.013251 | 0.000272 | 0.015019 | Solute Carrier Family 16 Member 6 |
| TRIB1 | 1.003009 | 0.000859 | 0.030302 | Tribbles Pseudokinase 1 |
| GCET2 | -1.00926 | 0.000601 | 0.024386 | Germinal Center Associated Signaling And Motility |
| AF150244 | -1.0194 | 0.000442 | 0.020461 | cDNA clone CBFBBE12 |
| ENST00000453166 | -1.02295 | 0.000833 | 0.029725 | Immunoglobulin Kappa Variable 2D-28 |
| ENST00000443866 | -1.03499 | 0.0002 | 0.013882 | Ciliary Rootlet Coiled-Coil, Rootletin Family Member 2 |
| A_32_P223985 | -1.03804 | 0.000725 | 0.027181 | Long noncoding RNA |
| A_24_P255384 | -1.07271 | 0.000786 | 0.028419 | Long noncoding RNA |
| ENST00000390353 | -1.07638 | 0.000981 | 0.032644 | T Cell Receptor Beta Variable 6-1 |
| A_32_P926336 | -1.07644 | 0.001729 | 0.043423 | Long noncoding RNA |
| A_32_P119604 | -1.09647 | 0.000734 | 0.027181 | Long noncoding RNA |
| A_24_P379629 | -1.0975 | 0.001122 | 0.034626 | Long noncoding RNA |
| A_23_P170830 | -1.09754 | 0.001261 | 0.036981 | Long noncoding RNA |

| | | | | |
|---|---|---|---|---|
| KLHL34 | -1.11447 | 0.00111 | 0.034626 | Kelch Like Family Member 34 |
| GIMAP8 | -1.11678 | 0.000171 | 0.013207 | GTPase, IMAP Family Member 8 |
| ENST00000390618 | -1.12422 | 0.000902 | 0.030771 | Immunoglobulin Heavy Variable 3-38 (Non-Functional) |
| A_32_P65804 | -1.15271 | 0.000216 | 0.013882 | Long noncoding RNA |
| A_24_P752999 | -1.16719 | 9.63E-05 | 0.008647 | Long noncoding RNA |
| VSIG1 | -1.18924 | 9.50E-07 | 0.000313 | V-Set And Immunoglobulin Domain Containing 1 |
| A_32_P67223 | -1.21181 | 0.001117 | 0.034626 | Long noncoding RNA |
| IL12RB2 | -1.23068 | 0.000276 | 0.015019 | Interleukin 12 Receptor Subunit Beta 2 |
| HAL | -1.23239 | 3.24E-06 | 0.00064 | Histidine Ammonia-Lyase |
| A_32_P175042 | -1.24197 | 0.000903 | 0.030771 | Long noncoding RNA |
| RFPL2 | -1.27583 | 0.000533 | 0.022966 | Ret Finger Protein Like 2 |
| A_32_P121140 | -1.27984 | 0.000246 | 0.01488 | Long noncoding RNA |
| LOC641518 | -1.32182 | 0.001312 | 0.037754 | Long-noncoding RNA 641518 |
| GPR34 | -1.35048 | 0.000321 | 0.016112 | G Protein-Coupled Receptor 34 |
| C2orf40 | -1.35273 | 2.36E-06 | 0.000636 | ECRG4 Augurin Precursor |
| ENST00000390625 | -1.51018 | 0.001277 | 0.037093 | Immunoglobulin Heavy Variable 3-49 |
| A_24_P868313 | -1.53375 | 0.000305 | 0.01559 | Long noncoding RNA |
| ENST00000474213 | -1.53822 | 8.11E-07 | 0.0003 | Immunoglobulin Kappa Variable 2D-30 |
| A_32_P90346 | -1.56933 | 3.09E-06 | 0.00064 | Long noncoding RNA |
| UTS2 | -1.63242 | 0.000164 | 0.013162 | Urotensin 2 |

| | | | | |
|---|---|---|---|---|
| RP11_327P2.4 | -1.65529 | 1.78E-06 | 0.000527 | Clone RP11-327P2 |
| SLC4A10 | -1.66138 | 3.11E-06 | 0.00064 | Solute Carrier Family 4 Member 10 |
| LRRN3 | -2.53057 | 4.74E-11 | 1.40E-07 | Leucine-Rich Repeat Neuronal Protein 3 |
